# The Energetics of Molecular Adaptation in Transcriptional Regulation

**DOI:** 10.1101/638270

**Authors:** Griffin Chure, Manuel Razo-Mejia, Nathan M. Belliveau, Tal Einav, Zofii A. Kaczmarek, Stephanie L. Barnes, Mitchell Lewis, Rob Phillips

## Abstract

Mutation is a critical mechanism by which evolution explores the functional landscape of proteins. Despite our ability to experimentally inflict mutations at will, it remains difficult to link sequence-level perturbations to systems-level responses. Here, we present a framework centered on measuring changes in the free energy of the system to link individual mutations in an allosteric transcriptional repressor to the parameters which govern its response. We find the energetic effects of the mutations can be categorized into several classes which have characteristic curves as a function of the inducer concentration. We experimentally test these diagnostic predictions using the well-characterized LacI repressor of *Escherichia coli*, probing several mutations in the DNA binding and inducer binding domains. We find that the change in gene expression due to a point mutation can be captured by modifying only a subset of the model parameters that describe the respective domain of the wild-type protein. These parameters appear to be insulated, with mutations in the DNA binding domain altering only the DNA affinity and those in the inducer binding domain altering only the allosteric parameters. Changing these subsets of parameters tunes the free energy of the system in a way that is concordant with theoretical expectations. Finally, we show that the induction profiles and resulting free energies associated with pairwise double mutants can be predicted with quantitative accuracy given knowledge of the single mutants, providing an avenue for identifying and quantifying epistatic interactions.

**Summary:** We present a biophysical model of allosteric transcriptional regulation that directly links the location of a mutation within a repressor to the biophysical parameters that describe its behavior. We explore the phenotypic space of a repressor with mutations in either the inducer binding or DNA binding domains. Using the LacI repressor in *E. coli*, we make sharp, falsifiable predictions and use this framework to generate a null hypothesis for how double mutants behave given knowledge of the single mutants. Linking mutations to the parameters which govern the system allows for quantitative predictions of how the free energy of the system changes as a result, permitting coarse graining of high-dimensional data into a single-parameter description of the mutational consequences.

---

Thermodynamic treatments of transcriptional regulation have been fruitful in their ability to generate quantitative predictions of gene expression as a function of a minimal set of physically meaningful variables (1–13). These models quantitatively describe numerous properties of input-output functions, such as the leakiness, saturation, dynamic range, steepness of response, and the [*EC*_50_] – the concentration of inducer at which the response is half maximal. The mathematical forms of these phenotypic properties are couched in terms of a minimal set of experimentally accessible variables, such as the inducer concentration, transcription factor copy number, and the DNA sequence of the binding site (10). While the amino acid sequence of the transcription factor is another controllable variable, it is seldom implemented in quantitative terms considering mutations with subtle changes in chemistry frequently result in unpredictable physiological consequences. In this work, we examine how a series of mutations in either the DNA binding or inducer binding domains of a transcriptional repressor influence the values of the biophysical parameters which govern its regulatory behavior.

We first present a theoretical framework for understanding how mutations in the repressor affect different parameters and alter the free energy of the system. The multi-dimensional parameter space of the aforementioned thermodynamic models is highly degenerate with multiple combinations of parameter values yielding the same phenotypic response. This degeneracy can be subsumed into the free energy of the system, transforming the input-output function into a one-dimensional description with the form of a Fermi function (14, 15). We find that the parameters capturing the allosteric nature of the repressor, the repressor copy number, and the DNA binding specificity contribute independently to the free energy of the system with different degrees of sensitivity. Furthermore, changes restricted to one of these three groups of parameters result in characteristic changes in the free energy relative to the wild-type repressor, providing falsifiable predictions of how different classes of mutations should behave.

Next, we test these descriptions experimentally using the well-characterized transcriptional repressor of the *lac* operon LacI in *E. coli* regulating expression of a fluorescent reporter. We introduce a series of point mutations in either the inducer binding or DNA binding domain. We then measure the full induction profile of each mutant, determine the minimal set of parameters that are affected by the mutation, and predict how each mutation tunes the free energy at different inducer concentrations, repressor copy numbers, and DNA binding strengths. We find in general that mutations in the DNA binding domain only influence DNA binding strength, and that mutations within the inducer binding domain affect only the parameters which dictate the allosteric response. The degree to which these parameters are insulated is notable, as the very nature of allostery suggests that all parameters are intimately connected, thus enabling binding events at one domain to be “sensed” by another.

With knowledge of how a collection of DNA binding and inducer binding single mutants behave, we predict the induction profiles and the free energy changes of pairwise double mutants with quantitative accuracy. We find that the energetic effects of each individual mutation are additive, indicating that epistatic interactions are absent between the mutations examined here. Our model provides a means for identifying and quantifying the extent of epistatic interactions in a more complex set of mutations, and can shed light on how the protein sequence and general regulatory architecture coevolve.

## Results

This work considers the inducible simple repression regulatory motif [depicted in Fig. 1(A)] from a thermodynamic perspective which has been thoroughly dissected and tested experimentally (4, 6, 10). While we direct the reader to the SI text for a complete derivation, the result of this extensive theory-experiment dialogue is a succinct input-output function [schematized in Fig. 1(B)] that computes the fold-change in gene expression relative to an unregulated promoter. This function is of the form

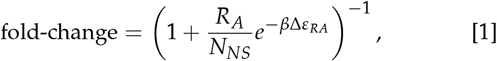

where *R*_*A*_ is the number of active repressors per cell, *N*_*NS*_ is the number of non-specific binding sites for the repressor, ∆*ε*_*RA*_ is the binding energy of the repressor to its specific binding site relative to the non-specific background, and *β* is defined as 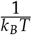 where *k*_*B*_ is the Boltzmann constant and *T* is the temperature. While this theory requires knowledge of the number of *active* repressors, we often only know the total number *R* which is the sum total of active and inactive repressors. We can define a prefactor *p*_act_(*c*) which captures the allosteric nature of the repressor and encodes the probability a repressor is in the active (repressive) state rather than the inactive state for a given inducer concentration *c*, namely,

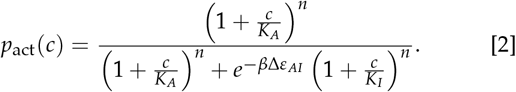

**Fig. 1.**
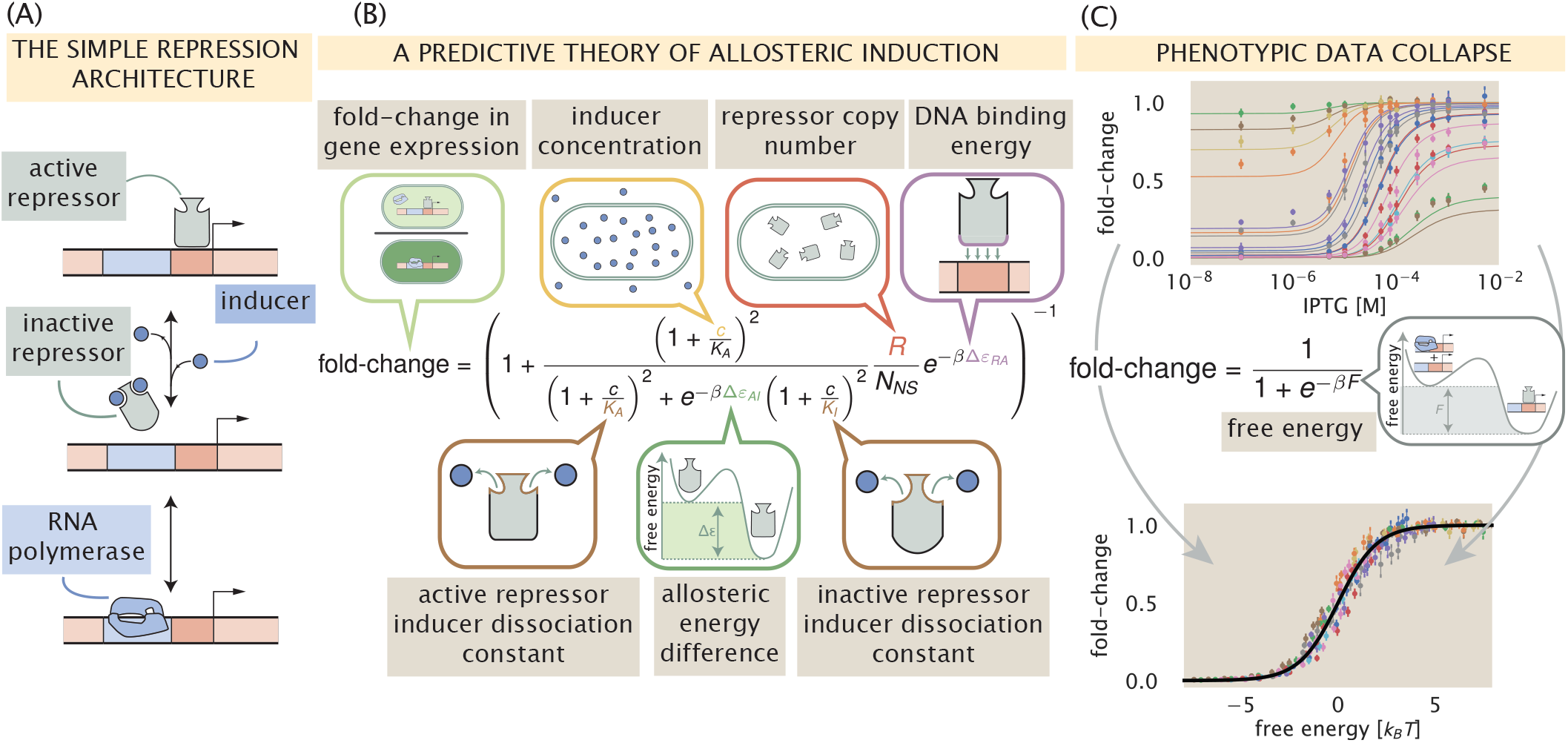
A predictive framework for phenotypic and energetic dissection of the simple repression motif. (A) The inducible simple repression architecture. When in the active state, the repressor (gray) binds the cognate operator sequence of the DNA (red box) with high specificity, preventing transcription by occluding binding of the RNA polymerase to the promoter (blue rectangle). Upon addition of an inducer molecule, the inactive state becomes energetically preferable and the repressor no longer binds the operator sequence with appreciable specificity. Once unbound from the operator, binding of the RNA polymerase (blue) is no longer blocked and transcription can occur. (B) The simple repression input-output function for an allosteric repressor with two inducer binding sites. The key parameters are identified in speech bubbles. (C) Fold-change in gene expression collapses as a function of the free energy. The input-output function in (B) can be re-written as a Fermi function with an energetic parameter *F* which is the energetic difference between the repressor bound and unbound states of the promoter. Top panel shows induction profiles reported in Razo-Mejia *et al.* 2018 (10) of eighteen different strains over twelve concentrations of the gratuitous inducer Isopropyl *β*-D-1-thiogalactopyranoside (IPTG). Upon calculation of the free energy, the data collapse onto a single master curve (bottom panel) defined by *F*.

Here, *K*_*A*_ and *K*_*I*_ are the dissociation constants of the inducer to the active and inactive repressor, ∆*ε*_*AI*_ is the energetic difference between the repressor active and inactive states, and *n* is the number of allosteric binding sites per repressor molecule (*n* = 2 for LacI). With this in hand, we can define *R*_*A*_ in Eq. (1) as *R*_*A*_ = *p*_act_(*c*)*R*.

A key feature of Eq. (1) and Eq. (2) is that the diverse phenomenology of the gene expression induction profile can be collapsed onto a single master curve by rewriting the input-output function in terms of the free energy *F* [also called the Bohr parameter (16)],

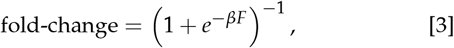

where

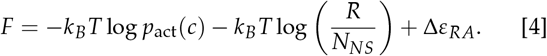

Hence, if different combinations of parameters yield the same free energy, they will give rise to the same fold-change in gene expression, enabling us to collapse multiple regulatory scenarios onto a single curve. This can be seen in Fig. 1(C) where eighteen unique inducer titration profiles of a LacI simple repression architecture collected and analyzed in Razo-Mejia *et al.* 2018 (10) collapse onto a single master curve. The tight distribution about this curve reveals that fold-change across a variety of genetically distinct individuals can be adequately described by a small number of parameters. Beyond predicting the induction profiles of different strains, the method of data collapse inspired by Eq. (3) and Eq. (4) can be used as a tool to identify mechanistic changes in the regulatory architecture (14). Similar data collapse approaches have been used previously in such a manner and have proved vital for distinguishing between changes in parameter values and changes in the fundamental behavior of the system (14, 15).

Assuming that a given mutation does not result in a non-functional protein, it is reasonable to say that any or all of the parameters in Eq. (1) can be affected by the mutation, changing the observed induction profile and therefore the free energy. To examine how the free energy of a mutant *F*^(mut)^ differs from that of the wild-type *F*^(wt)^, we define Δ*F* = *F*^(mut)^ − *F*^(wt)^, which has the form

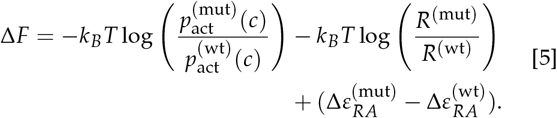

∆*F* describes how a mutation translates a point across the master curve shown in Fig. 1(C). As we will show in the coming paragraphs [illustrated in Fig. 2], this formulation coarse grains the myriad parameters shown in Eq. (1) and Eq. (2) into three distinct quantities, each with different sensitivities to parametric changes. By examining how a mutation changes the free energy changes as a function of the inducer concentration, one can draw conclusions as to which parameters have been modified based solely on the shape of the curve. To help the reader understand how various perturbations to the parameters tune the free energy, we have hosted an interactive figure on the paper website which makes exploration of parameter space a simpler task.

**Fig. 2.**
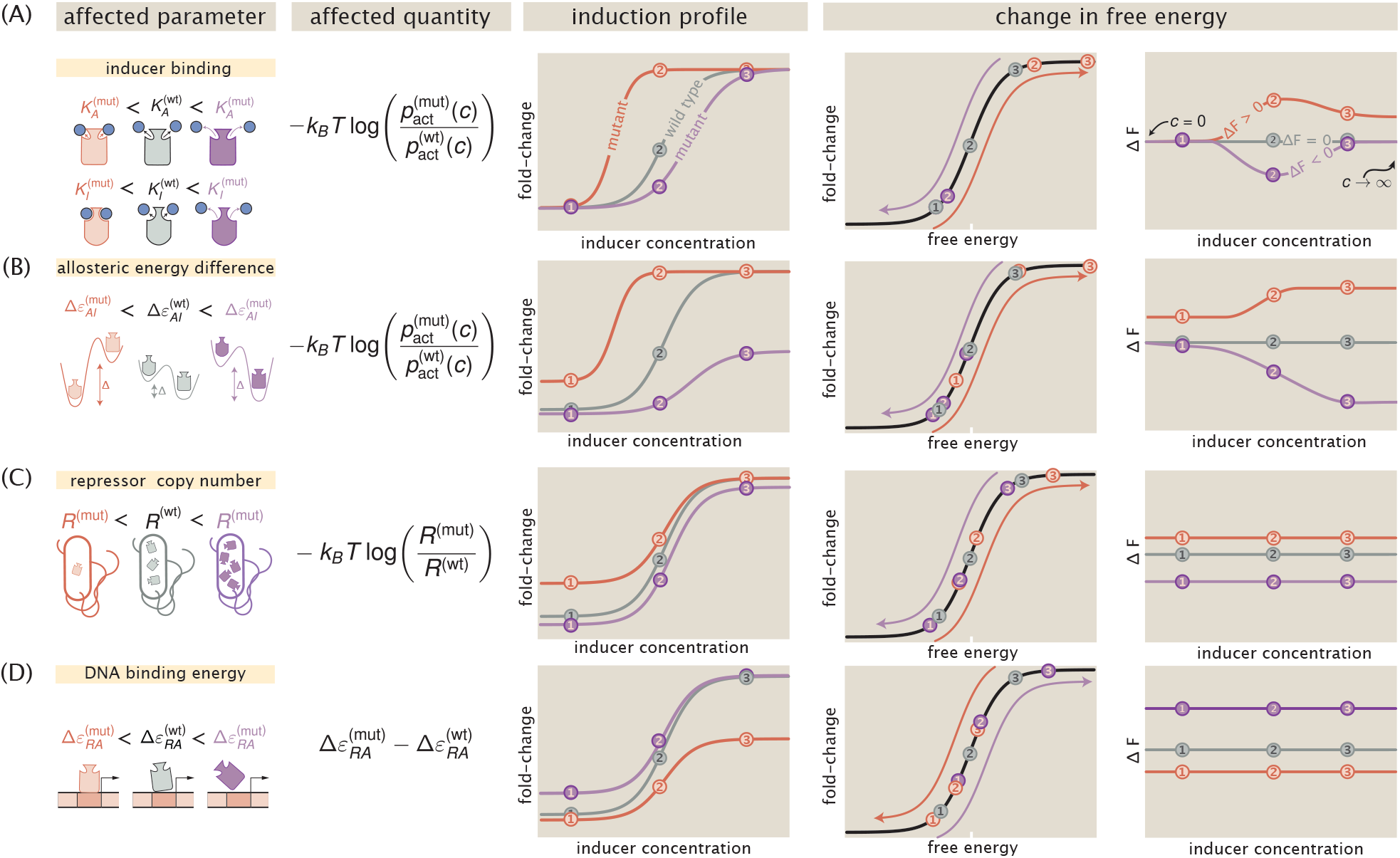
Parametric changes due to mutations alter the free energy. The first column schematizes the changed parameters and the second column reflects which quantity in Eq. (5) is affected. The third column shows representative induction profiles from mutants which have smaller (red) and larger (purple) values for the parameters than the wild-type (grey). The fourth and fifth columns illustrate how the free energy is changed as a result. Purple and red arrows indicate the direction in which the points are translated about the master curve. Three concentrations (points labeled 1, 2, and 3) are shown to illustrate how each point is moved in free energy space.

The first term in Eq. (5) is the log ratio of the probability of a mutant repressor being active relative to the wild type at a given inducer concentration *c*. This quantity defines how changes to any of the allosteric parameters – such as inducer binding constants *K*_*A*_ and *K*_*I*_, or active/inactive state energetic difference ∆*ε*_*AI*_ – alter the free energy *F*, which can be interpreted as the free energy difference between the repressor bound and unbound states of the promoter. Fig. 2 (A) illustrates how changes to the inducer binding constants *K*_*A*_ and *K*_*I*_ alone alter the induction profiles and resulting free energy as a function of the inducer concentration. In the limit where *c* = 0, the values of *K*_*A*_ and *K*_*I*_ do not factor into the calculation of *p*_act_(*c*) given by Eq. (2), meaning that ∆*ε*_*AI*_ is the lone parameter setting the residual activity of the repressor. Thus, if only *K*_*A*_ and *K*_*I*_ are altered by a mutation, then ∆*F* should be 0*k*_*B*_*T* when *c* = 0, illustrated by the overlapping red, purple, and grey curves in the right-hand plot of Fig. 2(A). However, if ∆*ε*_*AI*_ is influenced by the mutation (either alone or in conjunction with *K*_*A*_ and *K*_*I*_), the leakiness will change, resulting in a non-zero ∆*F* when *c* = 0. This is illustrated in Fig. 2 (B) where ∆*ε*_*AI*_ is the only parameter affected by the mutation.

It is important to note that for a mutation which perturbs only the inducer binding constants, the dependence of ∆*F* on the inducer concentration can be non-monotonic. While the precise values of *K*_*A*_ and *K*_*I*_ control the sensitivity of the repressor to inducer concentration, it is the ratio *K*_*A*_/*K*_*I*_ that defines whether this non-monotonic behavior is observed. This can be seen more clearly when we consider the limit of saturating inducer concentration,

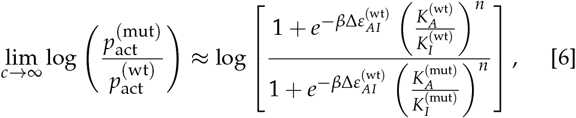

which illustrates that ∆*F* returns to zero at saturating inducer concentration when *K*_*A*_/*K*_*I*_ is the same for both the mutant and wild-type repressors, so long as ∆*ε*_*AI*_ is unperturbed. Non-monotonicity can *only* be achieved by changing *K*_*A*_ and *K*_*I*_ and therefore serves as a diagnostic for classifying mutational effects reliant solely on measuring the change in free energy.

The second term in Eq. (5) captures how changes in the repressor copy number contributes to changes in free energy. It is important to note that this contribution to the free energy change depends on the total number of repressors in the cell, not just those in the active state. This emphasizes that changes in the expression of the repressor are energetically divorced from changes to the allosteric nature of the repressor. As a consequence, the change in free energy is constant for all inducer concentrations, as is schematized in Fig. 2(C). Because magnitude of the change in free energy scales logarithmically with changing repressor copy number, a mutation which increases expression from 1 to 10 repressors per cell is more impactful from an energetic standpoint (*k*_*B*_*T* log(10) ≈ 2.3 *k*_*B*_*T*) than an increase from 90 to 100 (*k*_*B*_*T* log(100/90) ≈ 0.1 *k*_*B*_*T*). Appreciable changes in the free energy only arise when variations in the repressor copy number are larger than or comparable to an order of magnitude. Changes of this magnitude are certainly possible from a single point mutation, as it has been shown that even synonymous substitutions can drastically change translation efficiency (17).

The third and final term in Eq. (5) is the difference in the DNA binding energy between the mutant and wild-type repressors. All else being equal, if the mutated state binds more tightly to the DNA than the wild type (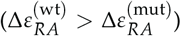, the net change in the free energy is negative, indicating that the repressor bound states become more energetically favorable due to the mutation. Much like in the case of changing repressor copy number, this quantity is independent of inducer concentration and is therefore also constant [Fig. 2(D)]. However, the magnitude of the change in free energy is linear with DNA binding affinity while it is logarithmic with respect to changes in the repressor copy number. Thus, to change the free energy by 1 *k*_*B*_*T*, the repressor copy number must change by a factor of ≈ 2.3 whereas the DNA binding energy must change by 1 *k*_*B*_*T*.

The unique behavior of each quantity in Eq. (5) and its sensitivity with respect to the parameters makes ∆*F* useful as a diagnostic tool to classify mutations. Given a set of fold-change measurements, a simple rearrangement of Eq. (3) permits the direct calculation of the free energy, assuming that the underlying physics of the regulatory architecture has not changed. Thus, it becomes possible to experimentally test the general assertions made in Fig. 2.

### DNA Binding Domain Mutations

With this arsenal of analytic diagnostics, we can begin to explore the mutational space of the repressor and map these mutations to the biophysical parameters they control. As one of the most thoroughly studied transcription factors, LacI has been subjected to numerous crystallographic and mutational studies (18–21). One such work generated a set of point mutations in the LacI repressor and examined the diversity of the phenotypic response to different allosteric effectors (5). However, experimental variables such as the repressor copy number or the number of specific binding sites were not known, making precise calculation of ∆*F* as presented here not tractable. Using this dataset as a guide, we chose a subset of the mutations and inserted them into our experimental strains of *E. coli* where these parameters are known and tightly controlled (4, 10).

We made three amino acid substitutions (Y20I, Q21A, and Q21M) that are critical for the DNA-repressor interaction. These mutations were introduced into the *lacI* sequence used in Garcia and Phillips 2011 (4) with four different ribosomal binding site sequences that were shown (via quantitative Western blotting) to tune the wild-type repressor copy number across three orders of magnitude. These mutant constructs were integrated into the *E. coli* chromosome harboring a Yellow Fluorescent Protein (YFP) reporter. The YFP promoter included the native O2 LacI operator sequence which the wild-type LacI repressor binds with high specificity (∆*ε*_*RA*_ = −13.9 *k*_*B*_*T*). The fold-change in gene expression for each mutant across twelve concentrations of IPTG was measured via flow cytometry. As we mutated only a single amino acid with the minimum number of base pair changes to the codons from the wild-type sequence, we find it unlikely that the repressor copy number was drastically altered from those reported in (4) for the wild-type sequence paired with the same ribosomal binding site sequences. In characterizing the effects of these DNA binding mutations, we take the repressor copy number to be unchanged. Any error introduced by this mutation should be manifest as a larger than predicted systematic shift in the free energy change when the repressor copy number is varied.

A naïve hypothesis for the effect of a mutation in the DNA binding domain is that *only* the DNA binding energy is altered. This hypothesis appears to contradict the core principle of allostery in that ligand binding in one domain influences binding in another, suggesting that changing any parameter modifies them all. The characteristic curves summarized in Fig. 2 give a means to discriminate between these two hypotheses by examining the change in the free energy. Using a single induction profile (white-faced points in Fig. 3), we estimated the DNA binding energy using a Bayesian approach, the details of which are discussed in the Materials and Methods as well as the SI text. The shaded red region for each mutant in Fig. 3 represents the 95% credible region of this fit whereas all other shaded regions are 95% credible regions of the predictions for other repressor copy numbers. We find that redetermining only the DNA binding energy accurately captures the majority of the induction profiles, indicating that other parameters are unaffected. One exception is for the lowest repressor copy numbers (*R* = 60 and *R* = 124 per cell) of mutant Q21A at low concentrations of IPTG. However, we note that this disagreement is comparable to that observed for the wild-type repressor binding to the weakest operator in Razo-Mejia *et al.* 2018 (10), illustrating that our model is imperfect in characterizing weakly repressing architectures. Including other parameters in the fit (such as ∆*ε*_*AI*_) does not significantly improve the accuracy of the predictions. Furthermore, the magnitude of this disagreement also depends on the choice of the fitting strain (see SI text).

**Fig. 3.**
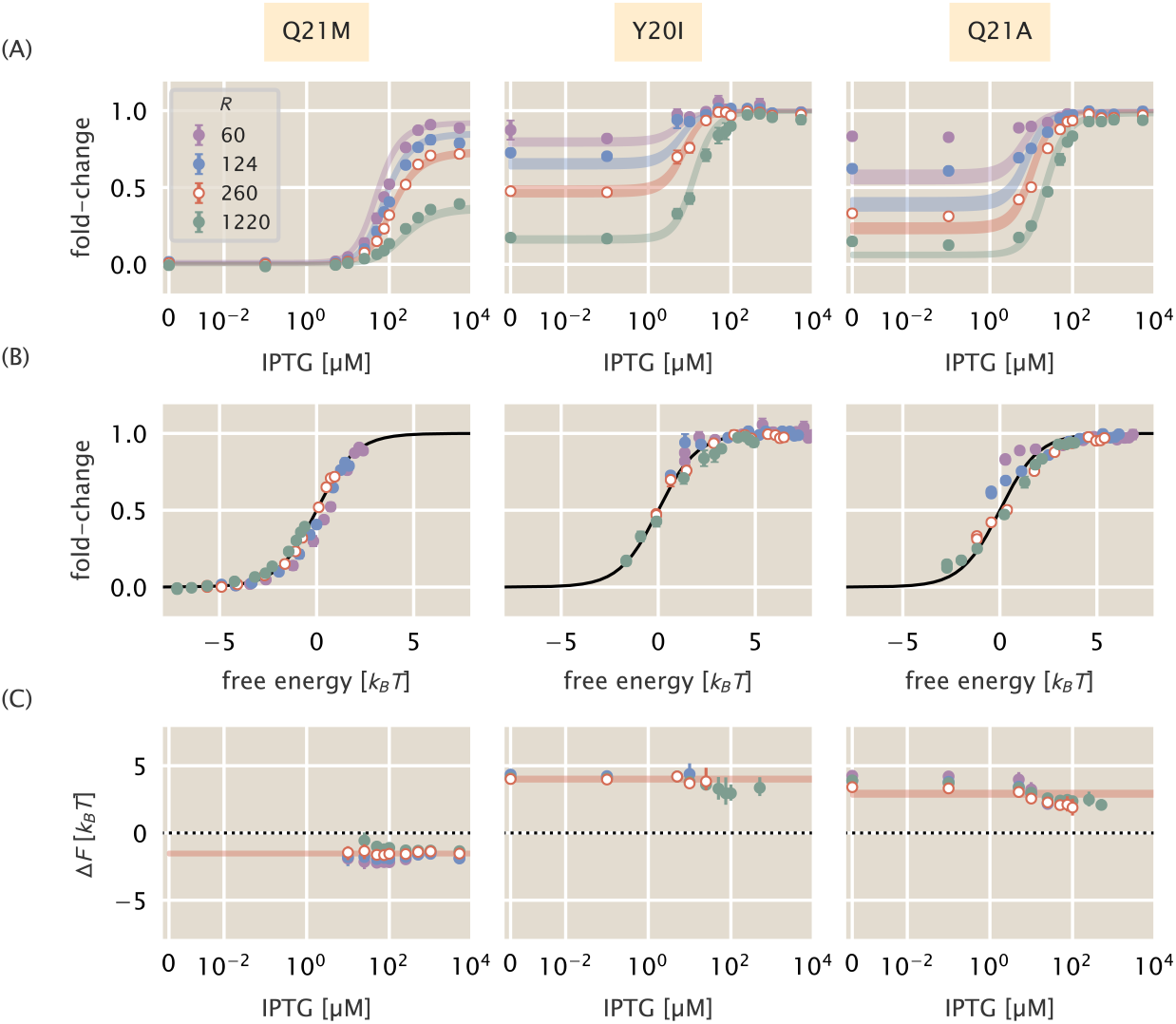
Induction profiles and free energy modifications of DNA binding domain mutations. Each column corresponds to the highlighted mutant at the top of the figure. Each strain was paired with the native O2 operator sequence. White-faced points correspond to the strain for each mutant from which the DNA binding energy was estimated. (A) Induction profiles of each mutant at four different repressor copy numbers as a function of the inducer concentration. Points correspond to the mean fold-change in gene expression of six to ten biological replicates. Error bars are the standard error of the mean. Shaded regions demarcate the 95% credible region of the induction profile generated by the estimated DNA binding energy. (B) Data collapse of all points for each mutant shown in (A) using only the DNA binding energy estimated from a single repressor copy number. Points correspond to the average fold-change in gene expression of six to ten biological replicates. Error bars are standard error of the mean. Where error bars are not visible, the relative error in measurement is smaller than the size of the marker. (C) The change in the free energy resulting from each mutation as a function of the inducer concentration. Points correspond to the median of the marginal posterior distribution for the free energy. Error bars represent the upper and lower bounds of the 95% credible region. Points in (A) at the detection limits of the flow cytometer (near fold-change values of 0 and 1) were neglected for calculation of the ∆*F*. The IPTG concentration is shown on a symmetric log scale with linear scaling ranging from 0 to 10^−2^ *µ*M and log scaling elsewhere.

Mutations Y20I and Q21A both weaken the affinity of the repressor to the DNA relative to the wild type strain (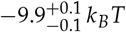 and 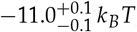, respectively). Here we report the median of the inferred posterior probability distribution with the superscripts and subscripts corresponding to the upper and lower bounds of the 95% credible region. These binding energies are comparable to that of the wild-type repressor affinity to the native LacI operator sequence O3, with a DNA binding energy of −9.7 *k*_*B*_*T*. The mutation Q21M increases the strength of the DNA-repressor interaction relative to the wild-type repressor with a binding energy of 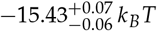, comparable to the affinity of the wild-type repressor to the native O1 operator sequence (−15.3*k*_*B*_*T*). It is notable that a single amino acid substitution of the repressor is capable of changing the strength of the DNA binding interaction well beyond that of many single base-pair mutations in the operator sequence (4, 22).

Using the new DNA binding energies, we can collapse all measurements of fold-change as a function of the free energy as shown in Fig. 3(B). This allows us to test the diagnostic power of the decomposition of the free energy described in Fig. 2. To compute the ∆*F* for each mutation, we inferred the observed mean free energy of the mutant strain for each inducer concentration and repressor copy number (see Materials and Methods as well as the SI text for a detailed explanation of the inference). We note that in the limit of extremely low or high fold-change, the inference of the free energy is either over-or under-estimated, respectively, introducing a systematic error. Thus, points which are close to these limits are omitted in the calculation of ∆*F*. We direct the reader to the SI text for a detailed discussion of this systematic error. With a measure of *F*^(mut)^ for each mutant at each repressor copy number, we compute the difference in free energy relative to the wild-type strain with the same repressor copy number and operator sequence, restricting all variability in ∆*F* solely to changes in ∆*ε*_*RA*_.

The change in free energy for each mutant is shown in Fig. 3(C). It can be seen that the ∆*F* for each mutant is constant as a function of the inducer concentration and is concordant with the prediction generated from fitting ∆*ε*_*RA*_ to a single repressor copy number [red lines Fig. 3(C)]. This is in line with the predictions outlined in Fig. 2(C) and (D), indicating that the allosteric parameters are “insulated”, meaning they are not affected by the DNA binding domain mutations. As the ∆*F* for all repressor copy numbers collapses onto the prediction, we can say that the expression of the repressor itself is the same or comparable with that of the wild type. If the repressor copy number were perturbed in addition to ∆*ε*_*RA*_, one would expect a shift away from the prediction that scales logarithmically with the change in repressor copy number. However, as the ∆*F* is approximately the same for each repressor copy number, it can be surmised that the mutation does not significantly change the expression or folding efficiency of the repressor itself. These results allow us to state that the DNA binding energy ∆*ε*_*RA*_ is the only parameter modified by the DNA mutants examined.

### Inducer Binding Domain Mutations

Much as in the case of the DNA binding mutants, we cannot safely assume *a priori* that a given mutation in the inducer binding domain affects only the inducer binding constants *K*_*A*_ and *K*_*I*_. While it is easy to associate the inducer binding constants with the inducer binding domain, the critical parameter in our allosteric model ∆*ε*_*AI*_ is harder to restrict to a single spatial region of the protein. As *K*_*A*_, *K*_*I*_, and ∆*ε*_*AI*_ are all parameters dictating the allosteric response, we consider two hypotheses in which inducer binding mutations alter either all three parameters or only *K*_*A*_ and *K*_*I*_.

We made four point mutations within the inducer binding domain of LacI (F164T, Q294V, Q294R, and Q294K) that have been shown previously to alter binding to multiple allosteric effectors (5). In contrast to the DNA binding domain mutants, we paired the inducer binding domain mutations with the three native LacI operator sequences (which have various affinities for the repressor) and a single ribosomal binding site sequence. This ribosomal binding site sequence, as reported in (4), expresses the wild-type LacI repressor to an average copy number of approximately 260 per cell. As the free energy differences resulting from point mutations in the DNA binding domain can be described solely by changes to ∆*ε*_*RA*_, we continue under the assumption that the inducer binding domain mutations do not significantly alter the repressor copy number.

The induction profiles for these four mutants are shown in Fig. 4(A). Of the mutations chosen, Q294R and Q294K appear to have the most significant impact, with Q294R abolishing the characteristic sigmoidal titration curve entirely. It is notable that both Q294R and Q294K have elevated expression in the absence of inducer compared to the other two mutants paired with the same operator sequence. Panel (A) in Fig. 2 illustrates that if only *K*_*A*_ and *K*_*I*_ were being affected by the mutations, the fold-change should be identical for all mutants in the absence of inducer. This discrepancy in the observed leakiness immediately suggests that more than *K*_*A*_ and *K*_*I*_ are affected for Q294K and Q294R.

**Fig. 4.**
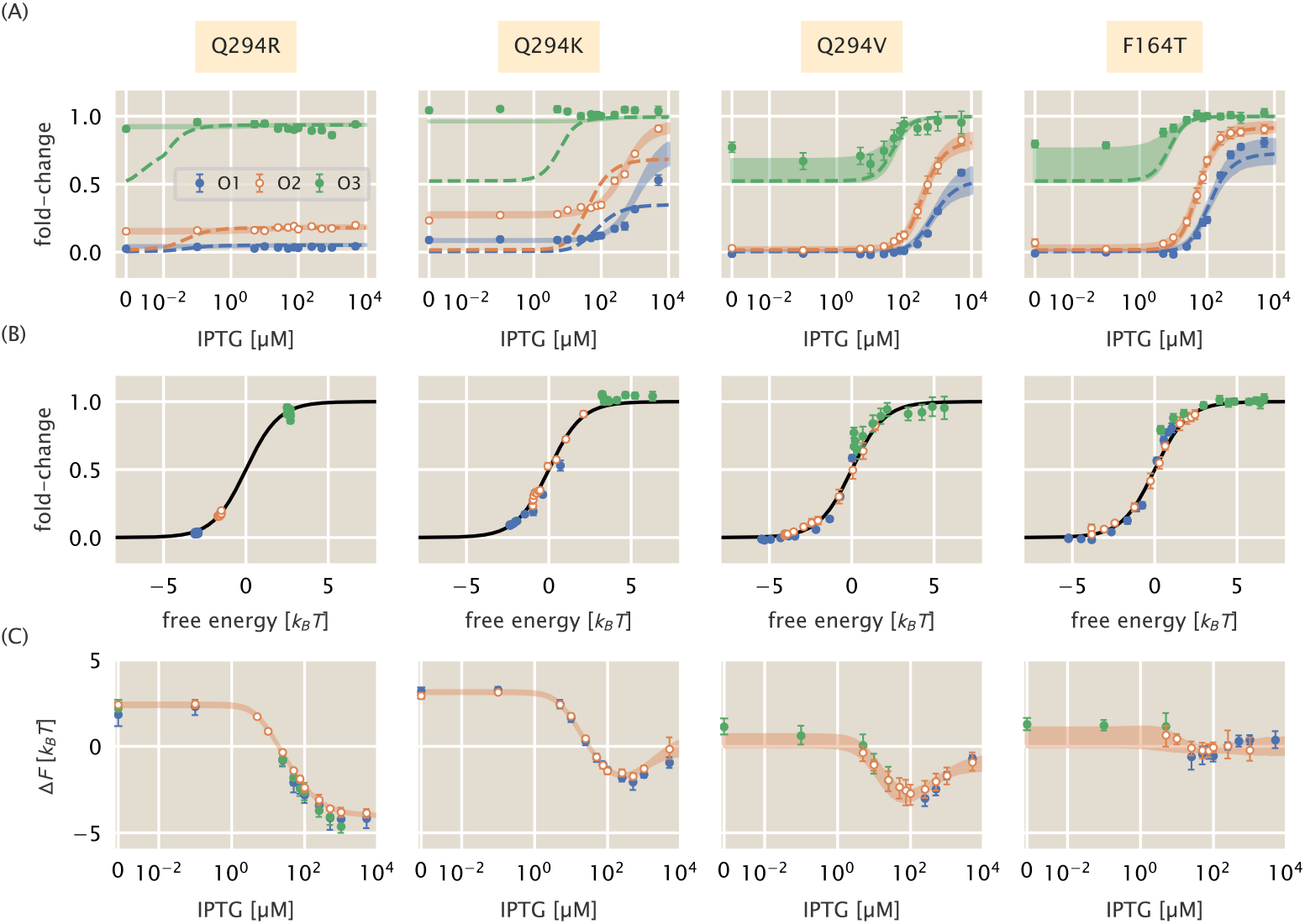
Induction profiles and free energy differences of inducer binding domain mutants. White faced points represent the strain to which the parameters were fit, namely the O2 operator sequence. Each column corresponds to the mutant highlighted at the top of the figure. All strains have *R* = 260 per cell. (A) The fold-change in gene expression as a function of the inducer concentration for three operator sequences of varying strength. Dashed lines correspond to the curve of best fit resulting from fitting *K*_*A*_ and *K*_*I*_ alone. Shaded curves correspond to the 95% credible region of the induction profile determined from fitting *K*_*A*_, *K*_*I*_, and ∆*ε*_*AI*_. Points correspond to the mean measurement of six to twelve biological replicates. Error bars are the standard error of the mean. (B) Points in (A) collapsed as a function of the free energy calculated from redetermining *K*_*A*_, *K*_*I*_, and ∆*ε*_*AI*_. (C) Change in free energy resulting from each mutation as a function of the inducer concentration. Points correspond to the median of the posterior distribution for the free energy. Error bars represent the upper and lower bounds of the 95% credible region. Shaded curves are the predictions. IPTG concentration is shown on a symmetric log scaling axis with the linear region spanning from 0 to 10^−2^ *µ*M and log scaling elsewhere.

Using a single induction profile for each mutant (shown in Fig. 4 as white-faced circles), we inferred the parameter combinations for both hypotheses and drew predictions for the induction profiles with other operator sequences. We find that the simplest hypothesis (in which only *K*_*A*_ and *K*_*I*_ are altered) does not permit accurate prediction of most induction profiles. These curves, shown as dotted lines in Fig. 4(A), fail spectacularly in the case of Q294R and Q294K, and undershoot the observed profiles for F164T and Q294V, especially when paired with the weak operator sequence O3. The change in the leakiness for Q294R and Q294K is particularly evident as the expression at *c* = 0 should be identical to the wild-type repressor under this hypothesis. Altering only *K*_*A*_ and *K*_*I*_ is not sufficient to accurately predict the induction profiles for F164T and Q294V, but not to the same degree as Q294K and Q294R. The disagreement is most evident for the weakest operator O3 [green lines in Fig. 4(A)], though we have discussed previously that the induction profiles for weak operators are difficult to accurately describe and can result in comparable disagreement for the wild-type repressor (10, 22).

Including ∆*ε*_*AI*_ as a perturbed parameter in addition to *K*_*A*_ and *K*_*I*_ improves the predicted profiles for all four mutants. By fitting these three parameters to a single strain, we are able to accurately predict the induction profiles of other operators as seen by the shaded lines in Fig. 4(A). With these modified parameters, all experimental measurements collapse as a function of their free energy as prescribed by Eq. (3) [Fig. 4(B)]. All four mutations significantly diminish the binding affinity of both states of the repressor to the inducer, as seen by the estimated parameter values reported in Tab. 1. As evident in the data alone, Q294R abrogates inducibility outright (*K*_*A*_ ≈ *K*_*I*_). For Q294K, the active state of the repressor can no longer bind inducer whereas the inactive state binds with weak affinity. The remaining two mutants, Q294V and F164T, both show diminished binding affinity of the inducer to both the active and inactive states of the repressor relative to the wild-type.

**Table 1.** Inferred values of *K*_*A*_, *K*_*I*_, and ∆*ε*_*AI*_ for inducer binding mutants

| Mutant | $K_A$ | $K_I$ | $\Delta\epsilon_{AI}$ [$k_B T$] | Reference |
| --- | --- | --- | --- | --- |
| WT | $139^{+29}_{-22} \mu\text{M}$ | $0.53^{+0.04}_{-0.04} \mu\text{M}$ | 4.5 | (10) |
| F164T | $165^{+90}_{-65} \mu\text{M}$ | $3^{+6}_{-3} \mu\text{M}$ | $1^{+5}_{-2}$ | This study |
| Q294V | $650^{+450}_{-250} \mu\text{M}$ | $8^{+8}_{-8} \mu\text{M}$ | $3^{+6}_{-3}$ | This study |
| Q294K | > 1 mM | $310^{+70}_{-60} \mu\text{M}$ | $-3.11^{+0.07}_{-0.07}$ | This study |
| Q294R | $9^{+20}_{-9} \mu\text{M}$ | $8^{+20}_{-8} \mu\text{M}$ | $-2.35^{+0.01}_{-0.09}$ | This study |

Given the collection of fold-change measurements, we computed the ∆*F* relative to the wild-type strain with the same operator and repressor copy number. This leaves differences in *p*_*act*_(*c*) as the sole contributor to the free energy difference, assuming our hypothesis that *K*_*A*_, *K*_*I*_, and ∆*ε*_*AI*_ are the only perturbed parameters is correct. The change in free energy can be seen in Fig. 4(C). For all mutants, the free energy difference inferred from the observed fold-change measurements falls within error of the predictions generated under the hypothesis that *K*_*A*_, *K*_*I*_, and ∆*ε*_*AI*_ are all affected by the mutation [shaded curves in Fig. 4(C)]. The profile of the free energy change exhibits some of the rich phenomenology illustrated in Fig. 2(A) and (B). Q294K, F164T, and Q294V exhibit a non-monotonic dependence on the inducer concentration, a feature that can only appear when *K*_*A*_ and *K*_*I*_ are altered. The non-zero ∆*F* at *c* = 0 for Q294R and Q294K coupled with an inducer concentration dependence is a telling sign that ∆*ε*_*AI*_ must be significantly modified. This shift in ∆*F* is positive in all cases, indicating that ∆*ε*_*AI*_ must have decreased, and that the inactive state has become more energetically favorable for these mutants than for the wild-type protein. Indeed the estimates for ∆*ε*_*AI*_ (Tab. 1) reveal both mutations Q294R and Q294K make the inactive state more favorable than the active state. Thus, for these two mutations, only ≈ 10% of the repressors are active in the absence of inducer, whereas the basal active fraction is ≈ 99% for the wild-type repressor (10).

Taken together, these parametric changes diminish the response of the regulatory architecture as a whole to changing inducer concentrations. They furthermore reveal that the parameters which govern the allosteric response are interdependent and no single parameter is insulated from the others. However, as *only* the allosteric parameters are changed, one can say that the allosteric parameters as a whole are insulated from the other components which define the regulatory response, such as repressor copy number and DNA binding affinity.

### Predicting Effects of Pairwise Double Mutations

Given full knowledge of each individual mutation, we can draw predictions of the behavior of the pairwise double mutants with no free parameters based on the simplest null hypothesis of no epistasis. The formalism of ∆*F* defined by Eq. (5) explicitly states that the contribution to the free energy of the system from the difference in DNA binding energy and the allosteric parameters are strictly additive. Thus, deviations from the predicted change in free energy would suggest epistatic interactions between the two mutations.

To test this additive model, we constructed nine double mutant strains, each having a unique inducer binding (F164T, Q294V, Q294K) and DNA binding mutation (Y20I, Q21A, Q21M). To make predictions with an appropriate representation of the uncertainty, we computed a large array of induction profiles given random draws from the posterior distribution for the DNA binding energy (determined from the single DNA binding mutants) as well as from the joint posterior for the allosteric parameters (determined from the single inducer binding mutants). These predictions, shown in Fig. 5(A) and (B) as shaded blue curves, capture all experimental measurements of the fold-change [Fig. 5(A)] and the inferred difference in free energy [Fig. 5(B)]. The latter indicates that there are no epistatic interactions between the mutations queried in this work, though if there were, systematic deviations from these predictions would shed light on how the epistasis is manifest.

**Fig. 5.**
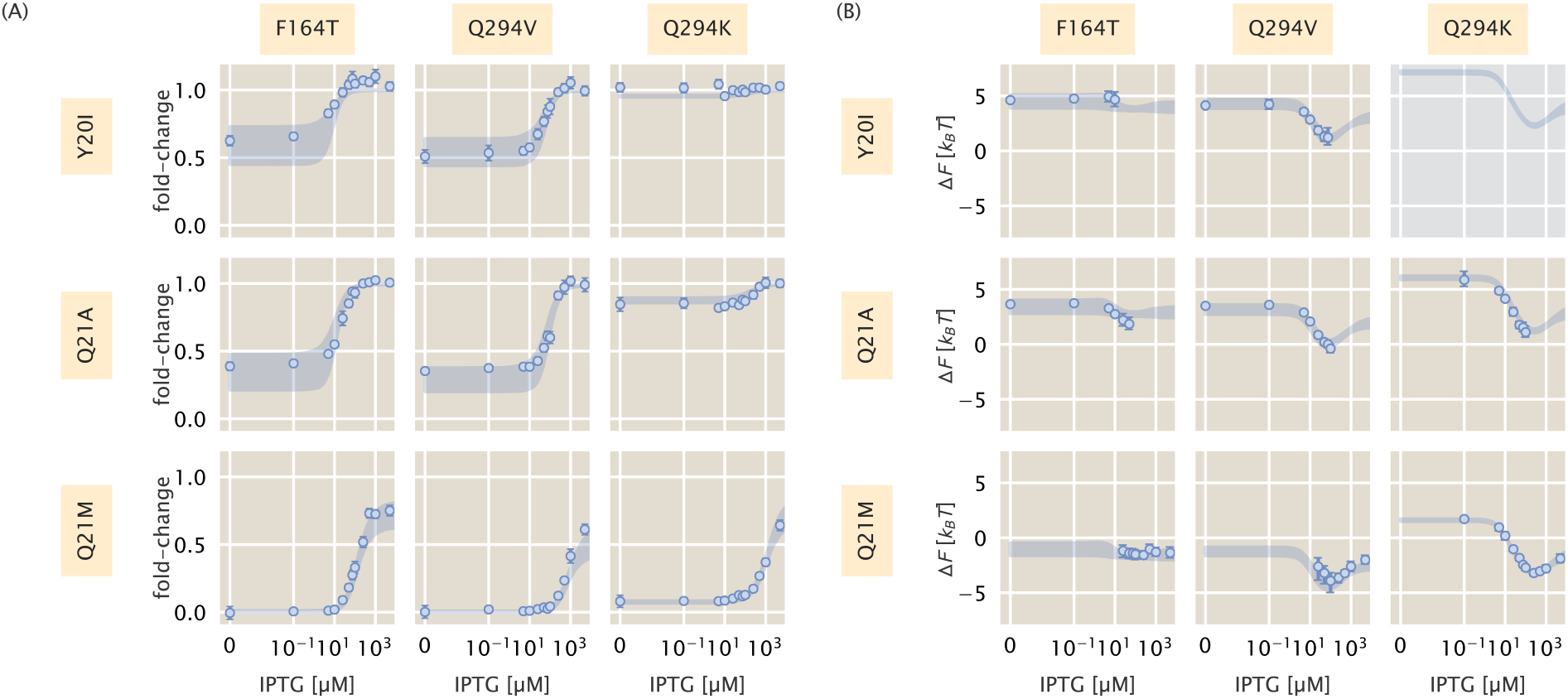
Induction and free energy profiles of DNA binding and inducer binding double mutants.(A) Fold-change in gene expression for each double mutant as a function of IPTG. Points and errors correspond to the mean and standard error of six to ten biological replicates. Where not visible, error bars are smaller than the corresponding marker. Shaded regions correspond to the 95% credible region of the prediction given knowledge of the single mutants. These were generated by drawing 10^4^ samples from the ∆*ε*_*RA*_ posterior distribution of the single DNA binding domain mutants and the joint probability distribution of *K*_*A*_, *K*_*I*_, and ∆*ε*_*AI*_ from the single inducer binding domain mutants. (B) The difference in free energy of each double mutant as a function of the reference free energy. Points and errors correspond to the median and bounds of the 95% credible region of the posterior distribution for the inferred ∆*F*. Shaded lines region are the predicted change in free energy, generated in the same manner as the shaded lines in (A). All measurements were taken from a strain with 260 repressors per cell paired with a reporter with the native O2 LacI operator sequence. In all plots, the IPTG concentration is shown on a symmetric log axis with linear scaling between 0 and 10^−2^ *µ*M and log scaling elsewhere.

The precise agreement between the predictions and measurements for Q294K paired with either Q21A or Q21M is striking as Q294K drastically changed ∆*ε*_*AI*_ in addition to *K*_*A*_ and *K*_*I*_. Our ability to predict the induction profile and free energy change underscores the extent to which the DNA binding energy and the allosteric parameters are insulated from one another. Despite this insulation, the repressor still functions as an allosteric molecule, emphasizing that the mutations we have inserted do not alter the pathway of communication between the two domains of the protein. As the double mutant Y20I-Q294K exhibits fold-change of approximately 1 across all IPTG concentrations [Fig. 5(A)], these mutations in tandem make repression so weak it is beyond the limits which are detectable by our experiments. As a consequence, we are unable to estimate ∆*F* nor experimentally verify the corresponding prediction [grey box in Fig. 5(B)]. However, as the predicted fold-change in gene expression is also approximately 1 for all *c*, we believe that the prediction shown for ∆*F* is likely accurate. One would be able to infer the ∆*F* to confirm these predictions using a more sensitive method for measuring the fold-change, such as single-cell microscopy or colorimetric assays.

### Discussion

Allosteric regulation is often couched as “biological action at a distance”. Despite extensive knowledge of protein structure and function, it remains difficult to translate the coordinates of the atomic constituents of a protein to the precise parameter values which define the functional response, making each mutant its own intellectual adventure. Bioinformatic approaches to understanding the sequence-structure relationship have permitted us to examine how the residues of allosteric proteins evolve, revealing conserved regions which hint to their function. Co-evolving residues reveal sectors of conserved interactions which traverse the protein that act as the allosteric communication channel between domains (23–25). Elucidating these sectors has advanced our understanding of how distinct domains “talk” to one another and has permitted direct engineering of allosteric responses into non-allosteric enzymes (26–28). Even so, we are left without a quantitative understanding of how these admittedly complex networks set the energetic difference between active and inactive states or how a given mutation influences binding affinity. In this context, a biophysical model in which the various parameters are intimately connected to the molecular details can be of use and can lead to quantitative predictions of the interplay between amino-acid identity and system-level response.

By considering how each parameter contributes to the observed change in free energy, we are able to tease out different classes of parameter perturbations which result in stereotyped responses to changing inducer concentration. These characteristic changes to the free energy can be used as a diagnostic tool to classify mutational effects. For example, we show in Fig. 2 that modulating the inducer binding constants *K*_*A*_ and *K*_*I*_ results in non-monotonic free energy changes that are dependent on the inducer concentration, a feature observed in the inducer binding mutants examined in this work. Simply looking at the inferred ∆*F* as a function of inducer concentration, which requires no fitting of the biophysical parameters, indicates that *K*_*A*_ and *K*_*I*_ must be modified considering those are the only parameters which can generate such a response.

Another key observation is that a perturbation to only *K*_*A*_ and *K*_*I*_ requires that the ∆*F* = 0 at *c* = 0. Deviations from this condition imply that more than the inducer binding constants must have changed. If this shift in ∆*F* off of 0 at *c* = 0 is not constant across all inducer concentrations, we can surmise that the energy difference between the allosteric states ∆*ε*_*AI*_ must also be modified. We again see this effect for all of our inducer mutants. By examining the inferred ∆*F*, we can immediately say that in addition to *K*_*A*_ and *K*_*I*_, ∆*ε*_*AI*_ must decrease relative to the wild-type value as ∆*F* > 0 at *c* = 0. When the allosteric parameters are fit to the induction profiles, we indeed see that this is the case, with all four mutations decreasing the energy gap between the active and inactive states. Two of these mutations, Q294R and Q294K, make the inactive state of the repressor *more* stable than the active state, which is not the case for the wild-type repressor (10).

Our formulation of ∆*F* indicates that shifts away from 0 that are independent of the inducer concentration can only arise from changes to the repressor copy number and/or DNA binding specificity, indicating that the allosteric parameters are untouched. We see that for three mutations in the DNA binding domain, ∆*F* is the same irrespective of the inducer concentration. Measurements of ∆*F* for these mutants with repressor copy numbers across three orders of magnitude yield approximately the same value, revealing that ∆*ε*_*RA*_ is the sole parameter altered via the mutations.

We note that the conclusions stated above can be qualitatively drawn without resorting to fitting various parameters and measuring the goodness-of-fit. Rather, the distinct behavior of ∆*F* is sufficient to determine which parameters are changing. Here, these conclusions are quantitatively confirmed by fitting these parameters to the induction profile, which results in accurate predictions of the fold-change and ∆*F* for nearly every strain across different mutations, repressor copy numbers, and operator sequence, all at different inducer concentrations. With a collection of evidence as to what parameters are changing for single mutations, we put our model to the test and drew predictions of how double mutants would behave both in terms of the titration curve and free energy profile.

A hypothesis that arises from our formulation of ∆*F* is that a simple summation of the energetic contribution of each mutation should be sufficient to predict the double mutants (so long as they are in separate domains). We find that such a calculation permits precise and accurate predictions of the double mutant phenotypes, indicating that there are no epistatic interactions between the mutations examined in this work. With an expectation of what the free energy differences should be, epistatic interactions could be understood by looking at how the measurements deviate from the prediction. For example, if epistatic interactions exist which appear as a systematic shift from the predicted ∆*F* independent of inducer concentration, one could conclude that DNA binding energy is not equal to that of the single mutation in the DNA binding domain alone. Similarly, systematic shifts that are dependent on the inducer concentration (i.e. not constant) indicate that the allosteric parameters must be influenced. If the expected difference in free energy is equal to 0 when *c* = 0, one could surmise that the modified parameter must not be ∆*ε*_*AI*_ nor ∆*ε*_*RA*_ as these would both result in a shift in leakiness, indicating that *K*_*A*_ and *K*_*I*_ are further modified.

Ultimately, we present this work as a proof-of-principle for using biophysical models to investigate how mutations influence the response of allosteric systems. We emphasize that such a treatment allows one to boil down the complex phenotypic responses of these systems to a single-parameter description which is easily interpretable as a free energy. The general utility of this approach is illustrated in Fig. 6 where gene expression data from previous work (4, 6, 10) along with all of the measurements presented in this work collapse onto the master curve defined by Eq. (3). While our model coarse grains many of the intricate details of transcriptional regulation into two states (one in which the repressor is bound to the promoter and one where it is not), it is sufficient to describe a wide range of regulatory scenarios. Given enough parametric knowledge of the system, it becomes possible to examine how modifications to the parameters move the physiological response along this reduced one-dimensional parameter space. This approach offers a glimpse at how mutational effects can be described in terms of energy rather than Hill coefficients and arbitrary prefactors. While we have explored a very small region of sequence space in this work, coupling of this approach with high-throughput sequencing-based methods to query a library of mutations within the protein will shed light on the phenotypic landscape centered at the wild-type sequence. Furthermore, pairing libraries of protein and operator sequence mutants will provide insight as to how the protein and regulatory sequence coevolve, a topic rich with opportunity for a dialogue between theory and experiment.

**Fig. 6.**
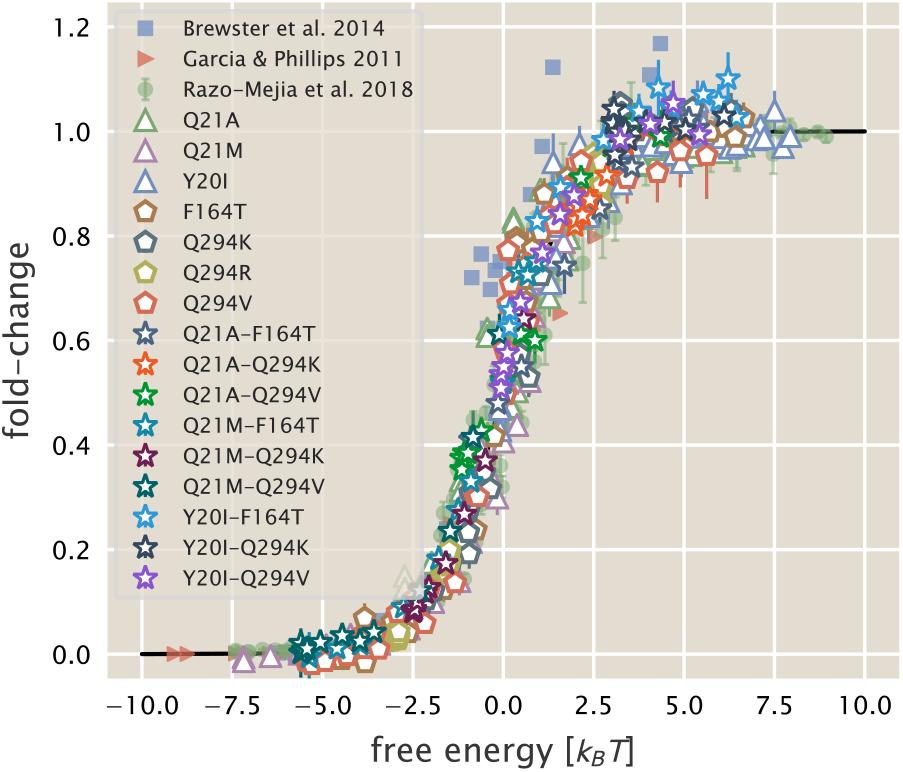
Data collapse of the simple repression regulatory architecture. All data are means of biological replicates. Where present, error bars correspond to the standard error of the mean of five to fifteen biological replicates. Red triangles indicate data from Garcia and Phillips (4) obtained by colorimetric assays. Blue squares are data from Brewster et al.(6) acquired from video microscopy. Green circles are data from Razo-Mejia et al. (10) obtained via flow cytometry. All other symbols correspond to the work presented here. An interactive version of this figure can be found on the paper website where the different data sets can be viewed in more detail.

## Materials and Methods

### Bacterial Strains and DNA Constructs

All wild-type strains from which the mutants were derived were generated in previous work from the Phillips group (4, 10). Briefly, mutations were first introduced into the *lacI* gene of our pZS3*1-lacI plasmid (4) using a combination of overhang PCR Gibson assembly as well as QuickChange mutagenesis (Agligent Technologies). The oligonucleotide sequences used to generate each mutant as well as the method are provided in the SI text.

For mutants generated through overhang PCR and Gibson assembly, oligonucleotide primers were purchased containing an overhang with the desired mutation and used to amplify the entire plasmid. Using the homology of the primer overhang, Gibson assembly was performed to circularize the DNA prior to electroporation into MG1655 *E. coli* cells. Integration of LacI mutants was performed with *l* Red recombineering (29) as described in Reference (4).

The mutants studied in this work were chosen from data reported in (5). In selecting mutations, we looked for mutants which suggested moderate to strong deviations from the behavior of the wild-type repressor. We note that the variant of LacI used in this work has an additional three amino acids (Met-Val-Asn) added to the N-terminus than the canonical LacI sequence reported in (30). For this reason, all mutants given here are with respect to our sequence and their positions are shifted by three to those studied in (5).

### Flow Cytometry

All fold-change measurements were performed on a MACSQuant flow cytometer as described in Razo-Mejia et al. (10). Briefly, saturated overnight cultures 500 *µ*L in volume were grown in deep-well 96 well plates covered with a breathable nylon cover (Lab Pak - Nitex Nylon, Sefar America, Cat. No. 241205). After approximately 12 to 15 hr, the cultures reached saturation and were diluted 1000-fold into a second 2 mL 96-deep-well plate where each well contained 500 *µ*L of M9 minimal media supplemented with 0.5% w/v glucose (anhydrous D-Glucose, Macron Chemicals) and the appropriate concentration of IPTG (Isopropyl *β*-D-1-thiogalactopyranoside, Dioxane Free, Research Products International). These were sealed with a breathable cover and were allowed to grow for approximately 8 hours until the OD_600nm_ ≈ 0.3. Cells were then diluted ten-fold into a round-bottom 96-well plate (Corning Cat. No. 3365) containing 90 *µ*L of M9 minimal media supplemented with 0.5% w/v glucose along with the corresponding IPTG concentrations.

The flow cytometer was calibrated prior to use with MACSQuant Calibration Beads (Cat. No. 130-093-607). During measurement, the cultures were held at approximately 4° C by placing the 96-well plate on a MACSQuant ice block. All fluorescence measurements were made using a 488 nm excitation wavelength with a 525/50 nm emission filter. The photomultiplier tube voltage settings for the instrument are the same as those used in Reference (10).

The data was processed using an automatic unsupervised gating procedure based on the front and side-scattering values, where we fit a two-dimensional Gaussian function to the log_10_ forward-scattering (FSC) and the log_10_ side-scattering (SSC) data. Here we assume that the region with highest density of points in these two channels corresponds to single-cell measurements and consider data points that fall within 40% of the highest density region of the two-dimensional Gaussian function. We direct the reader to Reference (10) for further detail and comparison of flow cytometry with single-cell microscopy.

### Bayesian Parameter Estimation

We used a Bayesian definition of probability in the statistical analysis of all mutants in this work. In the SI text, we derive in detail the statistical models used for the various parameters as well as multiple diagnostic tests. Here, we give a generic description of our approach. To be succinct in notation, we consider a generic parameter *θ* which represents ∆*ε*_*RA*_, *K*_*A*_, *K*_*I*_, and/or ∆*ε*_*AI*_ depending on the specific LacI mutant.

As prescribed by Bayes’ theorem, we are interested in the posterior probability distribution

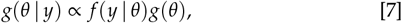

where we use *g* and *f* to represent probability densities over parameters and data, respectively, and *y* to represent a set of fold-change measurements. The likelihood of observing our dataset *y* given a value of *θ* is captured by *f* (*y* | *θ*). All prior information we have about the possible values of *θ* are described by *g*(*θ*).

In all inferential models used in this work, we assumed that all experimental measurements at a given inducer concentration were normally distributed about a mean value *µ* dictated by Eq. (1) with a variance *σ*^2^,

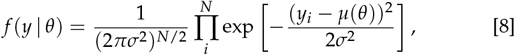

where *N* is the number of measurements in the data set *y*.

This choice of likelihood is justified as each individual measurement at a given inducer concentration is a biological replicate and independent of all other experiments. By using a Gaussian likelihood, we introduce another parameter *σ*. As *σ* must be positive and greater than zero, we define as a prior distribution a half-normal distribution with a standard deviation *ϕ*,

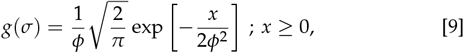

where *x* is a given range of values for *σ*. A standard deviation of *ϕ* = 0.1 was chosen given our knowledge of the scale of our measurement error from other experiments. As the absolute measurement of fold-change is restricted between 0 and 1.0, and given our knowledge of the sensitivity of the experiment, it is reasonable to assume that the error will be closer to 0 than to 1.0. Further justification of this choice of prior through simulation based methods are given in the SI text. The prior distribution for *θ* is dependent on the parameter and its associated physical and physiological restrictions. Detailed discussion of our chosen prior distributions for each model can also be found in the SI text.

All statistical modeling and parameter inference was performed using Markov chain Monte Carlo (MCMC). Specifically, Hamiltonian Monte Carlo sampling was used as sis implemented in the Stan probabilistic programming language (31). All statistical models saved as `.stan` models and can be accessed at the GitHub repository associated with this work (DOI: 10.5281/zenodo.2721798) or can be downloaded directly from the paper website.

### Inference of Free Energy From Fold-Change Data

While the fold-change in gene expression is restricted to be between 0 and 1, experimental noise can generate fold-change measurements beyond these bounds. To determine the free energy for a given set of fold-change measurements (for one unique strain at a single inducer concentration), we modeled the observed fold-change measurements as being drawn from a normal distribution with a mean *µ* and standard deviation *σ*. Using Bayes’ theorem, we can write the posterior distribution as

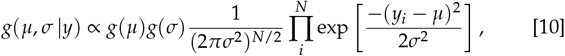

where *y* is a collection of fold-change measurements. The prior distribution for *µ* was chosen to be uniform between 0 and 1 while the prior on *σ* was chosen to be half normal, as written in Eq. (9). The posterior distribution was sampled independently for each set of fold-change measurements using MCMC. The. stan model for this inference is available on the paper website.

For each MCMC sample of *µ*, the free energy was calculated as

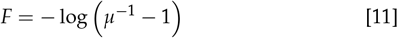

which is simply the rearrangement of Eq. (3). Using simulated data, we determined that when *µ* < *σ* or (1 − *µ*) < *σ*, the mean fold-change in gene expression was over or underestimated for the lower and upper limit, respectively. This means that there are maximum and minimum levels of fold-change that can be detected using flow cytometry which are set by the distribution of fold-change measurements resulting from various sources of day-to-day variation. This results in a systematic error in the calculation of the free energy, making proper inference beyond these limits difficult. This bounds the range in which we can confidently infer this quantity with flow cytometry. We hypothesize that more sensitive methods, such as single cell microscopy, colorimetric assays, or direct counting of mRNA transcripts via Fluorescence *In Situ* Hybridization (FISH) would improve the measurement of ∆*F*. We further discuss details of this limitation in the SI text.

### Data and Code Availability

All data was collected, stored, and preserved using the Git version control software. Code for data processing, analysis, and figure generation is available on the GitHub repository (https://www.github.com/rpgroup-pboc/mwc_mutants) or can be accessed via the paper website. Raw flow cytometry data is stored on the CaltechDATA data repository and can be accessed via DOI 10.22002/D1.1241.

## ACKNOWLEDGMENTS

We thank Pamela Björkman, Rachel Galimidi, and Priyanthi Gnanapragasam for access and training for the use of the Miltenyi Biotec MACSQuant flow cytometer. The experimental efforts first took place at the Physiology summer course at the Marine Biological Laboratory in Woods Hole, MA, operated by the University of Chicago. We thank Ambika Nadkarni and Damian Dudka for their work on the project during the course. We also thank Suzannah Beeler, Justin Bois, Robert Brewster, Soichi Hirokawa, Heun Jin Lee, and Muir Morrison for thoughtful advice and discussion. This work was supported by La Fondation Pierre-Gilles de Gennes, the Rosen Center at Caltech, the NIH DP1 OD0002179 (Director’s Pioneer Award), R01 GM085286, and 1R35 GM118043 (MIRA). Nathan M. Belliveau was supported by a Howard Hughes Medical Institute International Student Research fellowship.

## Supporting Information Text

### 1. Derivation of the Simple Repression Input-Output Function

In this section, we derive the input-output function for the inducible simple repression motif. This section summarizes the results from Garcia and Phillips 2011 (1) and Razo-Mejia et al. 2018 (2) and we direct the reader to these references for further detail.

We begin by defining the simple repression motif as a regulatory architecture in which binding of a repressor to its cognate binding site occludes binding of an RNA polymerase (RNAP) to the promoter, thereby hindering gene expression (3, 4). The repressor in this work is considered to be an allosteric molecule which fluctuates between an active and inactive state in thermal equilibrium. Binding of an allosteric effector molecule (i.e. an inducer) to a binding site in one domain of the repressor can stabilize the inactive state relative to the active state. The repressor can still bind the DNA in the inactive state but the sequence specificity is reduced and binding to the cognate sequence becomes comparable to nonspecific binding. Such regulatory motifs have been well characterized from a thermodynamic perspective in which the system is considered to be in equilibrium at the time scales relevant to molecular binding events. Under such a model, we can assume that the level of gene expression is proportional to the the probability of RNAP being bound to the promoter and this has frequently been applied in thermodynamic models of transcription (1–10).

The probability of the promoter being occupied by either a polymerase, repressor, or neither is dictated by the Boltzmann distribution,

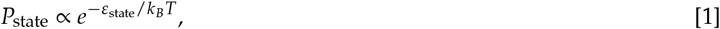

where *ε*_state_ is the energy of the state of interest. This energy is scaled to the thermal energy of the system *k*_*B*_*T* where *k*_*B*_ is Boltzmann’s constant and *T* is the temperature in units of *K*. The goal of this section is to translate generic proportionality in Eq. (1) into the relevant states of our system.

The occupancy states of the promoter and corresponding statistical weights can be seen in Fig. S1 (A). Here we use *P* to denote the number of RNAP per cell, *R*_*A*_ as the number of active repressors, and *R*_*I*_ as the number of inactive repressors. We assume there is a single specific binding site on the genome and *N*_*NS*_ nonspecific binding sites. The polymerase, active, and inactive repressor in this work are considered to bind the DNA with different strengths. The energy of binding for each species is given as ∆*ε*_*P*_, ∆*ε*_*RA*_, or ∆*ε*_*RI*_, which captures the energetic difference between nonspecific and specific binding for that species. Since we consider a single specific binding site for both the polymerase and repressor, we can say that *N*_*NS*_ >> *P* and *N*_*NS*_ >> *R*_*A*_ + *R*_*I*_, allowing the multiplicity of arranging *P* polymerases and *R* repressors to be approximately equal to *P*/*N*_*NS*_ and (*R*_*A*_ + *R*_*I*_)/*N*_*NS*_, respectively.

**Fig. S1.**
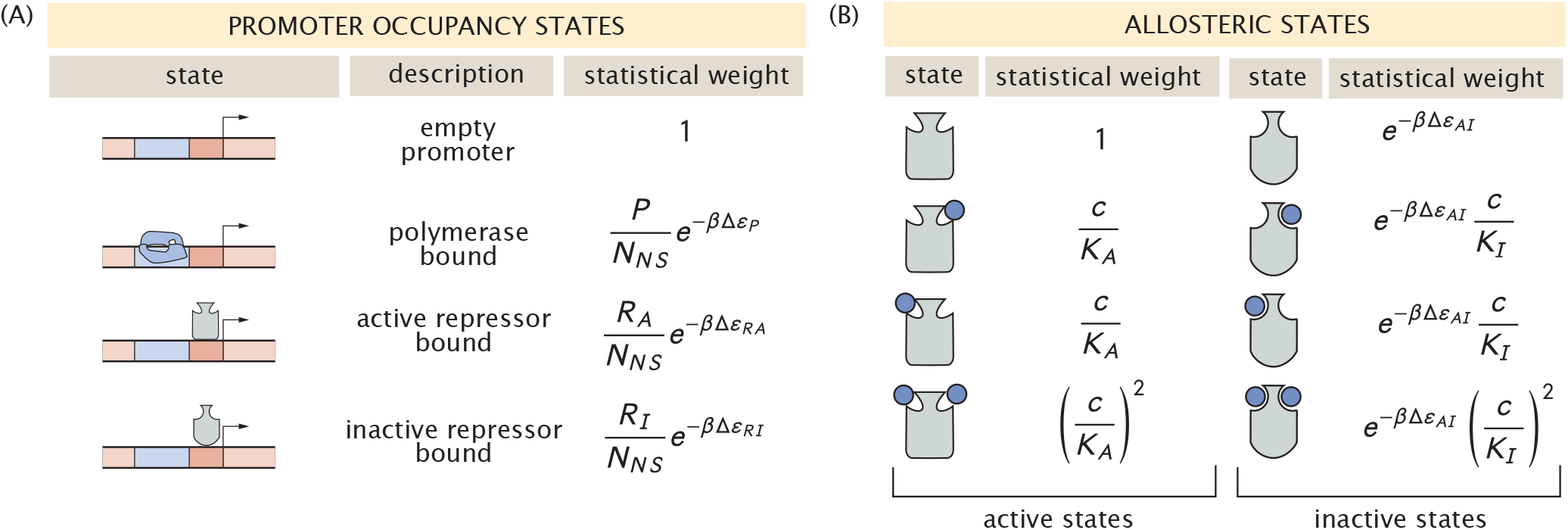
States and statistical weights for a simple repression motif with an allosteric repressor. (A) Occupancy states of the promoter in which the statistical weights are relative to the unoccupied state. *P*, *R*_*A*_, and *R*_*I*_ correspond to the average number of RNAP, active repressors, and inactive repressors per cell, respectively. The relative DNA binding energies are given as ∆*ε*_*P*_, ∆*ε*_*RA*_ and ∆*ε*_*RI*_. (B) Allosteric states of the repressor and the statistical weights relative to the active repressor with no inducer bound. The dissociation constant for the inducer to the active and inactive state of the repressor are denoted as *K*_*A*_ and *K*_*I*_, respectively, *c* is the inducer concentration, and ∆*ε*_*AI*_ is the energetic difference between the active and inactive states of the repressor.

With these states and statistical weights enumerated, we can now define the probability of a polymerase being bound to the promoter as

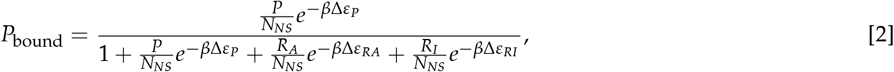

where we have defined *β* as 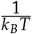.

It is experimentally difficult to measure *P*_bound_ directly as identifying the direct proportionality to gene expression is not straightforward. However, we can easily measure the fold-change in gene expression, defined as the probability of a polymerase being bound to the promoter in the presence of repressor relative to constitutive expression,

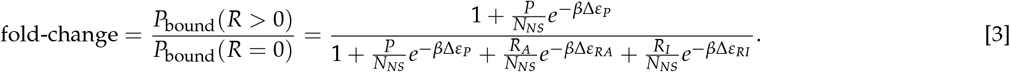

This can be simplified by making two well justified approximations. We can assume that binding of the inactive repressor to the specific binding site is approximately equal to nonspecific binding, 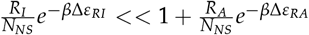. Secondly, we can state that binding of RNAP to the promoter is weak, 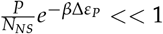. Assuming *P* ≈ 10^3^ (11), *N*_*NS*_ ≈ 4.6 × 10^6^ (the length of the *E. coli* genome in base pairs) and ∆*ε*_*P*_ ≈ −2 to −5*k*_*B*_*T* (12), the probability of this state comes to ≈ 1% and can be neglected. Using these approximations, we can state that the fold-change in gene expression has the form

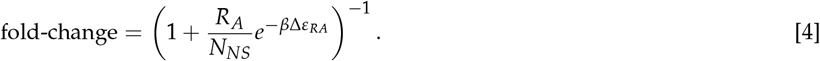

In order to make falsifiable predictions, we must have a precise knowledge of the number of active repressors in the cell *R*_*A*_. While determining this quantity is fraught with experimental difficulties, it is relatively easy to determine the total number of repressors per cell *R* through quantitative western blotting (1), fluorescence based methods (9), or proteomic studies (13). We can compute the number of active repressors at a given inducer concentration *c* by multiplying the total number of repressors by the probability of a repressor being active at that inducer concentration,

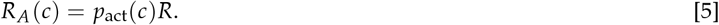

Similarly to computing *P*_bound_, we can compute the possible states and statistical weights of the repressor activity, shown in Fig. S1 (B). Following the model of Monod, Wyman, and Changeux (14), we have defined all statistical weights relative to the active repressor with no bound inducer molecules. We have defined the dissociation constant of the inducer to the active and inactive repressor as *K*_*A*_ and *K*_*I*_, respectively, and have assigned an energetic penalty 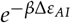 to all inactive states of the repressor. The energetic term ∆*ε*_*AI*_ represents the relative energy difference between the active and inactive states, ∆*ε*_*AI*_ = *ε*_*I*_ − *ε*_*A*_. For the *lac* repressor used in this work, the value of ∆*ε*_*AI*_ has been inferred to be 4.5 *k*_*B*_*T*, indicating that the active state is energetically preferred and with no inducer, approximately 99% of the repressors are in the active state.

Using these states and weights, we can compute *P*_act_(*c*) as

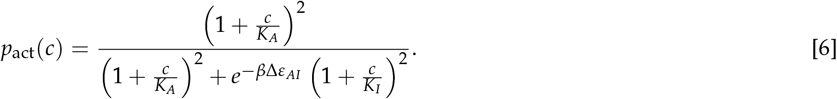

Using Eq. (4) - Eq. (6), we can then state that at a given inducer concentration *c*, the fold-change in gene expression can be defined as

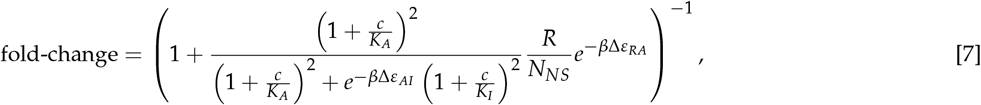

which is the result stated in Eq. 1 of the main text. We emphasize that equilibrium models as derived here have frequently been used to characterize the simple repression motif (1–5, 9) in addition to non-equilibrium approaches which have the same functional form as Eq. (7) (15).

### 2. Bayesian Parameter Estimation For DNA Binding Mutants

In this section, we outline the statistical model used in this work to estimate the DNA binding energy for a given mutation in the DNA binding domain. We begin with a derivation of our statistical model using Bayes’ theorem and then perform a series of principled steps to validate our choices of priors, ensure computational feasibility, and assess the validity of the model given the collected data. This work follows the analysis pipeline outlined by Michael Betancourt in his case-study entitled “Towards A Principled Bayesian Workflow.”

The second subsection *“Building a Generative Statistical Model”* lays out the statistical model used in this work to estimate the DNA binding energy and the error term *σ*. The subsequent subsections – *“Prior Predictive Checks”*, *“Simulation Based Calibration”*, and *“Posterior Predictive Checks”* – define and summarize a series of tests that ensure that the parameters of the statistical model can be identified and are computationally tractable. To understand how we defined our statistical model, only the second subsection is needed.

#### Calculation of the Fold-Change in Gene Expression

We appreciate the subtleties of the efficiency of photon detection in the flow cytometer, fluorophore maturation and folding, and autofluorescence correction, and we understand the importance in modeling the effects that these processes have on the reported value of the fold-change. However, in order to be consistent with the methods used in the literature, we took a more simplistic approach to calculate the fold-change. Given a set of fluorescence measurements of the constitutive expression control (*R* = 0), an autofluorescence control (no YFP), and the experimental strain (*R* > 0), we calculate the fold-change as

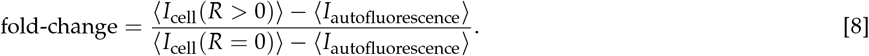

It is important to note here that for a given biological replicate, we consider only a point estimate of the mean fluorescence for each sample and perform a simple subtraction to adjust for background fluorescence. For the analysis going forward, all mentions of measured fold-change are determined by this calculation.

#### Building a Generative Statistical Model

To identify the minimal parameter set affected by a mutation, we assume that mutations in the DNA binding domain of the repressor alters only the DNA binding energy ∆*ε*_*RA*_, while the other parameters of the repressor are left unperturbed from their wild-type values. As a first approach, we can assume that all of the other parameters are known without error and can be taken as constants in our physical model. Ultimately, we want to know how probable a particular value of ∆*ε*_*RA*_ is given a set of experimental measurements *y*. Bayes’ theorem computes this distribution, termed the *posterior distribution* as

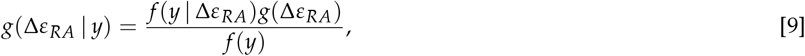

where we have used *g* and *f* to represent probability densities over parameters and data, respectively. The expression *f* (*y*|∆*ε*_*RA*_) captures the likelihood of observing our data set *y* given a value for the DNA binding energy under our physical model. All knowledge we have of what the DNA binding energy *could* be, while remaining completely ignorant of the experimental measurements, is defined in *g*(∆*ε*_*RA*_), referred to as the *prior distribution*. Finally, the likelihood that we would observe the data set *y* while being ignorant of our physical model is defined by the denominator *f* (*y*). In this work, this term serves only as a normalization factor and as a result will be treated as a constant. We can therefore say that the posterior distribution of ∆*ε*_*RA*_ is proportional to the joint distribution between the likelihood and the prior,

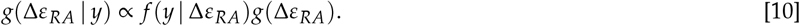

We are now tasked with translating this generic notation into a concrete functional form. Our physical model given by Eq. (7) computes the average fold-change in gene expression. Speaking practically, we make several replicate measurements of the fold-change to reduce the effects of random errors. As each replicate is independent of the others, it is reasonable to expect that these measurements will be normally distributed about the theoretical value of the fold-change *µ*, computed for a given Δ*ε*_*RA*_. We can write this mathematically for each measurement as

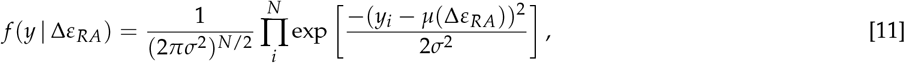

where *N* is the number of measurements in *y* and *y*_*i*_ is the *i*^th^ experimental fold-change measurement. We can write this likelihood in shorthand as

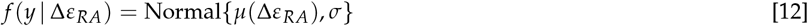

which we will use for the remainder of this section.

Using a normal distribution for our likelihood has introduced a new parameter *σ* which describes the spread of our measurements about the true value. We must therefore include it in our parameter estimation and assign an appropriate prior distribution such that the posterior distribution becomes

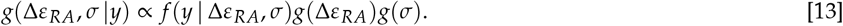

We are now tasked with assigning functional forms to the priors *g*(∆*ε*_*RA*_) and *g*(*σ*). Though one hopes that the result of the inference is not too dependent on the choice of prior, it is important to choose one that is in agreement with our physical and physiological intuition of the system.

We can impose physically reasonable bounds on the possible values of the DNA binding energy ∆*ε*_*RA*_. We can say that it is unlikely that any given mutation in the DNA binding domain will result in an affinity greater than that of biotin to streptavidin [1 fM ≈ −35 k_B_T, BNID 107139 (16)], one of the strongest known non-covalent bonds. Similarly, it’s unlikely that a given mutation will result in a large, positive binding energy, indicating non-specific binding is preferable to specific binding (~ 1 to 10 k_B_T). While it is unlikely for the DNA binding energy to exceed these bounds, it’s not impossible, meaning we should not impose these limits as hard boundaries. Rather, we can define a weakly informative prior as a normal distribution with a mean and standard deviation as the average of these bounds,

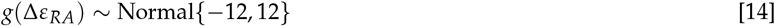

whose probability density function in shown in Fig. S2 (A).

By definition, fold-change is restricted to the bounds [0, 1]. Measurement noise and fluctuations in autofluorescence background subtraction means that experimental measurements of fold-change can extend beyond these bounds, though not substantially. By definition, the scale parameter *σ* must be positive and greater than zero. We also know that for the measurements to be of any use, the error should be less than the available range of fold-change, 1.0. We can choose such a prior as a half normal distribution

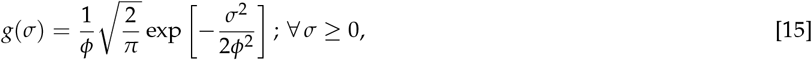

where *ϕ* is the standard deviation. By choosing *ϕ* = 0.1, it is unlikely that *σ* ≥ 1 yet not impossible, permitting the occasional measurement significantly outside of the theoretical bounds. The probability density function for this prior is shown in Fig. S2(B).

While these choices for the priors seem reasonable, we can check their appropriateness by using them to simulate a data set and checking that the hypothetical fold-change measurements obey our physical and physiological intuition.

#### Prior Predictive Checks

If our choice of prior distribution for each parameter is appropriate, we should be able to simulate data sets using these priors that match our expectations. In essence, we would hope that these prior choices would generate some data sets with fold-change measurements above 1 or below zero, but they should be infrequent. If we end up getting primarily negative values for fold-change, for example, then we can surmise that there is something wrong in our definition of the prior distribution. This method, coined a *prior predictive check*, was first put forward by Isidore Good in 1950 (17) and has received newfound attention in computational statistics.

We perform the simulation in the following manner. We first draw a random value for ∆*ε*_*RA*_ out of its prior distribution stated in Eq. (14) and calculate what the mean fold-change should be given our theory described in Eq. (7). With this in hand, we draw a random value for *σ* from its prior distribution, specified in Eq. (15). We then generate a simulated dataset by drawing ≈70 fold-change values across twelve inducer concentrations from the likelihood distribution which we defined in Eq. (12). This roughly matches the number of measurements made for each mutant in this work. We repeat this procedure for 800 draws from the prior distributions, which is enough to observe the occasional extreme fold-change value from the likelihood. As the DNA binding energy is the only parameter of our physical model that we are estimating, we had to choose values for the others. We kept the values of the inducer binding constants *K*_*A*_ and *K*_*I*_ the same as the wild-type repressor (139 *µ*M and 0.53 *µ*M, respectively). We chose to use *R* = 260 repressors per cell as this is the repressor copy number we used in the main text to estimate the DNA binding energies of the three mutants.

The draws from the priors are shown in S2(A) and (B) as black points above the corresponding distribution. To display the results, we computed the percentiles of the simulated data sets at each inducer concentration. These percentiles are shown as red shaded regions in Fig. S2(C). The 5th percentile (dark red band) has the characteristic profile of an induction curve. Given that the prior distribution for ∆*ε*_*RA*_ is centered at −12 *k*_*B*_*T* and we chose *R* = 260, we expect the generated data sets to cluster about the induction profile defined by these values. More importantly, approximately 95% of the generated data sets fall between fold-change values of −0.1 and 1.1, which is within the realm of possibility given the systematic and biological noise in our experiments. The 99^th^ percentile maximum is approximately 1.3 and the minimum approximately −0.3. While we could tune our choice of prior further to minimize draws this far from the theoretical bounds, we err on the side of caution and accept these values as it is possible that fold-change measurements this high or low can be observed, albeit rarely.

Through these prior predictive checks, we feel confident that these choices of priors are appropriate for the parameters we wish to estimate. We can now move forward and make sure that the statistical model as a whole is valid and computationally tractable.

#### Sensitivity Analysis and Simulation Based Calibration

Satisfied with our choice of prior distributions, we can proceed to check other properties of the statistical model and root out any pathologies lurking in our model assumptions.

To build trust in our model, we could generate a data set 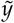 with a *known* value for *σ* and ∆*ε*_*RA*_, estimate the posterior distribution 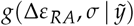, and determine how well we were able to retrieve the true value of the parameters. However, running this once or twice for handpicked values of *σ* and ∆*ε*_*RA*_ won’t reveal edge-cases in which the inference fails, some of which may exist in our data. Rather than performing this operation once, we can run this process over a variety of data sets where the ground truth parameter value is drawn from the prior distribution (as we did for the prior predictive checks). For an arbitrary parameter *θ*, the joint distribution between the ground truth value 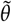, the inferred value *θ*, and the simulated data set 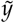 can be written as

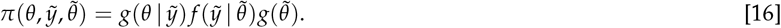

**Fig. S2.**
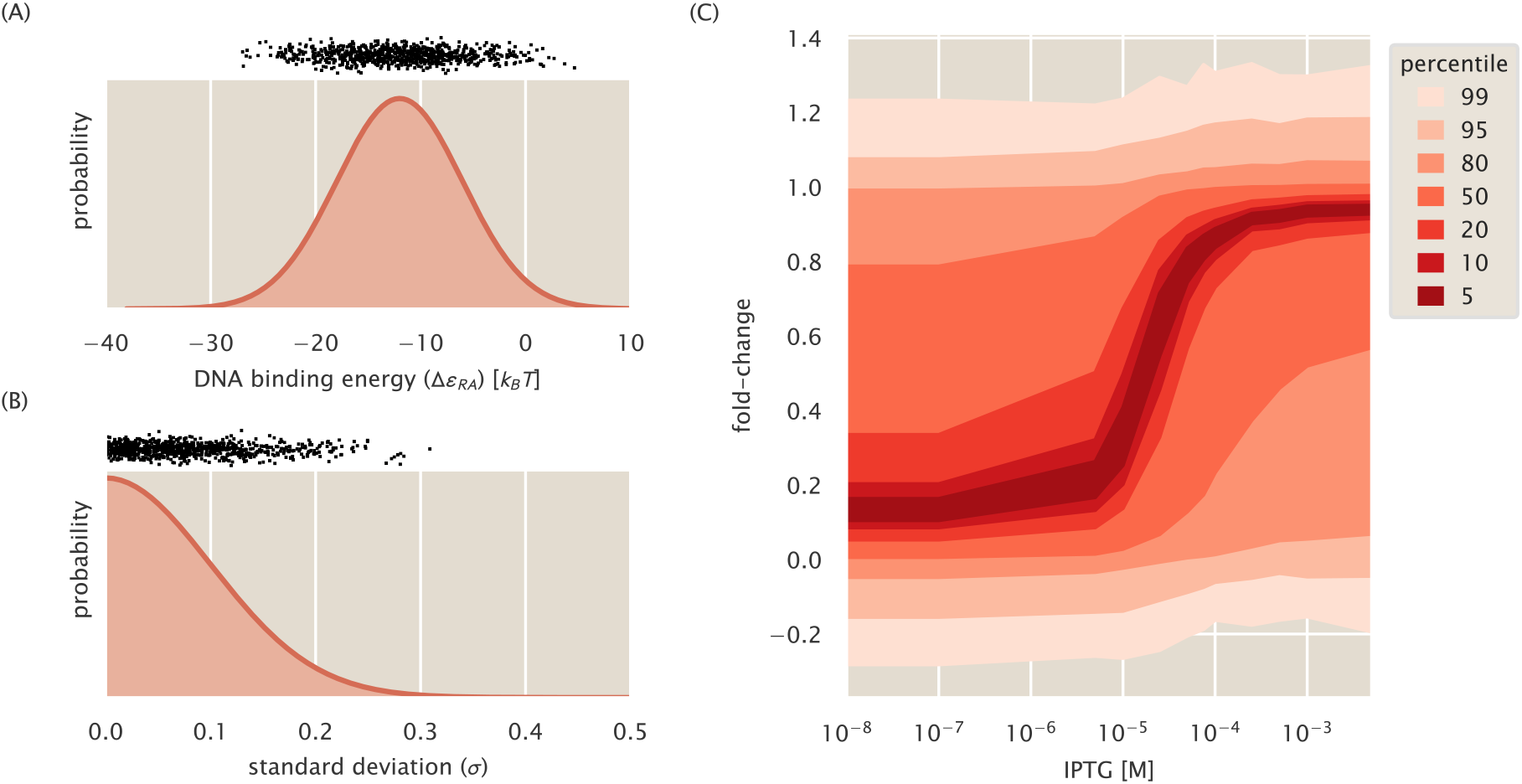
Prior distributions and prior predictive check for estimation of the DNA binding energy. (A) Prior probability density function for DNA binding energy ∆*ε*_*RA*_ as ~ Normal(−12, 12). (B) Prior probability density function for the standard deviation in measurement noise *σ* as ~ HalfNormal(0, 0.1). (C) Percentiles of values drawn from the likelihood distribution given draws from prior distributions given *R* = 260, *K*_*A*_ = 139 × 10^−6^ M, *K*_*I*_ = 0.53 × 10^−6^ M, and ∆*ε*_*AI*_ = 4.5 *k*_*B*_*T*, which match the parameters used for the predictions in Razo-Mejia et al. 2018 (2). Black points at top of (A) and (B) represent draws used to generate fold-change measurements from the likelihood distribution. Percentiles in (C) generated from 800 draws from the prior distributions. For each draw from the prior distributions, a data set of 70 measurements over 12 IPTG concentrations (ranging from 0 to 5000 *µ*M) were generated from the likelihood.

If this process is run for a large number of simulations, Eq. (16) can be marginalized over all data sets 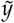 and all ground truth values 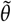 to yield the original prior distribution,

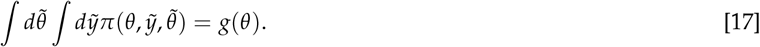

This result, described by Talts et al. 2018 (18), holds true for *any* statistical model and is a natural self consistency property of Bayesian inference. Any deviation between the distribution of our inferred values for *θ* and the original prior distribution *g*(*θ*) indicates that either our statistical model is malformed or the computational method is not behaving as expected. There are a variety of ways we can ensure that this condition is satisfied, which we outline below.

Using the data set generated for the prior predictive checks [shown in Fig. S2(C)], we sampled the posterior distribution and compute ∆*ε*_*RA*_ and *σ* for each simulation and checked that they matched the original prior distribution. To perform the inference, we use Markov chain Monte Carlo (MCMC) to sample the posterior distribution. Specifically, we use the Hamiltonian Monte Carlo algorithm implemented in the Stan probabilistic programming language (19). The specific code files can be accessed through the paper website or the associated GitHub repository. The original prior distribution and the distribution of inferred parameter values can be seen in Fig. S3 (A) and (B). For both ∆*ε*_*RA*_ and *σ*, we can accurately recover the ground truth distribution (blue) via sampling with MCMC (red). For ∆*ε*_*RA*_, there appears to be an upper and lower limit past which we are unable to accurately infer the binding energy. This can be seen in both the histogram [Fig. S3(A)] and the empirical cumulative distribution [Fig. S3(B)] as deviations from the ground truth when DNA binding is below ≈ −25*k*_*B*_*T* or above ≈ −5*k*_*B*_*T*. These limits hinder our ability to comment on exceptionally strong or weak binding affinities. However, as all mutants queried in this work exhibited binding energies between these limits, we surmise that the inferential scheme permits us to draw conclusions about the inferred DNA binding strengths.

Rather than examining the agreement of the data-averaged posterior and the ground truth prior distribution solely by eye, we can compute summary statistics using the mean *µ* and standard deviation *σ* of the posterior and prior distributions which permit easier identification of pathologies in the inference. One such quantity is the posterior *z*-score, which is defined as

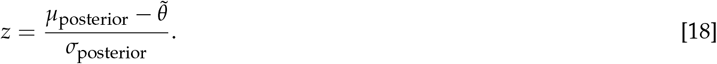

This statistic summarizes how accurately the posterior recovers the ground truth value beyond simply reporting the mean, median, or mode of the posterior distribution. *Z*-scores around 0 indicate that the posterior is concentrating tightly about the true value of the parameter whereas large values (either positive or negative) indicate that the posterior is concentrating elsewhere. A useful feature of this metric is that the width of the posterior is also considered. It is possible that the posterior could have a mean very close to the ground truth value, but have an incredibly narrow distribution/spread such that it does not overlap with the ground-truth. Only comparing the mean value to the ground truth would suggest that the inference “worked”. However with a small standard deviation generates a very large *z*-score, telling us that something has gone awry.

**Fig. S3.**
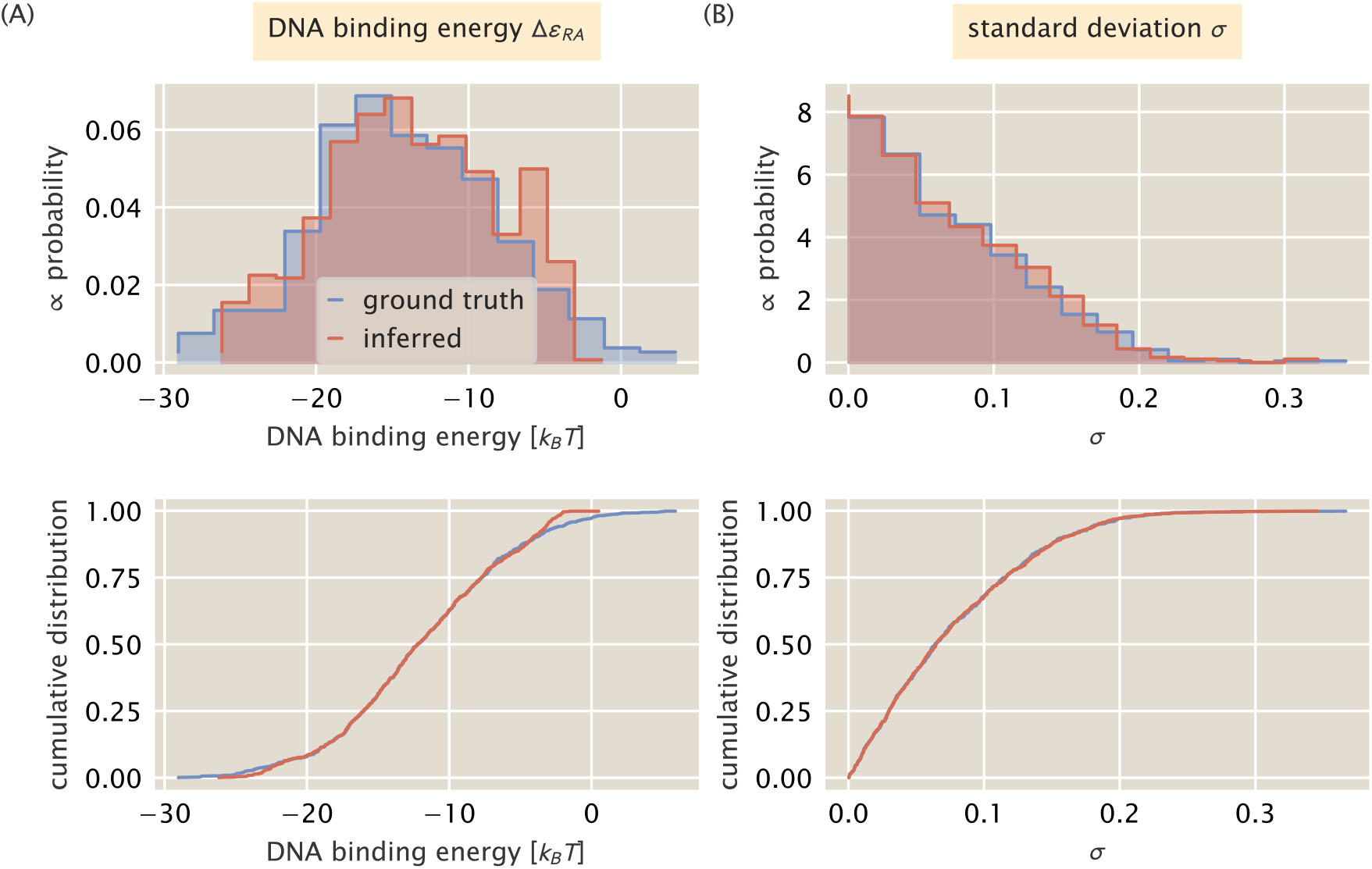
Comparison of averaged posterior and prior distributions for ∆*ε*_*RA*_ and *σ*. (A) Distribution of the average values for the DNA binding energy ∆*ε*_*RA*_(red) overlaid with the ground truth distribution (blue). (B) Data averaged posterior (red) for the standard deviation of fold-change measurements overlaid with the ground truth distribution (blue). Top and bottom show the same data with different visualizations.

If our inferential model is behaving properly, the width of the posterior distribution should be significantly smaller than the width of the prior, meaning that the posterior is being informed by the data. The level to which the posterior is being informed by the data can be easily calculated given knowledge of both the prior and posterior distribution. This quantity, aptly named the shrinkage *s*, can be computed as

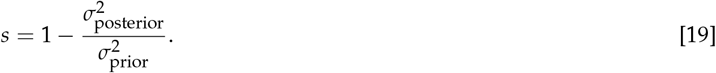

When the shrinkage is close to zero, the variance of the posterior is approximately the same as the variance of the prior, model is not being properly informed by the data. When *s* ≈ 1, the variance of the posterior is much smaller than the variance of the prior, indicating that the it is being highly informed by the data. A shrinkage less than 0 indicates that the posterior is wider than the prior distribution, revealing a severe pathology in either the model itself or the implementation.

In Fig. S4, we compute these summary statistics for each parameter. For both ∆*ε*_*RA*_ and *σ*, we see clustering of the *z*-score about 0 with the extrema reaching ≈ ±3. This suggests that for the vast majority of our simulated data sets, the posterior distribution concentrated about the ground truth value. We also see that for both parameters, the posterior shrinkage *s* is ≈ 1, indicating that the posterior is being highly informed by the data. There is a second distribution centered ≈ 0.8 for ∆*ε*_*RA*_, indicating that for a subset of the data sets, the posterior is only ≈ 80% narrower than the prior distribution. These samples are those that were drawn outside of the limits of ≈ −25 to − 5 *k*_*B*_*T* where the inferential power is limited. Nevertheless, the posterior still significantly shrank, indicating that the data strongly informs the posterior.

The general self-consistency condition given by Eq. (17) provides another route to ensure that the model is computationally tractable. Say that we draw a value for the DNA binding energy from the prior distribution, simulate a data set, and sample the posterior using MCMC. The result of this sampling is a collection of *N* values of the parameter which may be above, below, or equal to the ground-truth value. From this set of values, we select *L* of them and rank order them by their value. Talts and colleagues (18) derived a general theorem which states that the number of samples less than the ground truth value of the parameter (termed the rank statistic) is uniformly distributed over the interval [0, *L*]. As Eq. (17) *must* hold true for any statistical model, deviations from uniformity signal that there is a problem in the implementation of the statistical model. How the distribution deviates is also informative as different types of failures result in different distributions. The nature of these deviations, along with a more formal proof of the uniform distribution of rank statistics can be found in Talts et al. 2018 (18) where it was originally derived.

Given the sampling statistics for each of the simulated data sets, we took 800 of the MCMC samples of the posterior distribution for each of the 800 simulated data sets and computed the rank statistic. The distributions are shown in Fig. S5 as both histograms and ECDFs for the DNA binding energy and standard deviation. The distribution of rank statistics for both parameters appears to be uniform. The gray band overlaying the histograms (top row) as well as the gray envelopes overlaying the ECDFs (bottom row) represent the 99^th^ percentile expected from a true uniform distribution. The uniformity of this distribution, along with the well-behaved *z*-scores and shrinkage for each parameter, tells us that there are no underlying pathologies in our statistical model and that it is computationally tractable. However, this does not mean that it is correct. Whether this model is valid for the actual observed data is the topic of the next section.

**Fig. S4.**
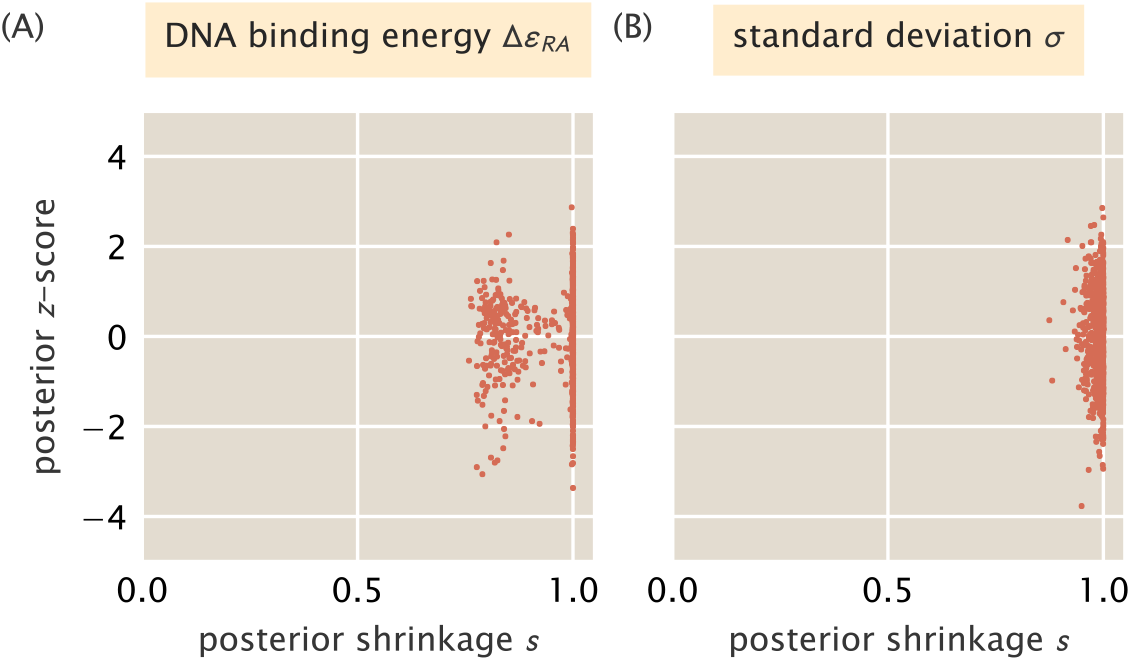
Inferential sensitivity for estimation of ∆*ε*_*RA*_ and *σ*. The posterior *z*-score for each posterior distribution inferred from a simulated data set is plotted against the shrinkage for (A) the DNA binding energy ∆*ε*_*RA*_ and (B) the standard deviation of fold-change measurements *σ*

#### Parameter Estimation and Posterior Predictive Checks

We now turn to applying our vetted statistical model to experimental measurements. While the same statistical model was applied to all three DNA binding mutants, here we only focus on the mutant Q21M for brevity.

Using a single induction profile, we sampled the posterior distribution over both the DNA binding energy ∆*ε*_*RA*_ and the standard deviation *σ* using MCMC implemented in the Stan programming language. The output of this process is a set of 4000 samples of both parameters along with the value of their log posterior probabilities, which serves as an approximate measure of the probability of each value. The individual samples are shown in Fig. S6. The joint distribution between ∆*ε*_*RA*_ and *σ* is shown in the lower left hand corner, and the marginal distributions for each parameter are shown above and to the right of the joint distribution, respectively. The joint distribution is color coded by the value of the log posterior, with yellow and blue corresponding to high and low probability, respectively. The symmetric shape of the joint distribution is a telling sign that there is no correlation between two parameters. The marginal distributions for each parameter are also relatively narrow, with the DNA binding energy covering a range of ≈ 0.6 *k*_*B*_*T* and *σ* spanning ≈ 0.02. To more precisely quantify the uncertainty, we computed the shortest interval of the marginal distribution for each parameter contains 95% of the probability. The bounds of this interval, coined the Bayesian credible region, can accommodate asymmetry in the marginal distribution since the upper and lower bounds of the estimate are reported. In the main text, we reported the DNA binding energy estimated from these data to be 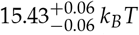, where the first value is the median of the distribution and the super- and subscripts correspond to the upper and lower bounds of the credible region, respectively.

While looking at the shape of the posterior distribution can be illuminating, it is not enough to tell us if the parameter values extracted make sense or accurately describe the data on which they were conditioned. To assess the validity of the statistical model in describing actual data, we again turn to simulation, this time using the posterior distributions for each parameter rather than the prior distributions. The likelihood of our statistical model assumes that across the entire induction profile, the observed fold-change is normally distributed about the theoretical prediction with a standard deviation *σ*. If this is an accurate depiction of the generative process, we should be able to draw values from the likelihood using the sampled values for ∆*ε*_*RA*_ and *σ* that are indistinguishable from the actual experimental measurements. This process is known as a *posterior predictive check* and is a Bayesian method of assessing goodness-of-fit.

For each sample from the posterior, we computed the theoretical mean fold-change given the sampled value for ∆*ε*_*RA*_ and Eq. (7). With this mean in hand, we used the corresponding sample for *σ* and drew a data set from the likelihood distribution the same size as the real data set used for the inference. As we did this for every sample of our MCMC output (a total of ≈ 4000), it is more instructive to compute the percentiles of the generated data than to show the entire output. In Fig. S6(B), the percentiles of the generated data sets are shown overlaid with the data used for the inference. We see that all of the data points fall within the 99^th^ percentile of simulated data sets with the 5^th^ percentile tracking the mean of the data at each inducer concentration. As there are no systematic deviations or experimental observations that fall far outside those generated from the statistical model, we can safely say that the statistical model derived here accurately describes the observed data.

**Fig. S5.**
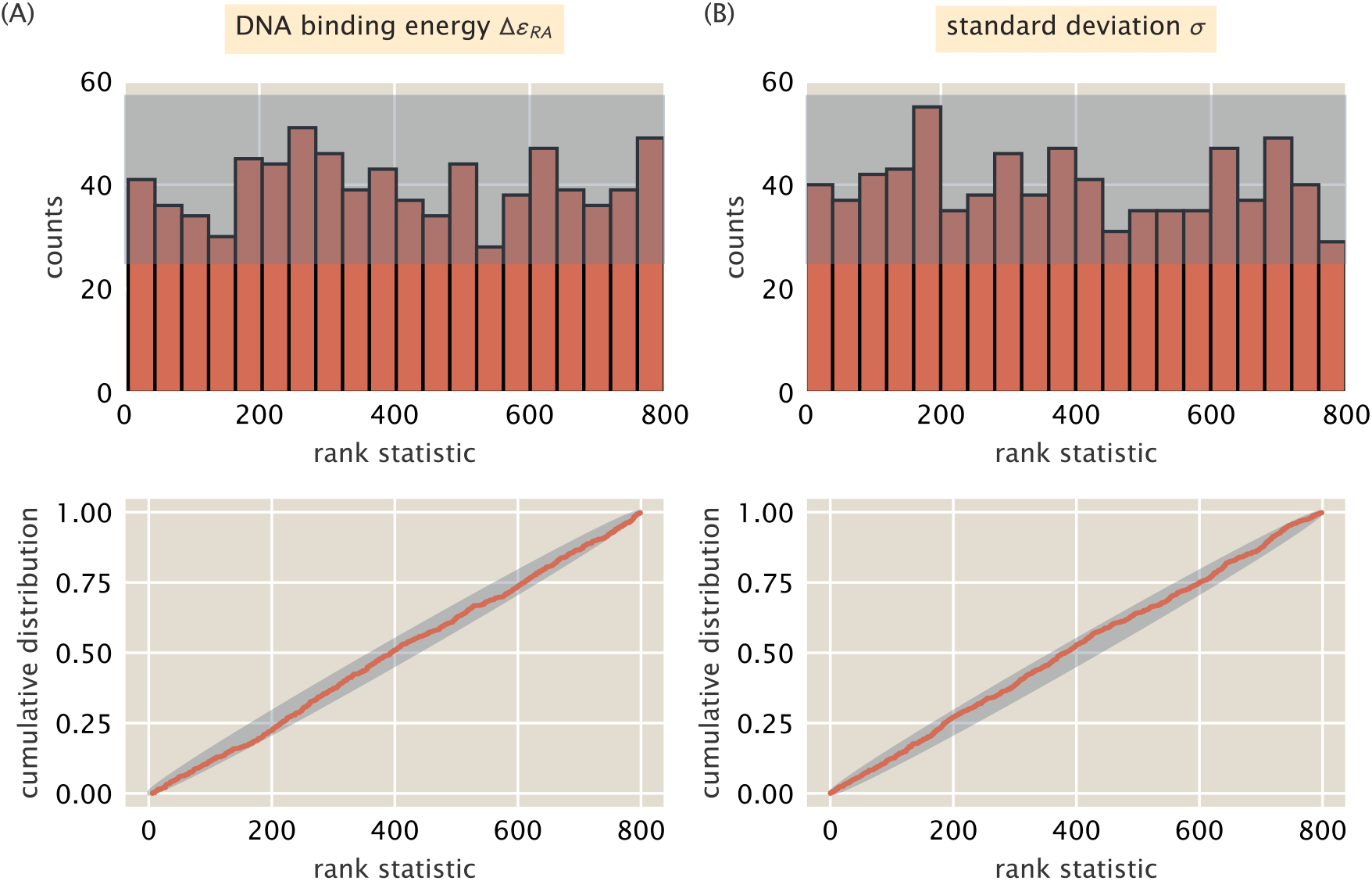
Rank distribution of the posterior samples from simulated data. Top row shows a histogram of the rank distribution with *n* = 20 bins. Bottom row is the cumulative distribution for the same data. Gray bands correspond to the 99th percentile of expected variation from a uniform distribution. (A) Distribution for the DNA binding energy ∆*ε*_*RA*_ and (B) for the standard deviation *σ*.

**Fig. S6.**
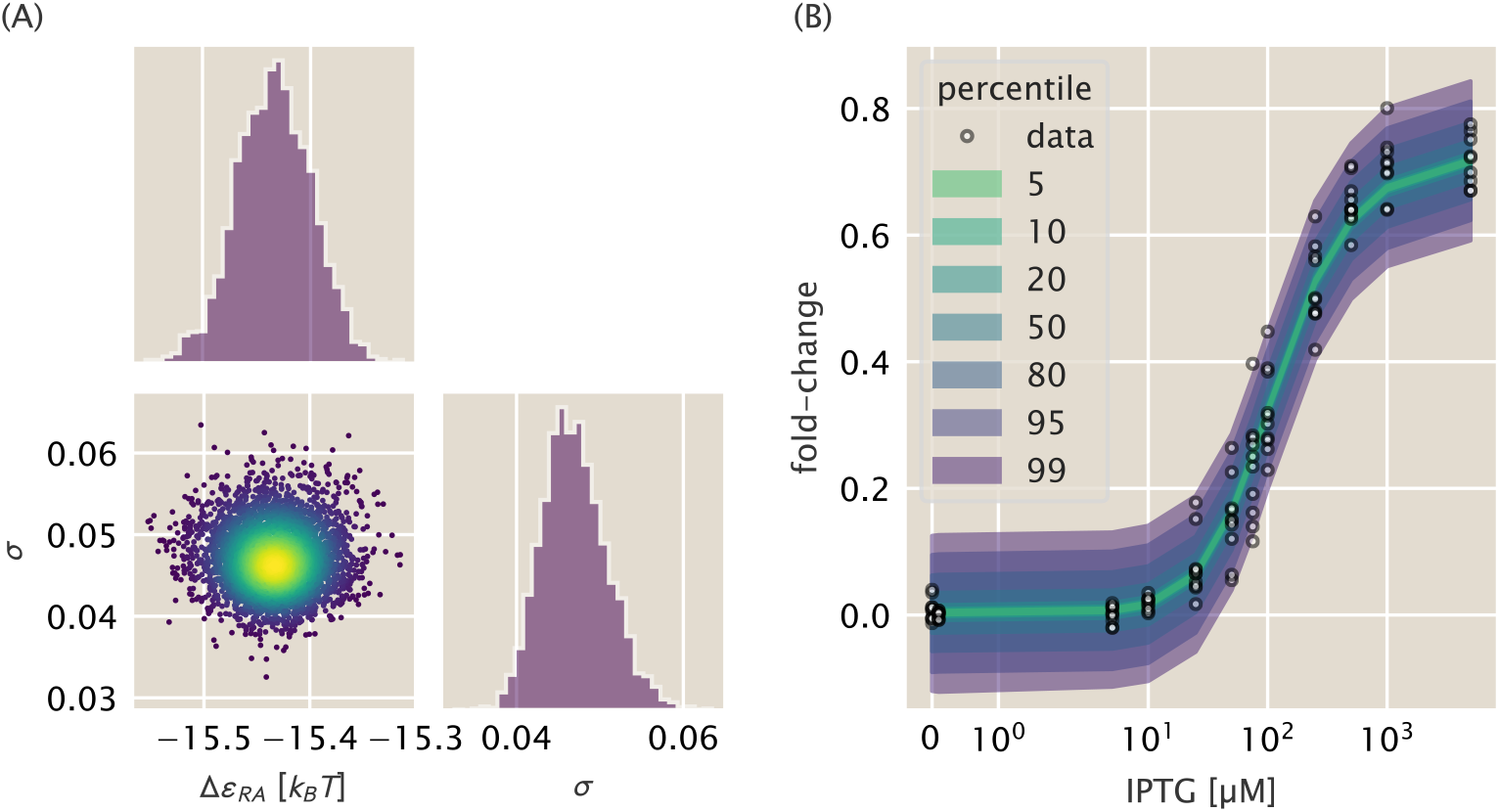
Markov Chain Monte Carlo (MCMC) samples and posterior predictive check for DNA binding mutant Q21M. (A) Marginal and joint sampling distributions for DNA binding energy ∆*ε*_*RA*_ and *σ*. Each point in the joint distribution is a single sample. Marginal distributions for each parameter are shown adjacent to joint distribution. Color in the joint distribution corresponds to the value of the log posterior with the progression of blue to yellow corresponding to increasing probability. (B) The posterior predictive check of model. The measurements of the fold-change in gene expression are shown as black open-faced circles. The percentiles are shown as colored bands and indicate the fraction of simulated data drawn from the likelihood that fall within the shaded region.

### 3. Inferring the Free Energy From Fold-Change Measurements

In this section, we describe the statistical model to infer the free energy *F* from a set of fold-change measurements. We follow the same principled workflow as described previously for the DNA binding estimation, including declaration of the generative model, prior predictive checks, simulation based calibration, and posterior predictive checks. Finally, we determine an empirical limit in our ability to infer the free energy and define a heuristic which can be used to identify measurements that are likely inaccurate. To understand the statistical model and the empirical limits of detection, only the subsections *Building A Generative Model* and *Sensitivity Limits and Systematic Errors in Inference* are necessary.

#### Building A Generative Model

In the main text, we showed that the fold-change equation defined in Eq. (7) can be rewritten in the form of a Fermi function,

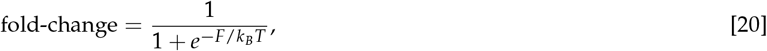

where *F* corresponds to the free energy difference between the repressor bound and unbound states of the promoter. While the theory prescribes a way for us to calculate the free energy based on our knowledge of the biophysical parameters, we can directly calculate the free energy of a measurement of fold-change by simply rearranging Eq. (20) as

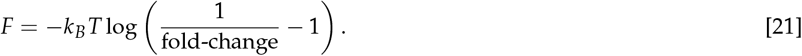

With perfect measurement of the fold-change in gene expression (assuming no experimental or measurement noise), the free energy can be directly calculated. However, actual measurements of the fold-change in gene expression can extend beyond the theoretical bounds of 0 and 1, for which the free energy is mathematically undefined.

As the fold-change measurements between biological replicates are independent, it is reasonable to assume that they are normally distributed about a mean value *μ* with a standard deviation *σ*. While the mean value is restricted to the bounds of [0, 1], fold-change measurements outside of these bounds are still possible given that they are distributed about the mean with a scale of *σ*. Thus, if we have knowledge of the mean fold-change in gene expression about which the observed fold-change is distributed, we can calculate the mean free energy as

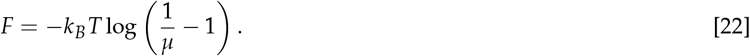

For a given set of fold-change measurements *y*, we wish to infer the posterior probability distribution for *μ* and *σ*, given by Bayes’ theorem as

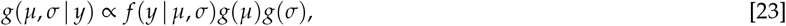

where we have dropped the normalization constant *f* (*y*) and assigned a proportionality between the posterior and joint probability distribution. Given that the measurements are independent, we define the likelihood *f* (*y* | *μ*, *σ*) as a normal distribution,

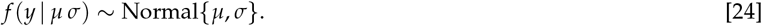

While the mean *μ* is restricted to the interval [0, 1], there is no reason *a priori* to think that it is more likely to be closer to either bound. To remain uninformative and be as permissive as possible, we define a prior distribution for *μ* as a Uniform distribution between 0 and 1,

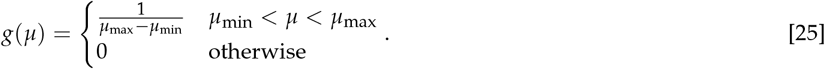

Here, *μ*_min_ = 0 and *μ*_max_ = 1, reducing *g*(*μ*) to 1. For *σ*, we can again assume a half-normal distribution with a standard deviation of 0.1 as was used for estimating the DNA binding energy [Eq. (15)],

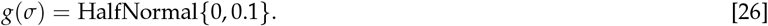

With a full generative model defined, we can now use prior predictive checks to ensure that our choices of prior are appropriate for the inference.

#### Prior Predictive Checks

To check the validity of the chosen priors, we pulled 1000 combinations of *μ* and *σ* from their respective distributions [Fig. S7(A)] and subsequently drew a set of 10 fold-change values (a number comparable to the number of biological replicates used in this work) from a normal distribution defined by *μ* and *σ*. To visualize the range of values generated from these checks, we computed the percentiles of the empirical cumulative distributions of the fold-change values, as can be seen in Fig. S7(C). Approximately 95% of the the generated fold-change measurements were between the theoretical bounds of [0, 1] whereas 5% of the data sets fell outside with the maximum and minimum values extending to ≈ 1.2 and −0.2, respectively. Given our familiarity with these experimental strains and the detection sensitivity of the flow cytometer, these excursions beyond the theoretical bounds agree with our intuition. Satisfied with our choice of prior distributions, we can proceed to check the sensitivity and computational tractability of our model through simulation based calibration.

**Fig. S7.**
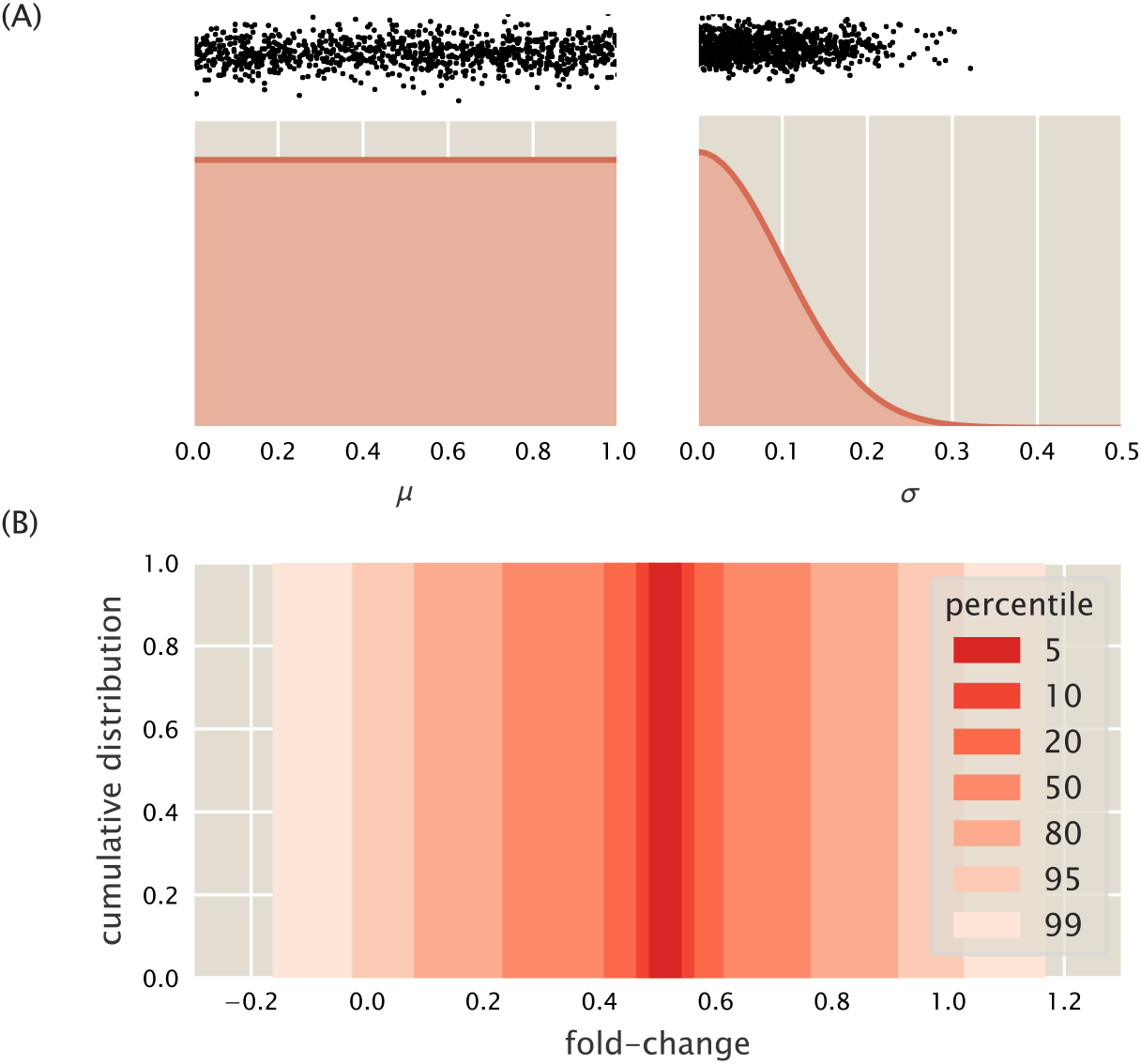
Prior predictive checks for inference of the mean fold-change. (A) The prior distributions for *μ* (left) and *σ* (right). The vertical axis is proportional to the probability of the value. Black points above distributions correspond to the values used to perform the prior predictive checks. (B) Percentiles of the data generated for each draw from the prior distributions shown as a cumulative distribution. Percentiles were calculated for 1000 generated data sets, each with 10 fold-change measurements drawn from the likelihood given the drawn values of *μ* and *σ*.

#### Simulation Based Calibration

To ensure that the parameters can be estimated with confidence, we sampled the posterior distribution of *μ* and *σ* for each data set generated from the prior predictive checks. For each inference, we computed the *z*-score and shrinkage for each parameter, shown in Fig. S8(A). For both parameters, the *z*-scores are approximately centered about zero, indicating that the posteriors concentrate about the ground truth value of the parameter. The *z*-scores for *σ* [black points in Fig. S10(A)] appear to be slightly off centered with more negative values than positive. This suggests that *σ* is more likely to be slightly overestimated in some cases. The shrinkage parameter for *μ* (red points) is very tightly distributed about 1.0, indicating that the prior is being strongly informed by the data. The shrinkage is more broadly distributed for for *σ* with a minimum value of ≈ 0.5. However, the median shrinkage for *σ* is ≈ 0.9, indicating that half of the inferences shrank the prior distribution by at least 90%. While we could revisit the model to try and improve the shrinkage values, we are more concerned with *μ* which shows high shrinkage and zero-centered *z*-scores.

To ensure that the model is computationally tractable, we computed the rank statistic of each parameter for each inference. The empirical cumulative distributions for *μ* (black) and *σ* (red) can be seen in Fig. S8(B). Both distributions appear to be uniform, falling within the 99^th^ percentile of the variation expected from a true uniform distribution. This indicates that the self-consistency relation defined by Eq. (17) holds for this statistical model. With a computationally tractable model in hand, we can now apply the statistical model to our data and verify that data sets drawn from the data-conditioned posterior are indistinguishable from the experimental measurements.

#### Posterior Predictive Checks

The same statistical model was applied to every unique set of fold-change measurements used in this work. Here, we focus only on the set of fold-change measurements for the double mutant Y20I-Q294V at 50 *μ*M IPTG. The samples from the posterior distribution conditioned on this dataset can be seen in Fig. S9(A). The joint distribution, shown in the lower left-hand corner, appears fairly symmetric, indicating that *μ* and *σ* are independent. There is a slight asymmetry in the sampling of *σ*, which can be more clearly seen in the corresponding marginal distribution to the right of the joint distribution.

For each MCMC sample of *μ* and *σ*, we drew 10 samples from a normal distribution defined by these parameters. From this collection of data sets, we computed the percentiles of the empirical cumulative distribution and plotted them over the data, as can be seen in Fig. S9 (B). We find that the observed data falls within the 99^th^ percentile of the generated data sets. This illustrates that the model can produce data which is identically distributed to the actual experimental measurements, validating our choice of statistical model.

#### Sensitivity Limits and Systematic Errors in Inference

Considering the results from the prior predictive checks, simulation based calibration, and posterior predictive checks, we can say that the statistical model for inferring *μ* and *σ* fold-change from a collection of noisy fold-change measurements is valid and computationally tractable. Upon applying this model to the experimental data of the wild-type strain (where the free energy is theoretically known), we observed that systematic errors arise when the fold-change is exceptionally high or low, making the resulting inference of the free energy inaccurate.

**Fig. S8.**
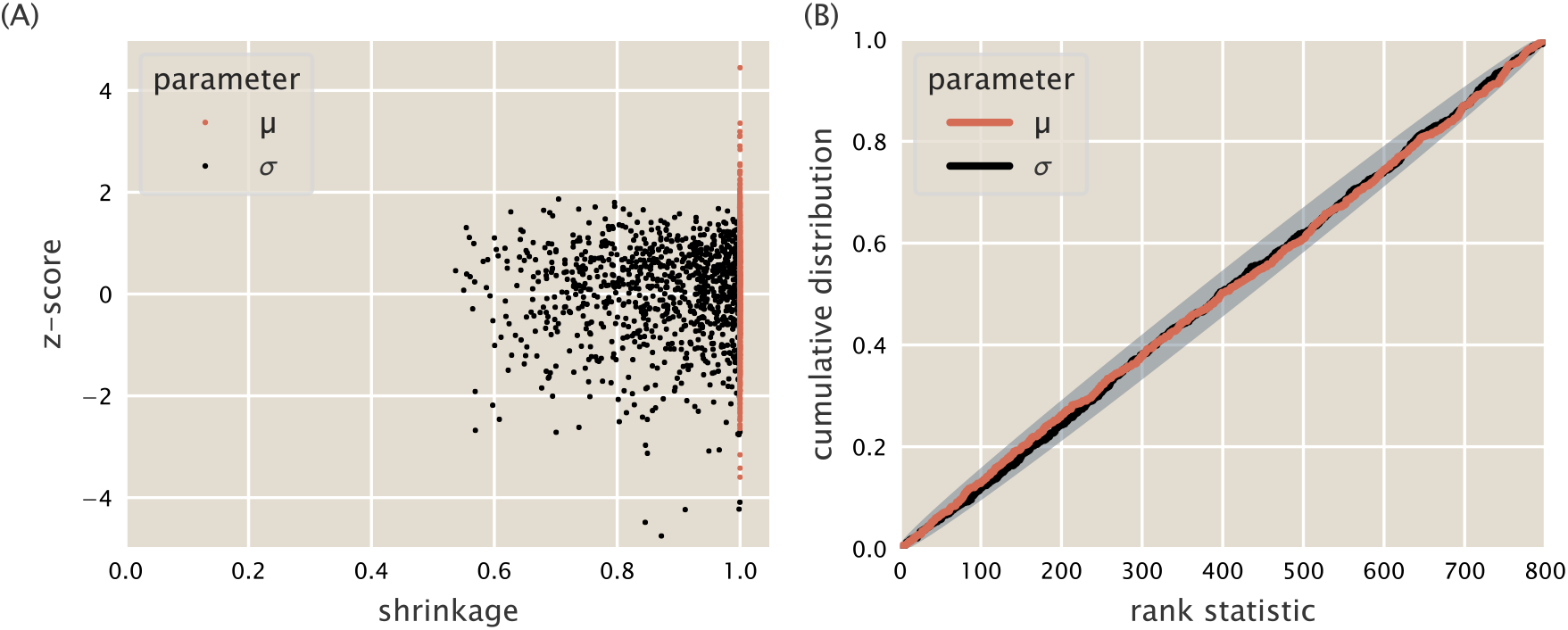
Sensitivity measurements and rank statistic distribution of the statistical model estimating *μ* and *σ*. (A) Posterior *z*-score of each inference plotted against the posterior shrinkage factor for the parameters *μ* (red points) and *σ* (black points). (B) Distribution of rank statistics for *μ* (red) and *σ* (black). Gray envelope represents the 99^th^ percentile of a true uniform distribution.

To elucidate the source of this systematic error, we return to a simulation based approach in which the true free energy is known [black points in Fig. S10(A)]. For a range of free energies, we computed the theoretical fold-change prescribed by Eq. (20). For each free energy value, we pulled a value for *σ* from the prior distribution defined in Eq. (15) and generated a data set of 10 measurements by drawing values from a normal distribution defined by the true fold-change and the drawn value of *σ* [red points in Fig. S10(A)]. We then sampled the statistical model over these data and inferred the mean fold-change *μ* [blue points in Fig. S10(A)]. By eye, the inferred points appear to collapse onto the master curve, in many cases overlapping the true values. However, the points with a free energy less than ≈ −2 *k*_*B*_*T* and greater than ≈ 2 *k*_*B*_*T* are slightly above or below the master curve, respectively. This becomes more obvious when the inferred free energy is plotted as a function of the true free energy, shown in Fig. S10(B). Points in which the difference between *μ* and the neearest boundary (0 or 1) is less than the value of *σ* are shown as purple or green. When this condition is met, the inferred mean free energy strays from the true value, introducing a systematic error. This suggests that the spread of the fold-change measurements sets the detection limit of fold-change close to either boundary. Thus, the narrower the spread in the fold-change the better the estimate of the fold-change near the boundaries.

These systematic errors can be seen in experimental measurements of the wild-type repressor. Data from Razo-Mejia et al. 2018(2) in which the IPTG titration profiles of seventeen different bacterial strains were measured is shown collapsed onto the master curve in Fig. S10(C) as red points. Here, each point corresponds to a single biological replicate. The inferred mean fold-change *μ* and 95% credible regions are shown as purple, blue, or green points. The color of these points correspond to the relative value of *μ* or 1 − *μ* to *σ*. The discrepancy between the predicted and inferred free energy of each measurement set can be seen in Fig. S10(D). The significant deviation from the predicted and inferred free energy occurs past the detection limit set by *σ*. In this work, we therefore opted to not display inferred free energies at the extrema where the inferred fold-change was closer to the boundaries than the correspoding standard deviation, as it reflects limitations in our measurement rather than a deviation from the theoretical predictions.

**Fig. S9.**
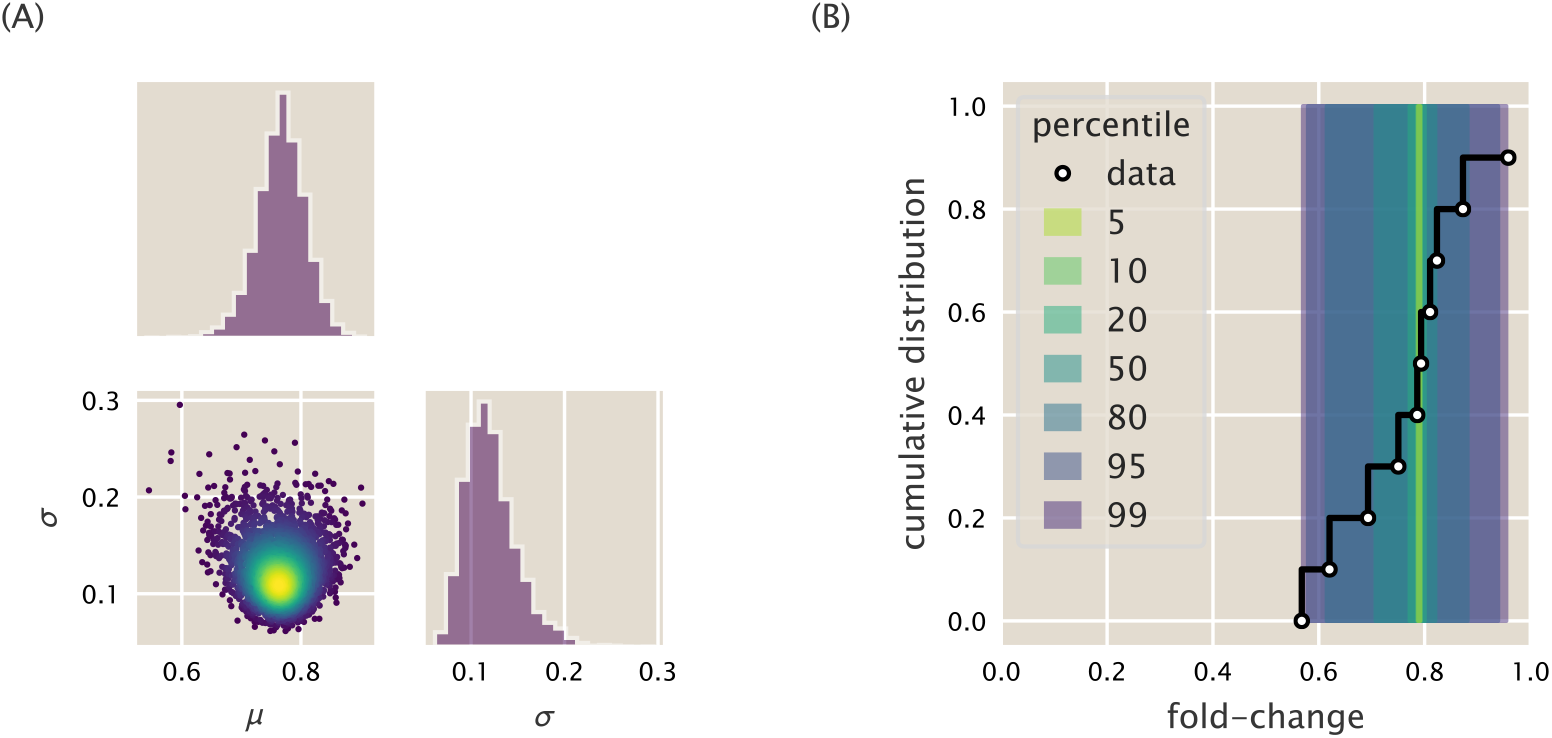
MCMC sampling output and posterior predictive checks of the statistical model for the mean fold-change *μ* and standard deviation *σ*. (A) Corner plot of sampling output. The joint distribution between *σ* and *μ* is shown in the lower left hand corner. Each point is an individual sample. Points are colored by the value of the log posterior with increasing probability corresponding to transitions from blue to yellow. Marginal distributions for each parameter are shown adjacent to the joint distribution. (B) Percentiles of the cumulative distributions from the posterior predictive checks are shown as shaded bars. Data on which the posterior was conditioned are shown as white circles connected by black lines.

**Fig. S10.**
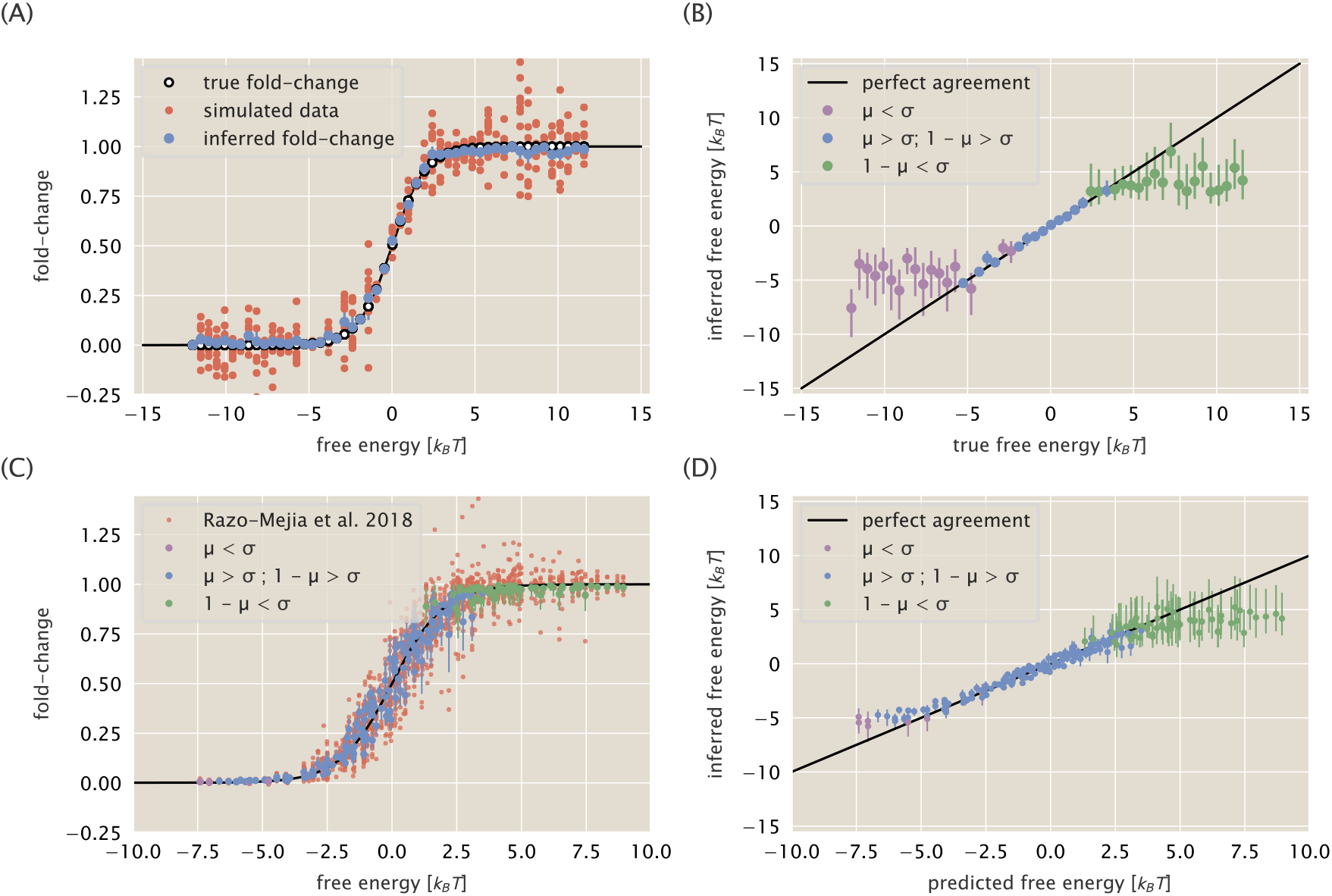
Identification of systematic error in simulated and real data when considering the free energy. (A) The true fold-change (black open circles), simulated fold-change distribution (red points), and inferred mean fold-change (blue) is plotted as a function of the true free energy. Error bars on inferred fold-change correspond to the 95% credible region of the mean fold-change *μ*. (B) Inferred free energy plotted as a function of the true free energy. Black line indicates perfect agreement between the ground truth free energy and inferred free energy. Blue points correspond to the inferred free energy where the median values of the parameters satisfy the condition *μ > σ* and 1 − *μ > σ*. Purple points correspond to the inferred mean fold-change *μ < σ*. Green points correspond to those where the inferred mean fold-change 1 −*μ < σ*. Error bars correspond to the bounds of the 95% credible region. (C) Biological replicate data from Razo-Mejia et al. 2018 (2) (red points) plotted as a function of the theoretical free energy. Inferred mean fold-change *μ* and the 95% credible region are shown as blue points. Purple and green points are colored by the same conditions as in (B). (D) Inferred free energy as a function of the predicted free energy colored by the satisfied condition. Error bars are the bounds of the 95% credible region. All inferred values in (A - D) are the median values of the posterior distribution.

### 4. Additional Characterization of DNA Binding Mutants

In the main text, we estimated the DNA binding energy o f each mutant using the mutant strains that had approximately 260 repressors per cell. In this section, we examine the effect of the choice of fit strain on the predictions of both the induction profiles and Δ*F* for each DNA binding domain mutant.

We applied the statistical model derived in Section 2 for each unique strain of the DNA binding mutants and estimated the DNA binding energy. The median of the posterior distribution along with the upper and lower bounds of the 95% credible region are reported in Table S1. We found that the choice of fitting strain did not strongly influence the estimate of the DNA binding energy. The largest deviations appear for the weakest binding mutants paired with the lowest repressor copy number. In these cases, such as for Q21A, the difference in binding energy between the repressor copy numbers is ≈ 1 *k*_*B*_*T* which is small compared to the overall DNA binding energy. Using these energies, we computed the predicted induction profiles of each mutant with different repressor copy numbers, shown in Fig. S11. In this plot, the rows correspond to the repressor copy number of the strain used to estimate the DNA binding energy. The columns correspond to the repressor copy number of the predicted strains. The diagonals, shaded in grey, show the induction profile of the fit strain along with the corresponding data. In all cases, we find that the predicted profiles are relatively accurate with the largest deviations resulting from using the lowest repressor copy number as the fit strain.

The predicted change in free energy Δ*F* using each fit strain can be seen in Fig. S12. In this figure, the rows represent the repressor copy number of the strain to which the DNA binding energy was fit whereas the columns correspond to each mutant. In each plot, we have shown the data for all repressor copy numbers with the fit strain represented by white filled circles. Much as for the induction profiles, we see little difference in the predicted Δ*F* for each strain, all of which accurately describe the inferred free energies. The ability to accurately predict the majority of the induction profiles of each mutant with repressor copy numbers ranging over two orders of magnitude strengthens our assessment that for these DNA binding domain mutations, only the DNA binding energy is modified.

**Fig. S11.**
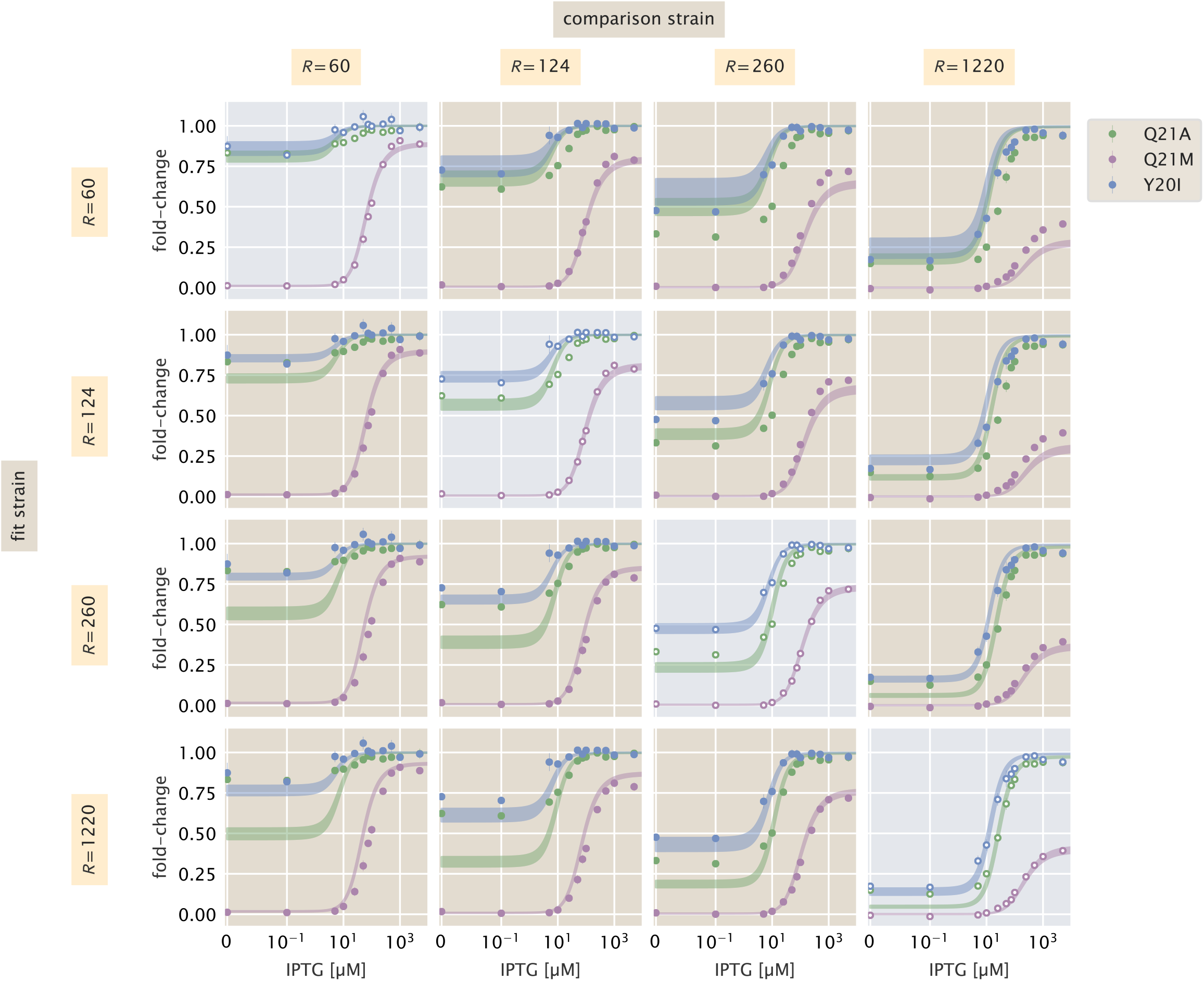
Pairwise comparisons of DNA binding mutant induction profiles. Rows correspond to the repressor copy number of the strain used to estimate the DNA binding energy for each mutant. Columns correspond to the repressor copy number of the strains that are predicted. Diagonals in which the data used to estimate the DNA binding energy are shown with a gray background.

**Fig. S12.**
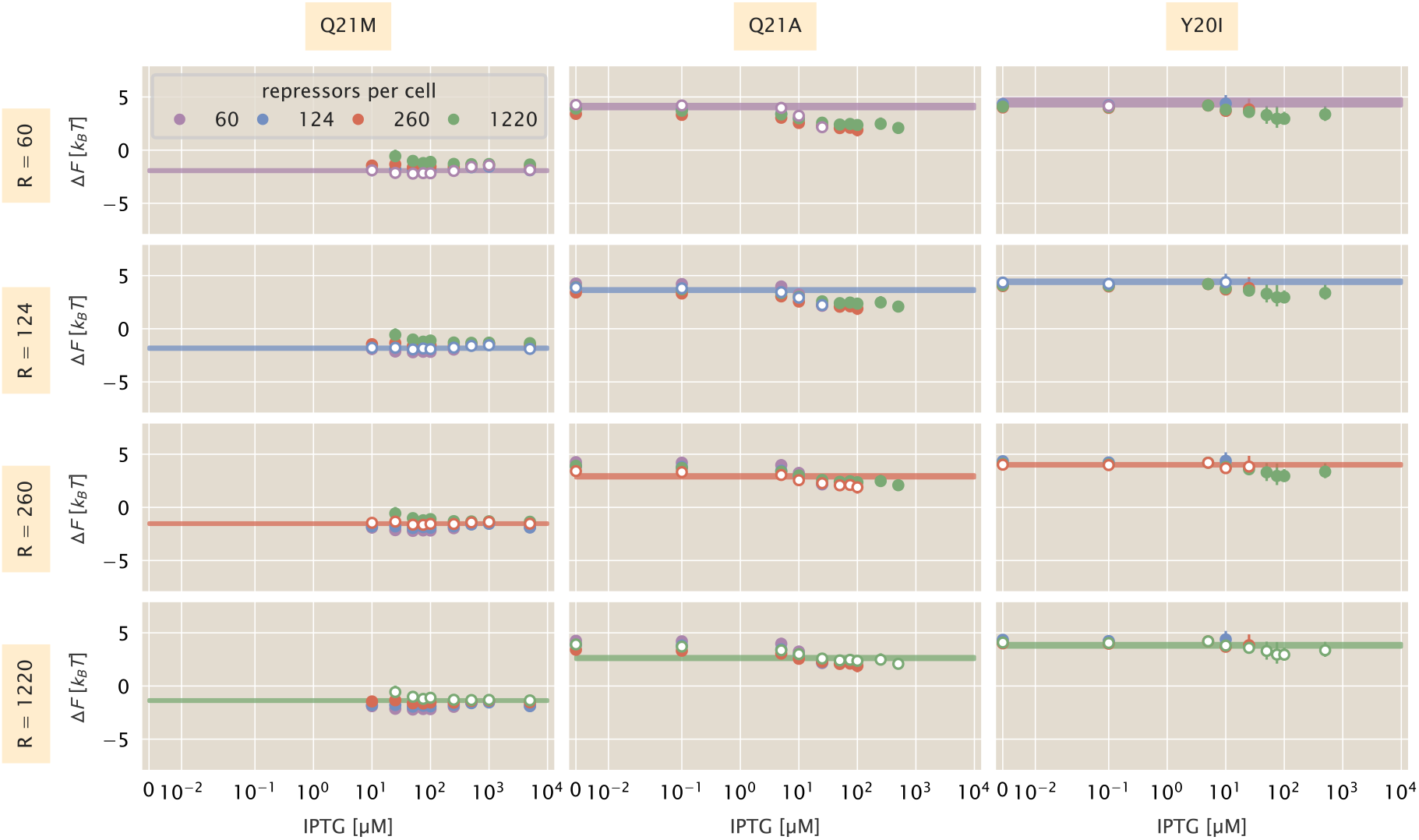
Dependence of fitting strain on Δ*F* predictions of DNA binding domain mutants. Rows correspond to the repressor copy number used to estimate the DNA binding energy. Columns correspond to the particular mutant. Colored lines are the bounds of the 95% credible region of the predicted Δ*F*. Open face points indicate the strain to which the DNA binding energy was fit.

**Table S1.** Estimated DNA binding energy for DNA binding domain mutants with different repressor copy numbers. Median of the posterior distribution with the upper and lower bounds of the 95% credible region are reported.

| Mutant | Repressors | DNA Binding Energy [$k_B T$] |
| --- | --- | --- |
| Q21A | 60 | $-9.8^{+0.2}_{-0.2}$ |
| | 124 | $-10.3^{+0.1}_{-0.1}$ |
| | 260 | $-11.0^{+0.1}_{-0.1}$ |
| | 1220 | $-11.3^{+0.1}_{-0.1}$ |
| Q21M | 60 | $-15.83^{+0.08}_{-0.08}$ |
| | 124 | $-15.7^{+0.1}_{-0.1}$ |
| | 260 | $-15.43^{+0.07}_{-0.06}$ |
| | 1220 | $-15.27^{+0.07}_{-0.07}$ |
| Y20I | 60 | $-9.4^{+0.3}_{-0.3}$ |
| | 124 | $-9.5^{+0.1}_{-0.1}$ |
| | 260 | $-9.9^{+0.1}_{-0.1}$ |
| | 1220 | $-10.1^{+0.2}_{-0.2}$ |

### 5. Bayesian Parameter Estimation for Inducer Binding Domain Mutants

In the main text, we put forward two naïve hypotheses for which parameters of Eq. (7) are affected by mutations in the inducer binding domain of the repressor. The first hypothesis was that only the inducer dissociation constants, *K*_*A*_ and *K*_*I*_, were perturbed from their wild-type values. Another hypothesis was that the inducer dissociation constants were affected in addition to the energetic difference between the active and inactive states of the repressor, Δ*ε* _*AI*_.

In this section, we first derive the statistical model for each hypothesis and then perform a series of diagnostic tests that expose the inferential limitations of each model. With well calibrated statistical models, we then apply each to an induction profile of the inducer binding mutant Q294K and assess the validity of each hypothesis. To understand the statistical models for each hypothesis, only the subsection *Building A Generative Statistical Model* is necessary.

#### Building a Generative Statistical Model

For both hypotheses, we assume that the underlying physical model defined in Eq. (7) is the same while a subset of the parameters are modified. As the fold-change measurements for each biological replicate are statistically independent, we can assume that they are normally distributed about the theoretical fold-change value. Thus, for each model, we must include a parameter *σ* which is the standard deviation of the distribution of fold-change measurements. For the first hypothesis, in which only *K*_*A*_ and *K*_*I*_ are changed, we are interested in sampling the posterior distribution

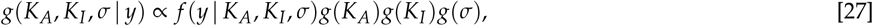

where *y* corresponds to the set of fold-change measurements. In the above model, we have assumed that the priors for *K*_*A*_ and *K*_*I*_ are independent. It is possible that it is more appropriate to assume that they are dependent and that a single prior distribution captures both parameters, *g*(*K*_*A*_, *K*_*I*_). However, assigning this prior is more difficult and requires strong knowledge *a priori* about the relationship between them. Therefore, we continue under the assumption that the priors are independent.

The generic posterior given in Eq. (27) can be extended to evaluate the second hypothesis in which Δ*ε*_*AI*_is also modified,

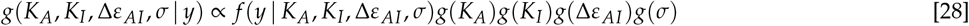

where we have included Δ*ε* _*AI*_as an estimated parameter and assigned a prior distribution.

As we have assumed that the fold-change measurements across replicates are independent and normally distributed, the likelihoods for each hypothesis can be written as

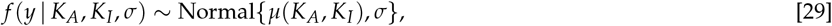

for the first hypothesis and

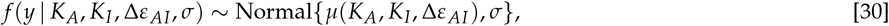

for the second. Here, we have assigned *μ*(…) as the mean of the normal distribution as a function of the parameters defined by our fold-change equation, Eq. (7).

With a likelihood distribution in hand, we now turn toward assigning functional forms to each prior distribution. As we have used in the previous sections [Sec. 2 and Sec. 3], we can assign a half-normal prior for *σ* with a standard deviation of 0.1, namely,

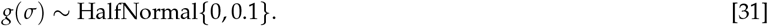

It is important to note that the inducer dissociation constants *K*_*A*_ and *K*_*I*_ are scale invariant, meaning that a change from 0.1 *μ*M to 1 *μ*M yields a decrease in affinity equal to a change from 10 *μ*M to 100 *μ*M. As such, it is better to sample the dissociation constants on a logarithmic scale. We can assign a log normal prior for each dissociation constant as

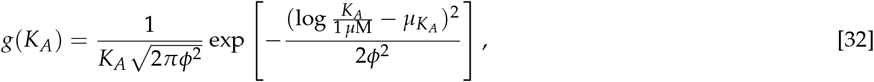

or with the short-hand notion of

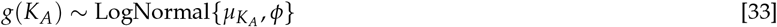

For *K*_*A*_, we assigned a mean *μK*_*A*_ = 2 and a standard deviation *ϕ* = 2. For *K*_*I*_, we chose a mean of *μK*_*I*_ = 0 and *ϕ* = 2, capturing our prior knowledge that *K*_*A*_ > *K*_*I*_ for the wild-type LacI. While the prior distributions are centered differently, they both show extensive overlap, permitting mutations in which *K*_*A*_ < *K*_*I*_. For Δ*ε* _*AI*_, we assign a normal distribution of the prior centered at 0 with a standard deviation of 5 *k*_*B*_*T*,

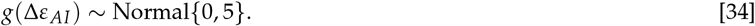

This permits values of Δ*ε* _*AI*_that are above or below zero, meaning that the inactive state of the repressor can be either more or less energetically favorable to the active state. A standard deviation of 5 *k*_*B*_*T* permits a wide range of energies with +5 *k*_*B*_*T* and −5 *k*_*B*_*T* corresponding to ≈ 99.5% and ≈ 0.5% of the repressors being active in the absence of inducer, respectively.

#### Prior predictive checks

To ensure that these choices of prior distributions are appropriate, we performed prior predictive checks for each hypothesis as previously described in Section 2. We drew 1000 values from the prior distributions shown in Fig. S13(A) for *K*_*A*_, *K*_*I*_, and Δ*ε* _*AI*_. Using the draws from the *K*_*A*_, and *K*_*I*_ priors alone, we generated datasets of ≈ 70 measurements. The percentiles of the fold-change values drawn for the 1000 simulations is shown in the top panel of Fig. S13(B).

It can be seen that in the absence of inducer, the fold-change values are close to zero and are with distributed about the leakiness value due to *σ*. This is in contrast to the data sets generated when Δ*ε* _*AI*_is permitted to vary along with *K*_*A*_ and *K*_*I*_. In the bottom panel of Fig. S13(B), the fold-change when *c* = 0 can extend above 1.0 which is possible only when Δ*ε* _*AI*_ is included, which sets what fraction of the repressors is active. Under both hypotheses, the 99^th^ percentile of the fold-change extends to just above 1 or just below 0, which matches our intuition of how the data should behave. Given these results, we are satisfied with these choices of priors and continue onto the next level of calibration of our model.

**Fig. S13.**
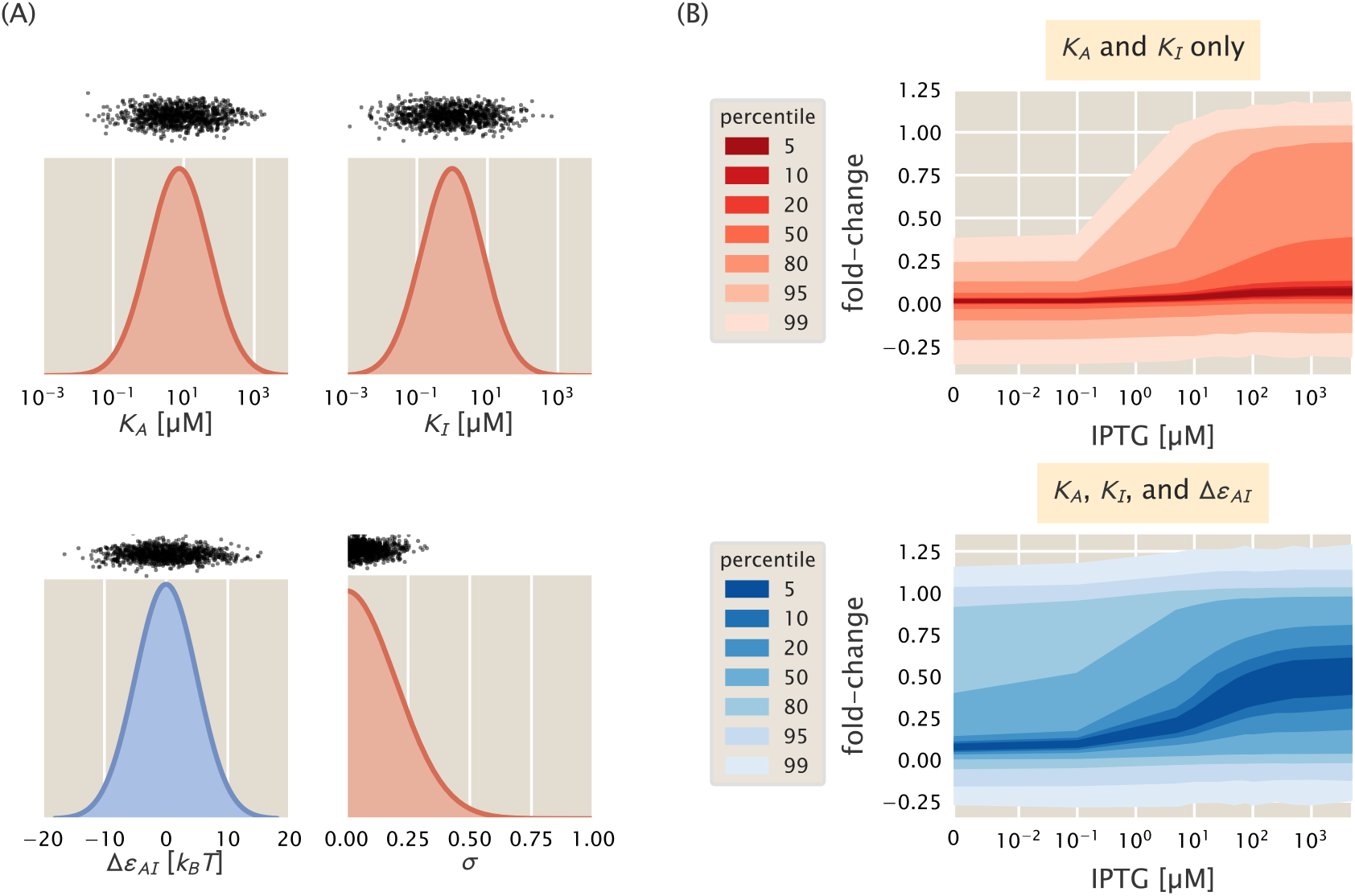
Prior predictive checks for two hypotheses of inducer binding domain mutants. (A) Probability density functions for *K*_*A*_, *K*_*I*_, Δ*ε _AI_*, and *σ*. Black points correspond to draws from the distributions used for prior predictive checks. (B) Percentiles of the simulated data sets using draws from the *K*_*A*_ and *K*_*I*_ distributions only (top, red bands) and using draws from *K*_*A*_, *K*_*I*_, and Δ*ε _AI_* (bottom, blue bands).

#### Simulation Based Calibration

With an appropriate choice of priors, we turn to simulation based calibration to root out any pathologies lurking in the model itself or the implementation through MCMC. For each parameter under each model, we compute the *z*-score and shrinkage of each inference, shown in Fig. S14. Under the first hypothesis in which *K*_*A*_ and *K*_*I*_ are the only perturbed parameters [Fig. S14(A)], we see all parameters have *z*-scores clustered around 0, indicating that the value of the ground-truth is being accurately estimated through the inference. While the shrinkage for *σ* is close to 1 (indicating the prior is being informed by the data), the shrinkage for *K*_*A*_ and *K*_*I*_ is heavily tailed with some values approaching zero. This is true for both statistical models, indicating that for some values of *K*_*A*_ and *K*_*I*_, the parameters are difficult to pin down with high certainty. In the application of these models to data, this will be revealed as large credible regions in the reported parameters. Under the second hypothesis in which all allosteric parameters are allowed to change, we see moderate shrinkage for Δ*ε* _*AI*_[purple points in Fig. S14(B)] with the minimum shrinkage being around 0.5. The samples resulting in low shrinkage correspond to values of Δ*ε* _*AI*_that are highly positive or highly negative, in which small changes in the active fraction of repressors cannot be accurately measured through our model. However, the median shrinkage for Δ*ε* _*AI*_is approximately 0.92, meaning that the the data highly informed the prior distributions for the majority of the inferences. The rank distributions for all parameters under each model appear to be highly uniform, indicating that both statistical models are computationally tractable.

With knowledge of the caveats of estimating *K*_*A*_ and *K*_*I*_ for both models, we proceed with our analysis and examine how accurately these models can capture the phenomenology of the data.

#### Posterior Predictive Checks

With a properly calibrated statistical model for each hypothesis, we now apply it to a representative dataset. While each model was applied to each inducer binding domain mutant, we only show the application to the mutant Q294K with 260 repressors per cell paired with the native *lac* operator O2.

**Fig. S14.**
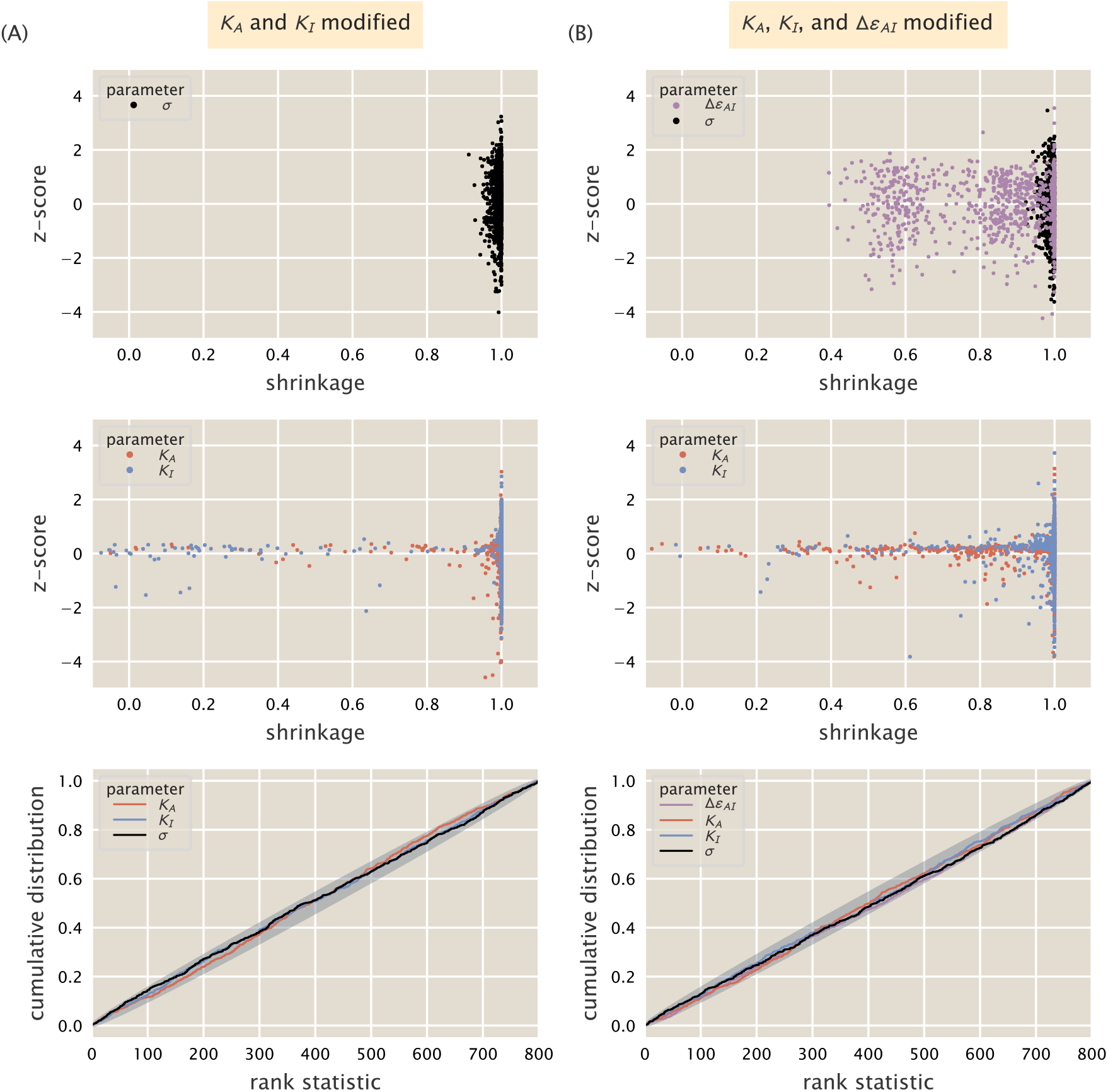
Simulation based calibration of statistical models for inducer binding domain mutants. (A) Sensitivity statistics and rank distribution for a statistical model in which *K*_*A*_ and *K*_*I*_ are the only parameters permitted to vary. (B) Sensitivity statistics and rank distribution for a model in which all allosteric parameters *K*_*A*_, *K*_*I*_, and Δ*ε _AI_* are allowed to be modified by the mutation. Gray envelope in the bottom plots correspond to the 99^th^ percentile of variation expected from a true uniform distribution.

The results from applying the statistical model in which only *K*_*A*_ and *K*_*I*_ can change is shown in Fig. S15. The joint and marginal distributions for each parameter [Fig. S15(A)] reveal a strong correlation between *K*_*A*_ and *K*_*I*_ whereas all other parameters are symmetric and independent. While the joint and marginal distributions look well behaved, the percentiles of the posterior predictive checks [Fig. S15(B)] are more suspect. While all data falls within the 95^th^ percentile, the overall trend of the data is not well predicted. Furthermore, the percentiles expand far below zero, indicating that the sampling of *σ* is compensating for the leakiness in the data being larger than it should be if only *K*_*A*_ and *K*_*I*_ were the changing parameters.

We see significant improvement when Δ*ε* _*AI*_is permitted to vary in addition to *K*_*A*_ and *K*_*I*_. Fig. S16(A) shows the joint and marginal distributions between all parameters from the MCMC sampling. We still see correlation between *K*_*A*_ and *K*_*I*_, although it is not as strong as in the case where they are the only parameters allowed to change due to the mutation. We also see that the marginal distribution for *σ* has shrunk significantly compared to the marginal distribution in Fig. S15(A). The percentiles of the posterior predictive checks, shown in Fig. S16(B) are much more in line with the experimental measurements, with the 5th percentile following the data for the entire induction profile.

In this section we have presented two hypotheses for the minimal parameter set needed to describe the inducer binding mutations, derived a statistical model for each, thoroughly calibrated its behavior, and applied it to a representative data set. The posterior predictive checks [Fig. S15 and Fig. S16] help us understand which hypothesis is more appropriate for that particular mutant. The incredibly wide percentiles and significant change in the leakiness that result from a model in which only *K*_*A*_ and *K*_*I*_ are perturbed suggests that more than those two parameters should be changing. We see significant improvement in the description of the data when Δ*ε* _*AI*_is altered, indicating that it is the more appropriate hypothesis of the two.

**Fig. S15.**
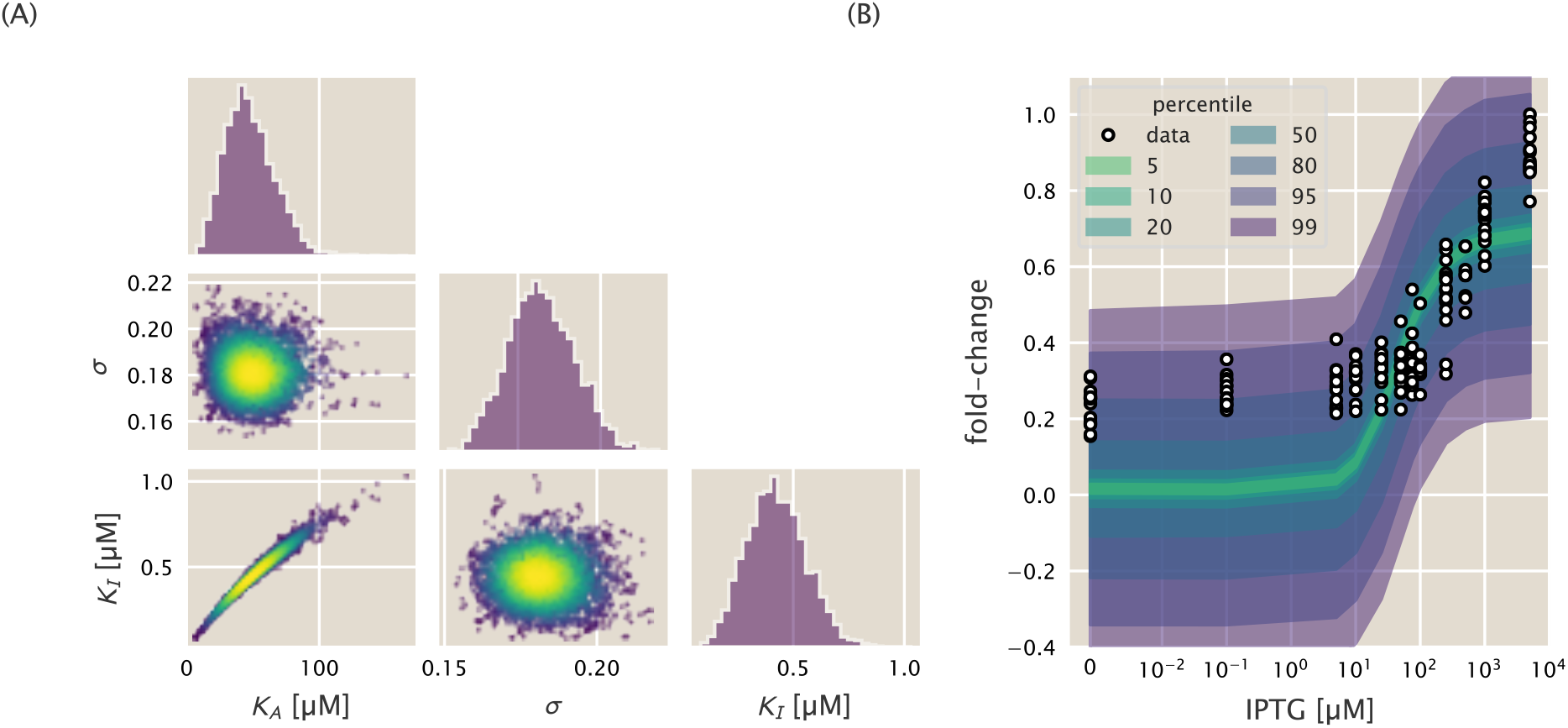
Posterior predictive checks for inducer binding domain mutants where only *K*_*A*_ and *K*_*I*_ are changed. (A) MCMC sampling output for each parameter. Joint distributions are colored by the value of the log posterior with increasing probability corresponding to transition from blue to yellow. (B) Percentiles of the data generated from the likelihood distribution for each sample of *K*_*A*_, *K*_*I*_, and *σ*. Overlaid points are the experimentally observed measurements.

**Fig. S16.**
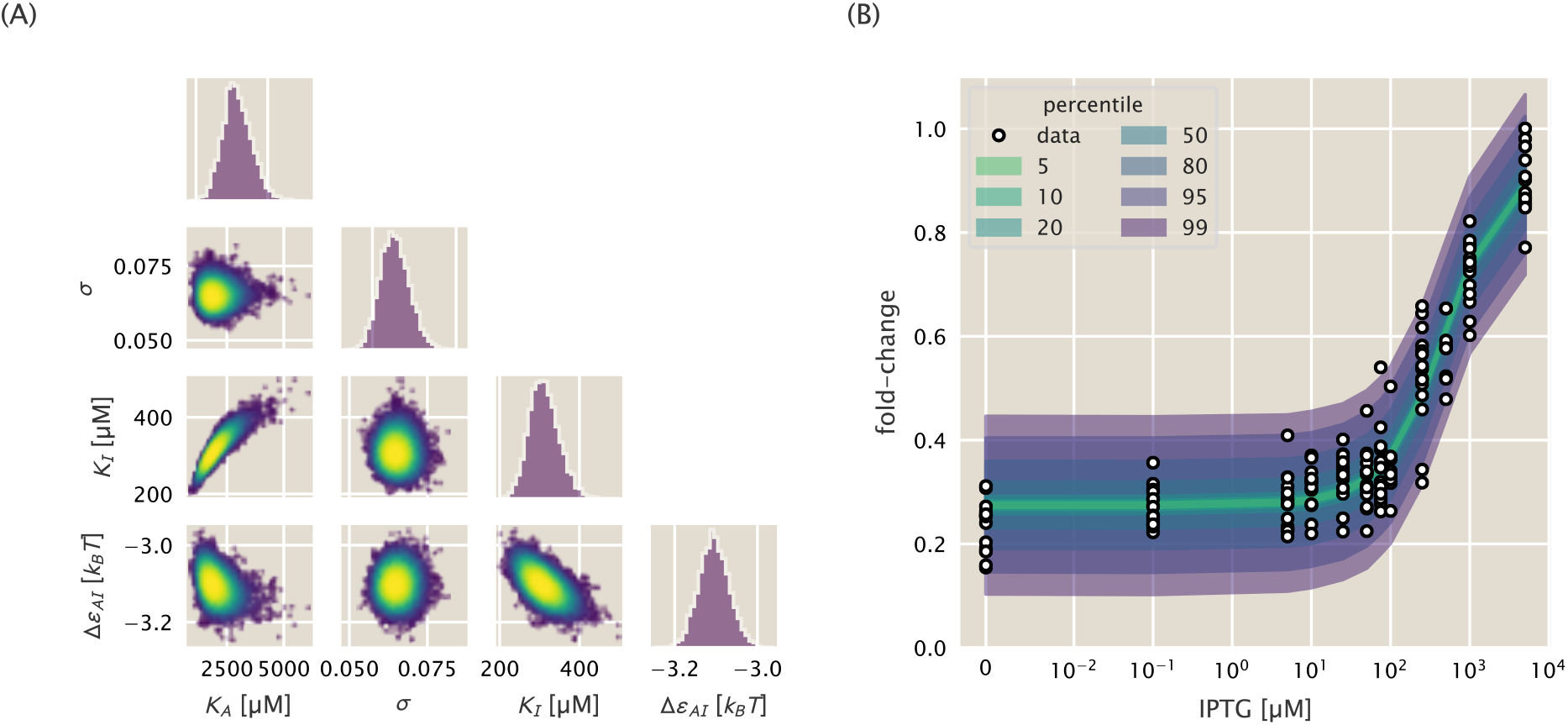
Posterior predictive checks for inducer binding domain mutants where all allosteric parameters can change. (A) MCMC sampling output for all parameters. Joint distributions are colored by the value of the log posterior with increasing probability corresponding to the transition from blue to yellow. Marginal distributions are shown adjacent to each joint distribution. (B) Percentiles of the data generated from the likelihood for each sample of *K*_*A*_, *K*_*I*_, Δ*ε _AI_*, and *σ*. The corresponding experimental data for Q294K are shown as black open-faced circles.

### 6. Additional Characterization of Inducer Binding Domain Mutants

To predict the induction profiles of the inducer binding mutants, we used only the induction profile of each mutant paired with the native O2 *lac* operator to infer the parameters. Here, we examine the influence the choice of fit strain has on the predictions of the induction profiles and Δ*F* for each mutant.

In the main text, we dismissed the hypothesis that only *K*_*A*_ and *K*_*I*_ were changing due to the mutation and based the fit to a single induction profile. In Fig. S17, the fits and predictions for each mutant paired with each operator sequence queried. Here, the rows correspond to the operator sequence of the fit strain while the columns correspond to the operator sequence of the predicted strain. The diagonals, colored in gray, show the fit induction profiles and the corresponding data. Regardless of the choice of fit strain, the predicted induction profiles of the repressor paired with the O3 operator are poor, with the leakiness in each case being significantly underestimated. We also see that fitting to O3 results in poor predictions with incredibly wide credible regions for the other two operators. In Razo-Mejia et al. 2018 (2), we also found that fitting *K*_*A*_ and *K*_*I*_ to the induction profile of O3 generally resulted in poor predictions of the other strains with comparably wide credible regions.

When Δ*ε* _*AI*_is included as a parameter, however, the predictive power is improved for all three operators, as can be seen in Fig. S18. While the credible regions are still wide when fit to the O3 operator, they are much narrower than under the first hypothesis. We emphasize that we are able to accurately predict the leakiness of nearly every strain by redetermining Δ*ε*_*AI*_ whereas the leakiness was not predicted when only *K*_*A*_ and *K*_*I*_ were considered. Thus, we conclude that all three allosteric parameters *K*_*A*_, *K*_*I*_, and Δ*ε* _*AI*_are modified for these four inducer binding domain mutations. The values of the inferred parameters are reported in Table S2.

We also examined the effect the choice of fit strain has on the predicted Δ*F*, shown in Fig. S19. We find that the predictions agree with the data regardless of the choice of fit strain. One exception is the prediction of the Q294K Δ*F* when the parameters fit to the O3 induction profile are used. As the induction profile for Q294K paired with O3 is effectively flat at a fold-change of 1, it is difficult to properly estimate the parameters of our sigmoidal function. We note all measurements of Δ*F* for Q294K are described by using either the parameters fit to either O1 or O3 induction profiles, suggesting that the choice of fit strain makes little difference.

**Fig. S17.**
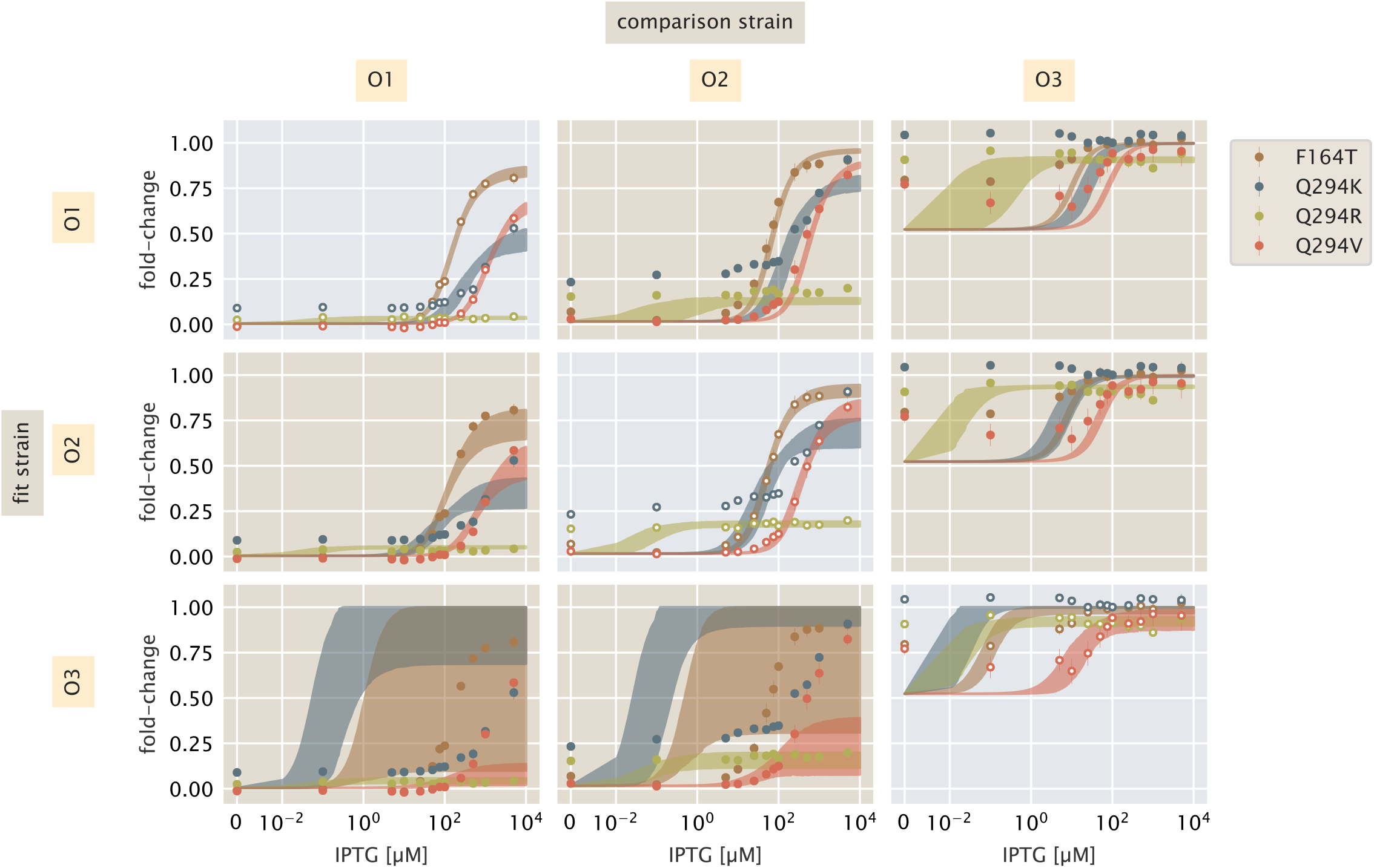
Pairwise comparison of fit strain versus predictions assuming only *K*_*A*_ and *K*_*I*_ are influenced by the mutation. Rows correspond to the operator sequence of the strain used for the parameter inference. Columns correspond to the operator sequence of the predicted strain. Colors identify the mutation. Diagonal positions (gray background) show the induction fit strain and profiles.

**Fig. S18.**
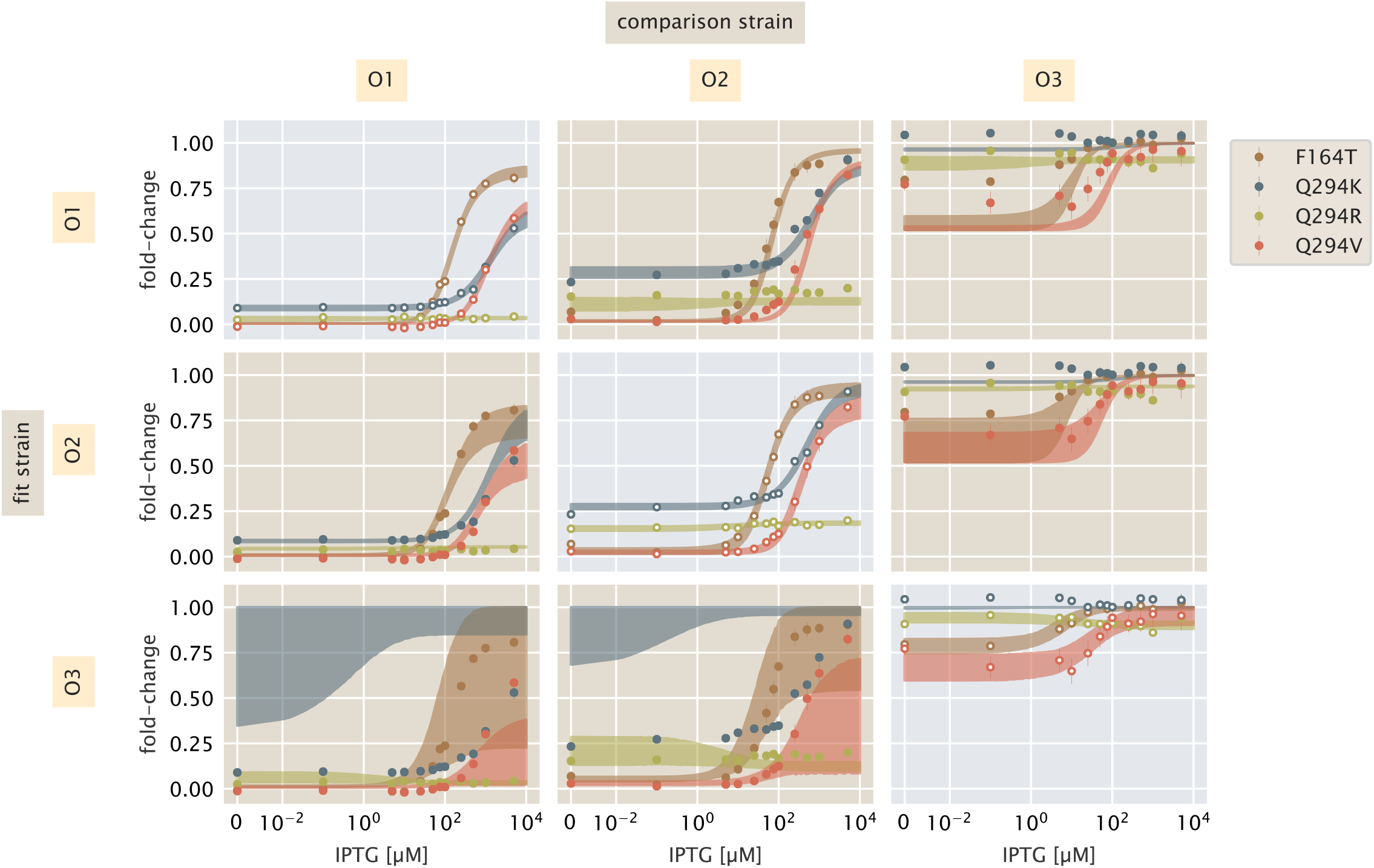
Pairwise comparison of fit strain versus predictions assuming all allosteric parameters are affected by the mutation. Rows correspond to the operator of the strain used to fit the parameters. Columns correspond to the operator of the strains whose induction profile is predicted. Mutants are identified by color. Diagonals (gray background) show the induction profiles of the strain to which the parameters were fit.

**Fig. S19.**
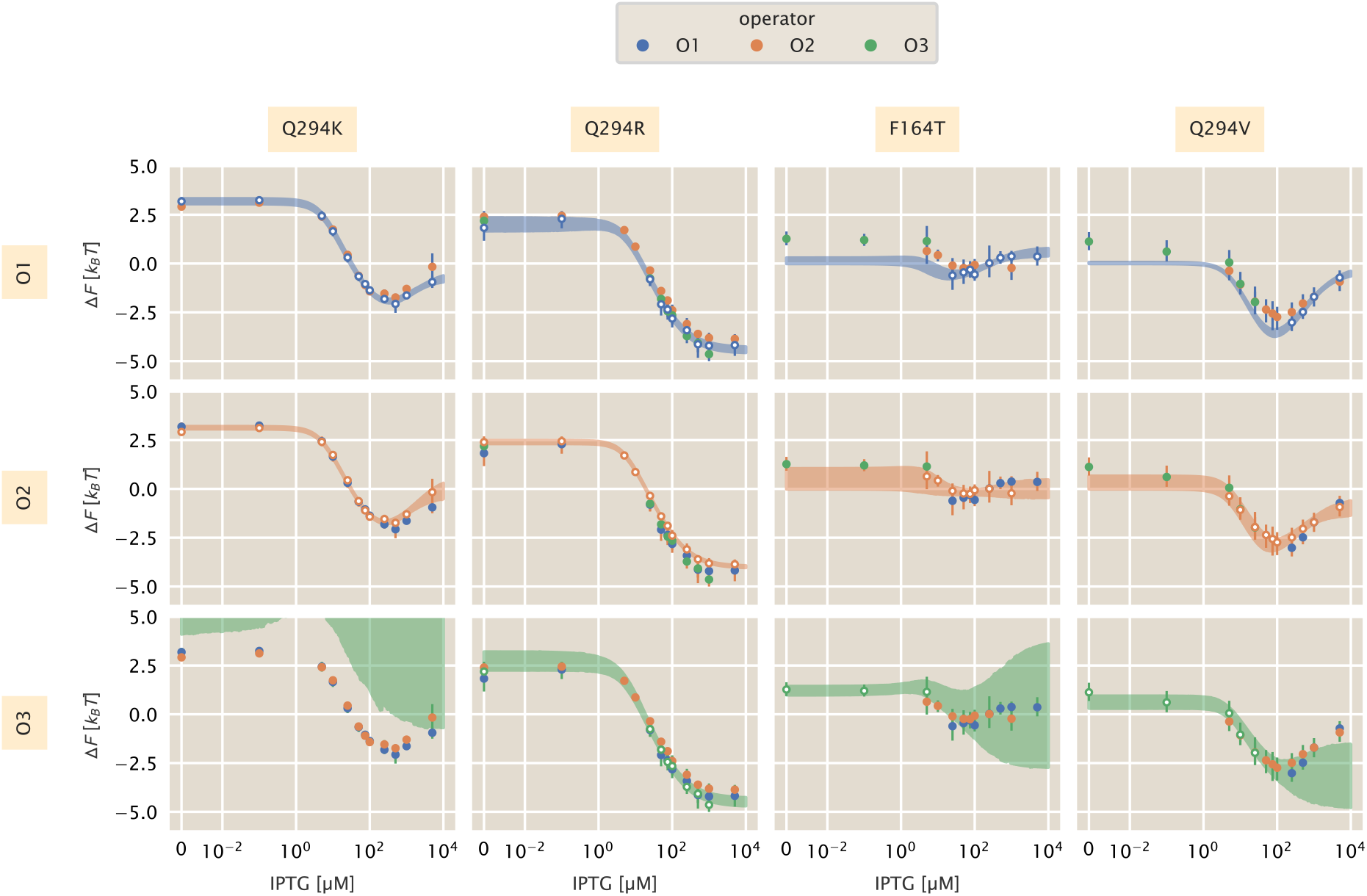
Comparison of choice of fit strain on predicted Δ*F* profiles. Rows correspond to the operator of the strain to which the parameters were fit. Columns correspond to mutations. Points are colored by their operator sequence. The data corresponding to the operator of the fit strain are shown as white-faced points.

**Table S2.** Inferred values of *K*_*A*_, *K*_*I*_, and Δ*ε*_*AI*_ for inducer binding domain mutants. Values reported are the mean of the posterior distribution with the upper and lower bounds of the 95% credible region.

| Mutant | Operator | $K_A$ [$\mu\text{M}$] | $K_I$ [$\mu\text{M}$] | $\Delta\epsilon_{AI}$ [$\text{k}_\text{B}$ T] |
| --- | --- | --- | --- | --- |
| F164T | O1 | $290^{+60}_{-56}$ | $1^{+4}_{-0.98}$ | $4^{+5}_{-3}$ |
| | O2 | $165^{+90}_{-65}$ | $3^{+6}_{-3}$ | $1^{+5}_{-2}$ |
| | O3 | $110^{+700}_{-105}$ | $7^{+5}_{-4}$ | $-0.9^{+0.4}_{-0.3}$ |
| Q294K | O1 | > 1 mM | $410^{+150}_{-100}$ | $-3.2^{+0.1}_{-0.1}$ |
| | O2 | > 1 mM | $310^{+70}_{-60}$ | $-3.11^{+0.07}_{-0.07}$ |
| | O3 | $10^{+200}_{-10}$ | $1^{+9}_{-1}$ | $-7^{+3}_{-5}$ |
| Q294R | O1 | $3^{+27}_{-3}$ | $2^{+20}_{-2}$ | $-1.9^{+0.4}_{-0.3}$ |
| | O2 | $9^{+20}_{-9}$ | $8^{+20}_{-8}$ | $-2.32^{+0.01}_{-0.09}$ |
| | O3 | $6^{+24}_{-6}$ | $9^{+30}_{-9}$ | $-2.6^{+0.4}_{-0.5}$ |
| Q294V | O1 | > 1 mM | $3^{+13}_{-3}$ | $6^{+4}_{-4}$ |
| | O2 | $650^{+450}_{-250}$ | $8^{+8}_{-8}$ | $3^{+6}_{-3}$ |
| | O3 | $100^{+400}_{-90}$ | $22^{+33}_{-18}$ | $0.1^{+0.8}_{-0.6}$ |

### 7. Parameter Estimation Using All Induction Profiles

In the main text and Sec. 4 and 6 of this supplementary text, we have laid out our strategy for inferring the the various parameters of our model to a single induction profile and using the resulting values to predict the free energy and induction profiles of other strains. In this section, we estimate the parameters using all induction profiles of a single mutant and using the estimated values to predict the free energy profiles.

The inferred DNA binding energies considering induction profiles of all repressor copy numbers for the three DNA binding mutants are reported in Tab. S3. These parameters are close to those reported in Tab. S1 for each repressor copy number with Q21A showing the largest differences. The resulting induction profiles and predicted change in free energy for these mutants can be seen in Fig. **??**. Overall, the induction profiles match the data to an appreciable agree. We acknowledge that even when using *all* repressor copy numbers, the fit to Q21A remains imperfect. However we contend that this disagreement is comparable to that observed in (2) which described the induction profile of the wild-type repressor. We find that the predicted change in free energy [bottom row in Fig. **??**(B)] narrows compared to that in Fig. S12 and Fig. 3 of the main text, confirming that considering all induction profiles improves our inference of the most-likely DNA binding energy. There appears to be a very slight trend in the Δ*F* for Q21A at higher inducer concentrations, though the overall change in free energy from 0 to 5000 *μ*M IPTG is small.

We also estimated the allosteric parameters (*K*_*A*_, *K*_*I*_, and Δ*ε* _*AI*_) for all inducer binding domain mutations using the induction profiles of all three operator sequences. The values, reported in Tab. S4 are very similar to those estimated from a single induction profile (Tab. S2). We note that for Q294R, it is difficult to properly estimate the values for *K*_*A*_ and *K*_*I*_ as the observed induction profile is approximately flat. The induction profiles and predicted change in free energy for each inducer binding mutant is shown in Fig. S21. We see notable improvement in the agreement between the induction profiles and the observed data, indicating that considering all data significantly shrinks the uncertainty of each parameter. The predicted change in free energy is also improved compared to that shown in Fig. S19. We emphasize that the observed free energy difference for each point assumes no knowledge of the underlying parameters and comes directly from measurements. The remarkable agreement between the predicted free energy and the observations illustrates that redetermining the allosteric parameters is sufficient to describe how the free energy changes as a result of the mutation.

**Table S3.**
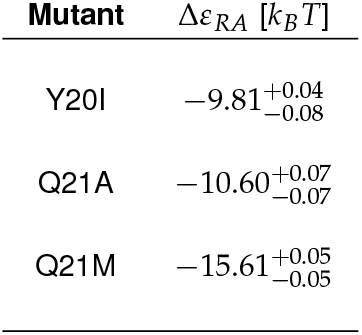
Estimated DNA binding energies for each DNA binding domain mutant using all repressor copy numbers

**Fig. S20.**
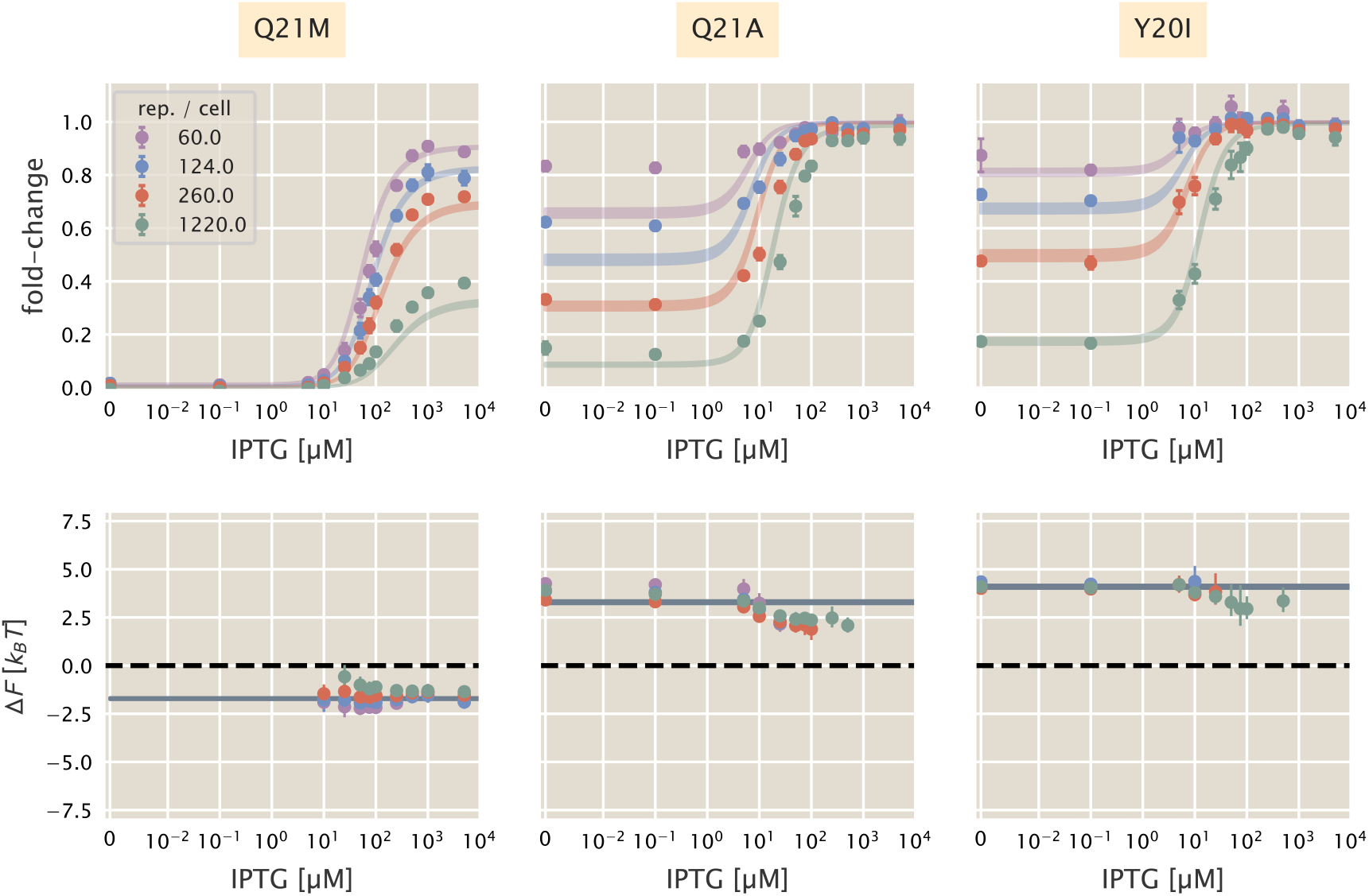
Induction profiles and predicted change in free energy using parameters estimated from the complete data sets. Top row shows fold-change measurements (points) as mean and standard error with ten to fifteen biological replicates. Shaded lines correspond to the 95% credible regions of the induction profiles using the estimated values of the DNA binding energies reported in Tab. S3. Bottom row shows the 95% credible regions of the predicted change in free energy (shaded lines) along with the inferred free energy of data shown in the top row. In all plots, the inducer concentration is shown on a symmetric log scale with linear scaling between 0 and 10^−2^ *μ*M and log scaling elsewhere.

**Table S4.**
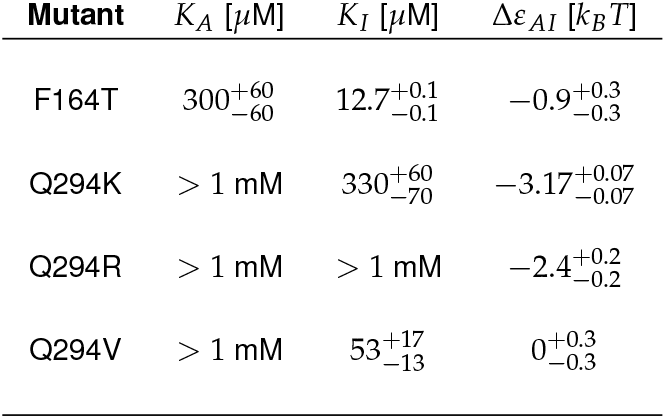
Estimated values for *K*_*A*_, *K*_*I*_, and Δ*ε*_*AI*_ for inducer binding domain mutations using induction profiles of all operator sequences.

**Fig. S21.**
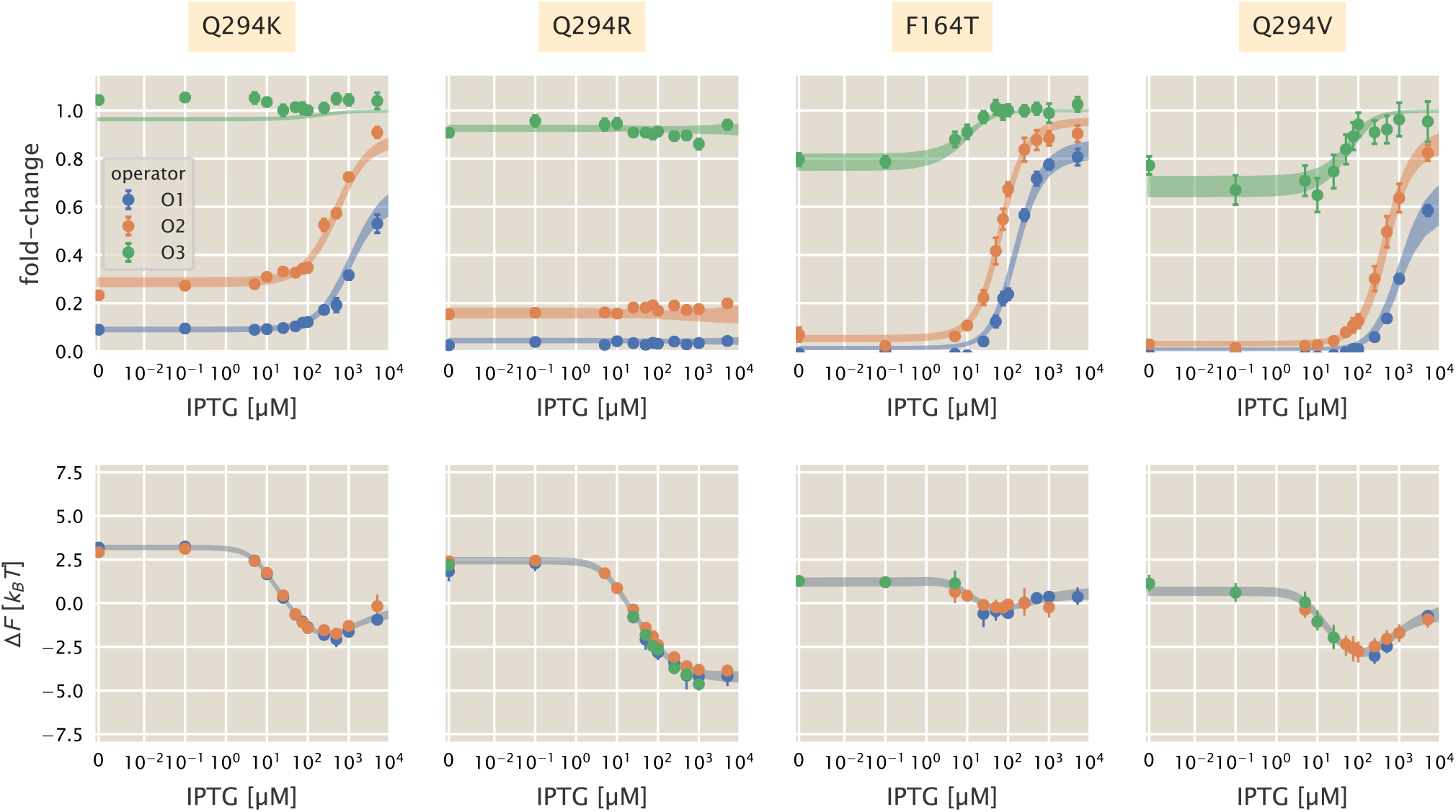
Induction profiles and predicted change in free energy using parameters estimated from the complete data sets for inducer binding domain mutnats. Top row shows fold-change measurements (points) as mean and standard error with ten to fifteen biological replicates. Shaded lines correspond to the 95% credible regions of the induction profiles using the estimated values of the allosteric parameters reported in Tab. S4. Bottom row shows the 95% credible regions of the predicted change in free energy (shaded lines) along with the inferred free energy of data shown in the top row. In all plots, the inducer concentration is shown on a symmetric log scale with linear scaling between 0 and 10^−2^ *μ*M and log scaling elsewhere.

### Strain and Oligonucleotide Information

**Table S5.**
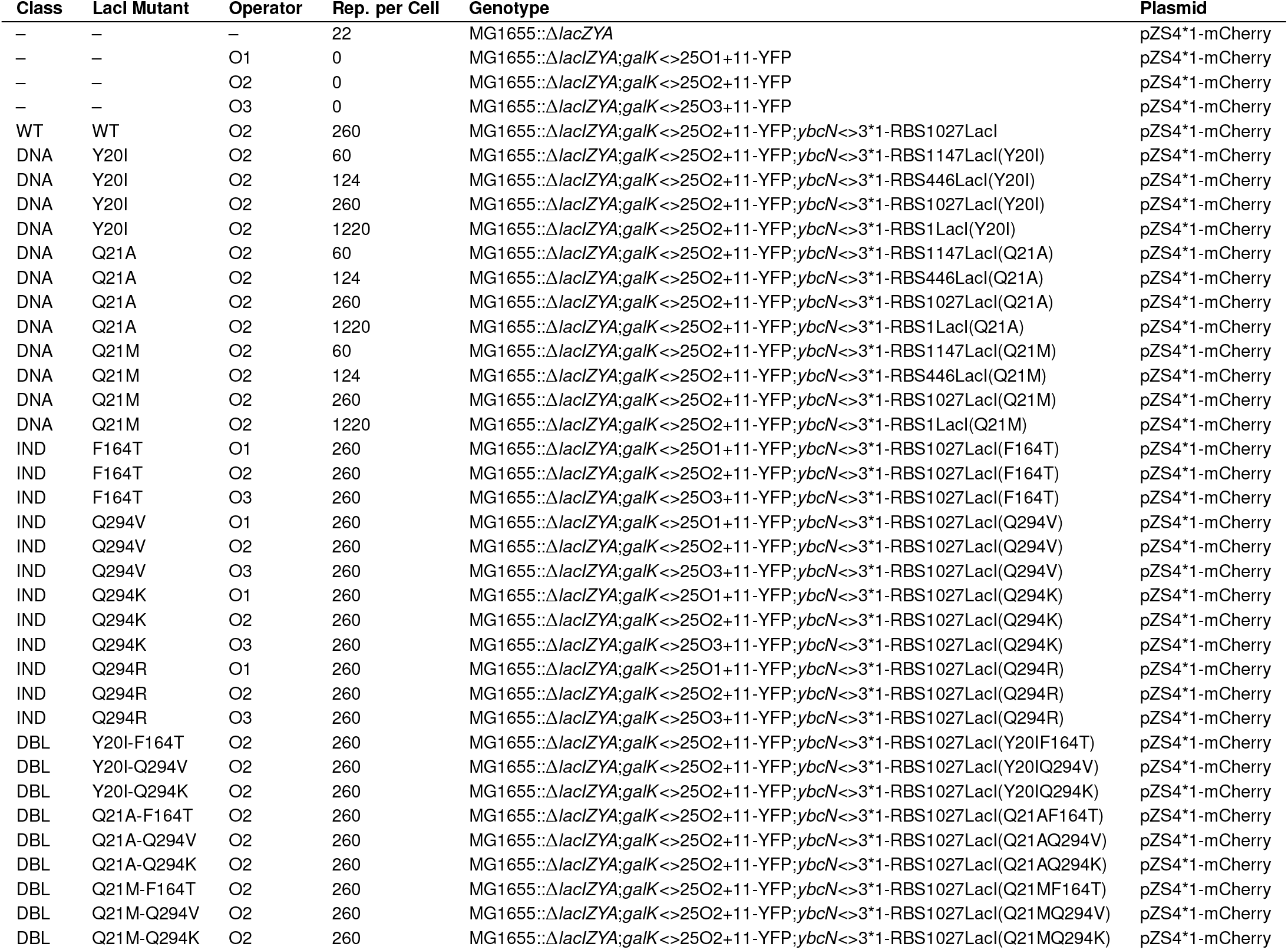
*Escherichia coli* strains used in this work

**Table S6.**
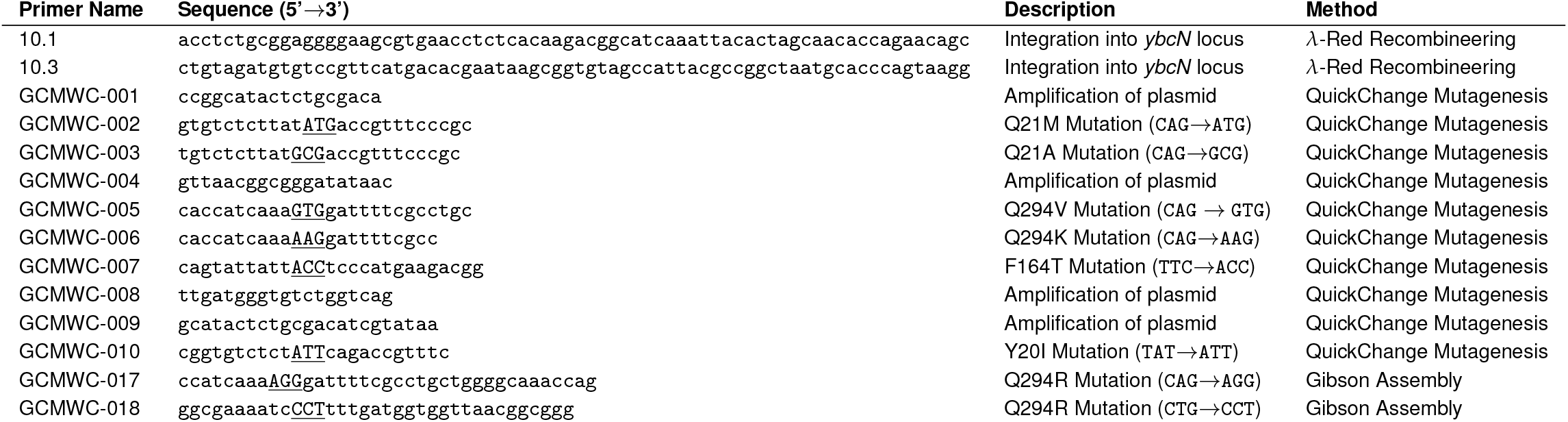
Oligonucleotides used for mutant generation.

## Notes

The authors declare no conflict of interest.

http://www.rpgroup.caltech.edu/mwc_mutants/

https://github.com/rpgroup-pboc/mwc_mutants

https://data.caltech.edu/records/1241

